# Structure, interdomain dynamics and pH-dependent autoactivation of pro-rhodesain, the main lysosomal cysteine protease from African trypanosomes

**DOI:** 10.1101/2020.11.10.363747

**Authors:** Patrick Johé, Elmar Jaenicke, Hannes Neuweiler, Tanja Schirmeister, Christian Kersten, Ute A. Hellmich

## Abstract

Rhodesain is the lysosomal cathepsin L-like cysteine protease of *T. brucei rhodesiense*, the causative agent of Human African Trypanosomiasis. The enzyme is essential for the proliferation and pathogenicity of the parasite as well as its ability to overcome the blood-brain barrier of the host. Lysosomal cathepsins are expressed as zymogens with an inactivating pro-domain that is cleaved under acidic conditions. A structure of the uncleaved maturation intermediate from a trypanosomal cathepsin L-like protease is currently not available. We thus established the heterologous expression of *T. brucei rhodesiense* pro-rhodesain in *E. coli* and determined its crystal structure. The trypanosomal pro-domain differs from non-parasitic pro-cathepsins by a unique, extended α-helix that blocks the active site and whose interactions resemble that of the antiprotozoal inhibitor K11777. Interdomain dynamics between pro- and core protease domain as observed by photoinduced electron transfer fluorescence correlation spectroscopy increase at low pH, where pro-rhodesain also undergoes autocleavage. Using the crystal structure, molecular dynamics simulations and mutagenesis, we identify a conserved interdomain salt bridge that prevents premature intramolecular cleavage at higher pH values and may thus present a control switch for the observed pH-sensitivity of pro-enzyme cleavage in (trypanosomal) CathL-like proteases.

## Introduction

Human African Trypanosomiasis (HAT) or African Sleeping Sickness is a so-called neglected tropical disease (NTD) as classified by the World Health Organization (www.who.int). HAT is caused by two subspecies of African trypanosomes, *Trypanosoma brucei gambiense* and *Trypanosoma brucei rhodesiense*.^[1]^ These unicellular, protozoan parasites are transmitted to the host via the bite of a Tse-Tse fly. The disease progresses in two stages. In the early hemolymphatic stage, the parasites are present in the blood and in lymphatic systems leading to unspecific symptoms like fever and headache. In the later meningoencephalitic stage, the parasites cross the blood-brain barrier and enter the central nervous system. At this point, patients suffer severe neurological symptoms, deregulation of sleep-wake cycles, coma and ultimately death.^[2]^ The few available treatment options often have severe side effects. For instance, the only approved drug for the treatment of a stage 2 *T. b. rhodesiense* infection is melarsoprol, an arsenic-based drug with severe side effects including a 5% lethality rate.^[3]^

*T. brucei* spp. parasites express two main lysosomal cysteine proteases that belong to the papain family, the cathepsin B-like member *T. brucei* Cathepsin B (TbCathB) and the cathepsin L-like protease rhodesain (also called TbCathL, brucipain or trypanopain). Rhodesain plays an important role in the progression into the second disease stage as it is involved in the crossing of parasites into the central nervous system via the blood-brain barrier (BBB) of the host.^[4,5]^ The expression of TbCathB mRNA varies strongly during the life cycle of the parasite, while rhodesain is constitutively expressed.^[6]^ In agreement with an important role in parasite survival, proliferation and pathogenicity, RNAi-based rhodesain knock-down or inhibition of rhodesain but not TbCathB led to diminished parasitic growth and an increased sensibility to lytic factors in human serum.^[7,8]^ Rhodesain is thus generally assumed to be the main lysosomal cysteine protease in trypanosomes and presents a promising anti-trypanosomal drug target. In general, proteases are attractive drug targets because the affinities of inhibitors towards active site residues can be tuned and potential inhibitors can easily be identified in high-throughput *in vitro* assay screenings.^[9]^ Nevertheless, structural similarities to host proteins can lead to problems with off-target effects. Rhodesain shares 46% (60%) sequence identity (similarity) with human cathepsin L (hsCathL). The irreversible, vinylsulfone-based rhodesain inhibitor K11777 (*N*-methylpiperazine-urea-Phe-homophenylalanine-vinylsulfone-benzene), was shown to prevent *T. brucei* from overcoming a barrier formed by brain microvascular endothelial cells in an *in vitro* BBB model.^[4,5]^ Encouragingly, while K11777 was indeed observed to not be biochemically selective, it was non-toxic to mammals, efficient against related protozoans and could cure mice from an *Trypanosoma cruzi* infection^[10–12]^.

In the cell, cathepsins are expressed as inactive zymogens where the ~215 amino acid papain-fold core protease domain is preceded by a ~20 amino acid signal peptide required for translation of the protein into the endoplasmatic reticulum, and a ~100 amino acid pro-domain required for correct folding and lysosomal targeting of the protease.^[13–17]^ In addition, the pro-domain interacts with the active site of the protease and keeps it in an autoinhibited state until it has been successfully trafficked to the acidic lysosome lumen. Here, the pro-domain is removed by proteolytic cleavage, which may occur intra- or intermolecularly.^[18–21]^ The cleavage site is located in the linker between the pro-domain and the core protease domain.^[22,23]^ While the rhodesain core protease domain has been crystallized in complex with different inhibitors including K11777 (PDB: 2P7U) ^[23–26]^, a structure of the zymogen as an important intermediate of the parasitic protease maturation process is currently not available. In addition, the molecular details of the pH-dependent cleavage process remain unclear.

Here, we present the 2.8 Å crystal structure of the active site C150A mutant of the *T. brucei rhodesiense* cysteine protease rhodesain in complex with its pro-domain under acidic conditions. The rhodesain pro-domain shares many structural features with other CathL pro-domains, but is unique in the presence of an extended α-helix in its C-terminal end. This helix interacts with the active site in an orientation similar to that of the inhibitor K11777. Using photoinduced electron transfer fluorescence correlation spectroscopy (PET-FCS), we investigated the dynamics of the pro-enzyme and found that low pH leads to enhanced dynamics in the “blocking peptide”, the region of the pro-domain covering the protease active site.^[27]^ We further found evidence that pro-rhodesain can be processed both intra- and intermolecularly in a pH-dependent manner and identified a highly conserved interdomain salt-bridge that can prevent premature intramolecular cleavage at high pH and may thus act as a “delay switch” for pro-cathepsin autocleavage.

## Results

### Heterologous expression and purification of pro-rhodesain

Cathepsin-like cysteine proteases are expressed in an inactive pro-form. pH-dependent cleavage of the pro-domain from the globular core protease domain is an important step in the protease maturation process contingent on the correct localization of the protein to the lysosome. The cathepsin L-like protease rhodesain from African trypanosomes is required for the progression of African Sleeping Sickness. For a better understanding of pro-rhodesain inhibition and activation, we aimed to determine the crystal structure of *T. brucei rhodesiense* pro-rhodesain. Typically, *P. pastoris* is used as a heterologous expression host for the production of parasitic cathepsin proteases.^[22,25]^ While rhodesain expression in *P. pastoris* relies on the secretion of the folded protein, all currently available expression and purification protocols for rhodesain from *E. coli* are based on its expression in inclusion bodies and subsequent protein refolding.^[28]^ To obtain the folded pro-enzyme for structural studies, we found both production routes to be ineffective. We thus optimized the purification of mature and pro-rhodesain from *E. coli* in a manner that circumvents secretion and refolding (see Fig. S1–S4 and extended Materials and Methods for details).

Purified mature rhodesain from *P. pastoris* and *E. coli* display similar structural and functional properties as elucidated by size exclusion chromatography, circular dichroism spectroscopy (Fig, 1A, B) and a fluorescent peptide-cleavage assay using the fluorescent peptide Z-Phe-Arg-AMC (Z-phenylalanine-arginine-7-amido-4-methylcoumarin) to probe rhodesain activity (see extended Materials and Methods).^[9]^ *E. coli* can thus be used as a reliable source for the production of active rhodesain for optimal flexibility in biophysical studies. To prevent auto-cleavage and to stabilize the pro-enzyme, we introduced a mutation into the active site of the protein, C150A. Pro-rhodesain C150A displays the expected increase in molecular weight during size exclusion chromatography as well as an additional small increase in α-helical content in CD spectroscopy measurements (Fig. 1C).

**Fig. 1:**
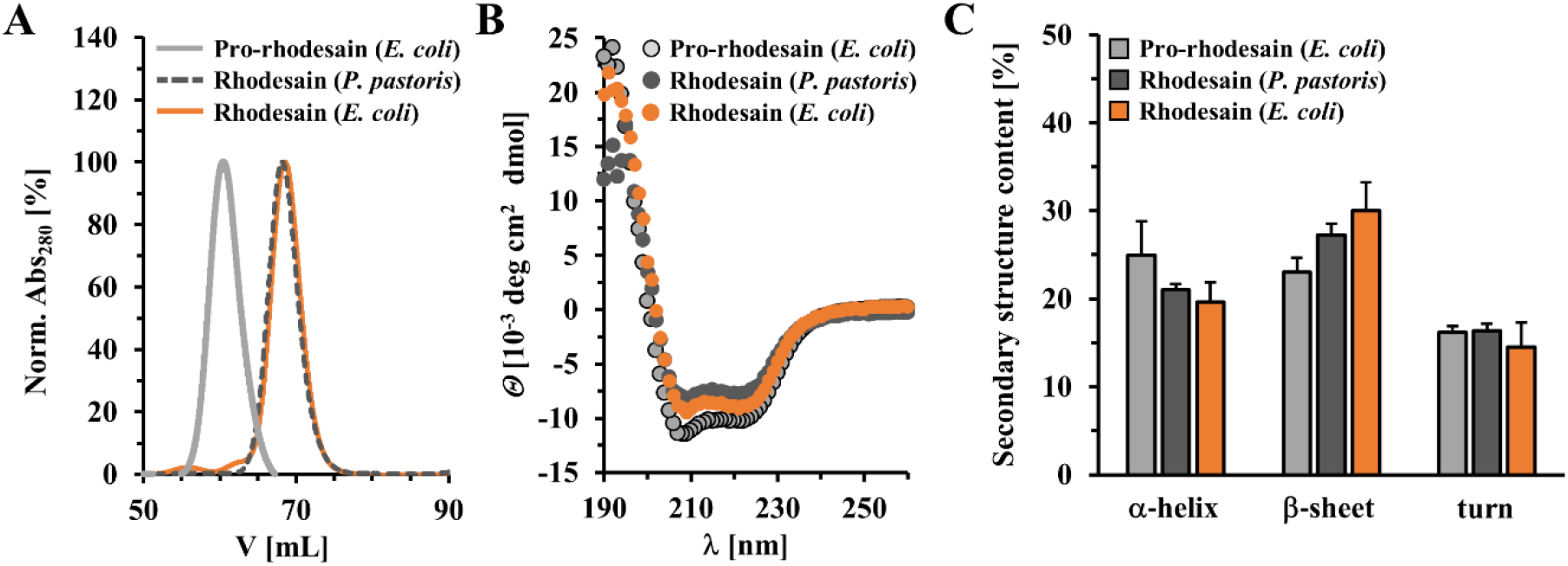
Comparison of heterologously expressed rhodesain purified from *P. pastoris* or *E. coli*. A) Size exclusion chromatography of mature rhodesain from *P. pastoris* (dashed-lines, dark grey) or *E. coli* (orange) and pro-rhodesain C150A from *E. coli* (light grey). B) Circular dichroism spectroscopy of mature rhodesain (*P. pastoris*, dark grey; *E. coli*, orange) as well as pro-rhodesain C150A (light grey). C) Secondary structure analysis of CD spectra shown in (B) using the BeStSel server (http://bestsel.elte.hu/index.php).^[29]^

### T. b. rhodesiense *pro-rhodesain crystal structure*

We obtained a 2.8 Å crystal structure for *T. b. rhodesiense* pro-rhodesain at acidic pH (Fig. 2, Table 1, Fig. S5). The pro-protease structure contains two parts, the N-terminal pro-domain (residues E38 to T123 according to uniprot numbering, ID:Q95PM0) and the C-terminal core protease domain (residues A126 to P342). The first 18 amino acids belonging to the pro-domain are not resolved. The first amino acid modeled into the electron density is therefore E38. A short loop between the first two helices of the pro-domain (residues 77/78) and residues G124/R125 at the C-terminus of the pro-domain are also not resolved. High disorder, in agreement with the crystallographic B factors in this region (Fig. 2C, D) coincides with the expected intermolecular cleavage site of pro-rhodesain at position T123 or R125.^[22,23]^ The entire C-terminus of the core protease up to residue P342 as well as the remaining residues of a TEV cleavage site (ENLYF) which was used to remove the C-terminal purification tag, could be reliably placed in the electron density. The remainder of the TEV cleavage site mediates crystal contacts to the neighboring protein’s pro domain and may have thus aided crystallization of the construct but does not interfere with the interaction of the pro-domain with its corresponding catalytic domain.

**Fig. 2:**
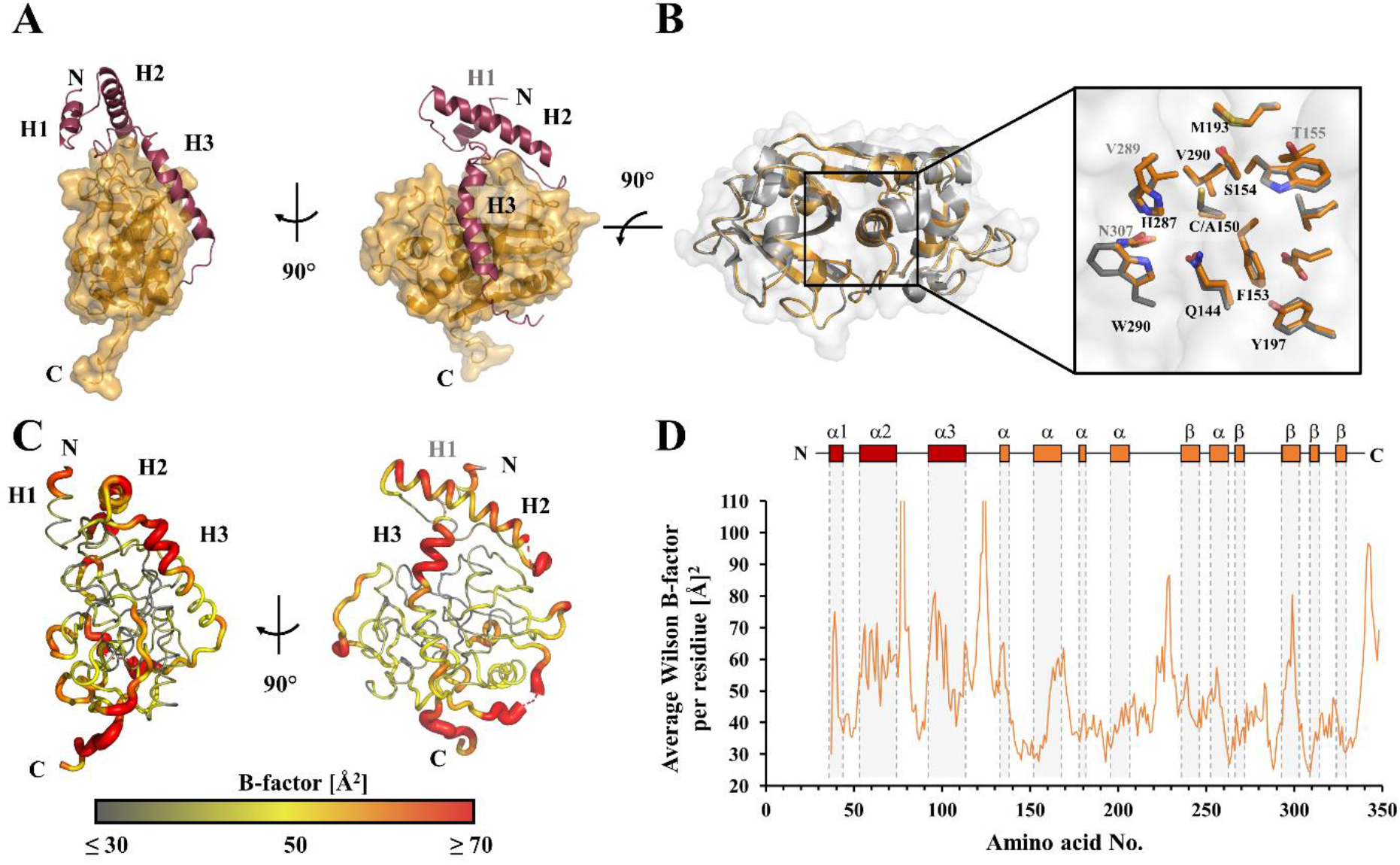
Crystal structure of *T. brucei rhodesiense* pro-rhodesain. A) 2.8 Å crystal structure of pro-rhodesain consisting of the pro-domain (red) composed of three α-helices, H1, H2 and H3 and the core protease (gold). B) Comparison of pro-rhodesain (gold) and mature rhodesain (grey, PDB: 2P7U) ^[25]^ including a close-up of the active site. Side chains within a distance of 8 Å around the active site cysteine or the C150A mutation are shown as sticks. C, D) Average B factors per residue of the pro-rhodesain structure. The low values (≤ 30 Å^2^) in the catalytic domain indicate that it is a mostly rigid protein with the exception of some flexible loop regions, thus resulting in an overall average B factor of 44.2 Å^2^. In the pro-domain, the average B factor is 55.6 Å indicating higher flexibility of the pro-peptide compared to the core protease domain. The residues with the highest B reside in the helix covering the active site cleft (H3) and the unstructured peptide chain connecting the pre- and the core protease domains.

**Table 1:** Crystallographic data collection and refinement statistics for *T. brucei rhodesiense* pro-rhodesain C150A. Numbers in parentheses characterize the highest resolution shell.

| PDB entry | 7AVM |
| --- | --- |
| **Data collection** |  |
| Wavelength | 1.54179 |
| Space group | H 3 |
| Cell parameters |  |
| a; b; c | 123.19 Å; 123.19 Å; 53.91 Å |
| $\alpha$; $\beta$; $\gamma$ | 90.00°; 90.00°; 120° |
| Resolution range [Å] | 37.92 - 2.8 (2.9-2.8) |
| Unique reflections | 7512 (729) |
| Multiplicity | 20.6 (19.8) |
| Overall completeness [%] | 97.23 (92.73) |
| $\langle I/\sigma(I) \rangle$ | 3.63 (at 2.81 Å) |
| **Refinement** |  |
| R<sub>work</sub>, R<sub>free</sub> | 0.255, 0.264 |
| Average B-Factor [Å<sup>2</sup>] | 41.2 |
| Protein residues | 308 |
| No. of solvent molecules | 38 |
| Ligand atoms | 6 |
| Total non-H atoms | 2379 |
| RMSD from ideal geometry |  |
| Bond lengths [Å] | 0.010 |
| Bond angles [°] | 1.60 |
| **Ramachandran** |  |
| favoured | 91.7% |
| allowed | 8.3% |
| outliers | 0% |

The rhodesain pro-domain is mainly α-helical. The catalytic core domain follows the typical two-lobed fold of the papain family of cysteine proteases (C1 family of peptidases) with an α-helical L-domain and a β-sheet containing R-domain. The cleft between these domains harbors the active site triad residues C150 (mutated to alanine in our construct), H287 and N301 (Fig. 2B). In the same position as in papain (PDBs: 9PAP, 3TNX) ^[30,31]^ and related proteases, pro-rhodesain harbors three intramolecular disulfide bonds between residues C147-C188 and C181-C226 in the L-domain as well as between C280-C328 in the R-domain (Fig. S5). As apparent from the electron density, these residues all form disulfide bridges in our structure. The backbone RMSD between our structure of the pro-rhodesain core catalytic domain (PDB: 7AVM, residues 127-342) and the previously determined structure of mature rhodesain (PDB: 2P7U)^[25]^ is 0.39 Å. This demonstrates that the presence of the pro-domain does not affect the structure of the catalytic domain, including the relative side-chain orientations of the residues of the active site catalytic triad (Fig. 2B).

The pro-domain constitutes about one third of the protein in immature cathepsin proteases and consists of three helices, H1-H3 (Fig. 2). In pro-rhodesain, the N-terminal helix 1 (H1, residues R40^pro^ -K47^pro^) is shorter compared to H1 from other crystallized CathL zymogens. The lack of resolution for the first 18 amino acids indicates that this region may be highly flexible and potentially not structured in pro-rhodesain. H1 is connected to the orthogonal helix 2 (H2, residues A55^pro^-A76^pro^). Via a long loop, H2 then leads into a long third helix (H3, residues E96^pro^-K113^pro^). This helix covers the active site and constitutes the main difference between the pro-domain of rhodesain and other CathL-like protease pro-domains (see below).

Besides the identity of their active site residue, proteases are grouped into clans (based on their evolutionary origin) and families (based on their similarity in amino acid sequence).^[32]^ CathL and CathB proteases both belong to the CA clan and the C1 family and share a papain-like fold. Nevertheless, their pro-enzymes differ in some features such as the presence of the ER(I/V)FNIN motif. This conserved motif is only found in CathL-like precursors within helix 2 of the pro-domain. In the trypanosomal CathL proteases, it is slightly modified.^[33]^ Here, the ER(A/V)FNAA motif (consensus sequence Ex_3_Rx_2_(A/V)(F/W)x_2_Nx_3_Ax_3_A, with x = any amino acid) (Fig. 3A, S6) is involved in the correct folding of the pro-domain and mediates intradomain interactions.^[34,35]^ For instance, F65^pro^ of the rhodesain pro-domain’s ER(A/V)FNAA motif mediates the interaction with H1 via extensive aromatic stacking (F41^pro^/F44^pro^ in H1 and F62^pro^/F65^pro^ in H2). The importance of these interactions is highlighted by the fact that it is also observed in e.g. pro-papain (PDB: 3TNX; F48^pro^/W51^pro^, F69^pro^/F72^pro^)^[31]^ and at least three of the four aromatic residues involved in this interaction are highly conserved across the pro-domains of other cathepsin zymogens, including human pro-CathL (PDB: 1CJL; W29^pro^/W32^pro^-W52^pro^) ^[36]^ (Fig, 3, S6).

**Fig. 3:**
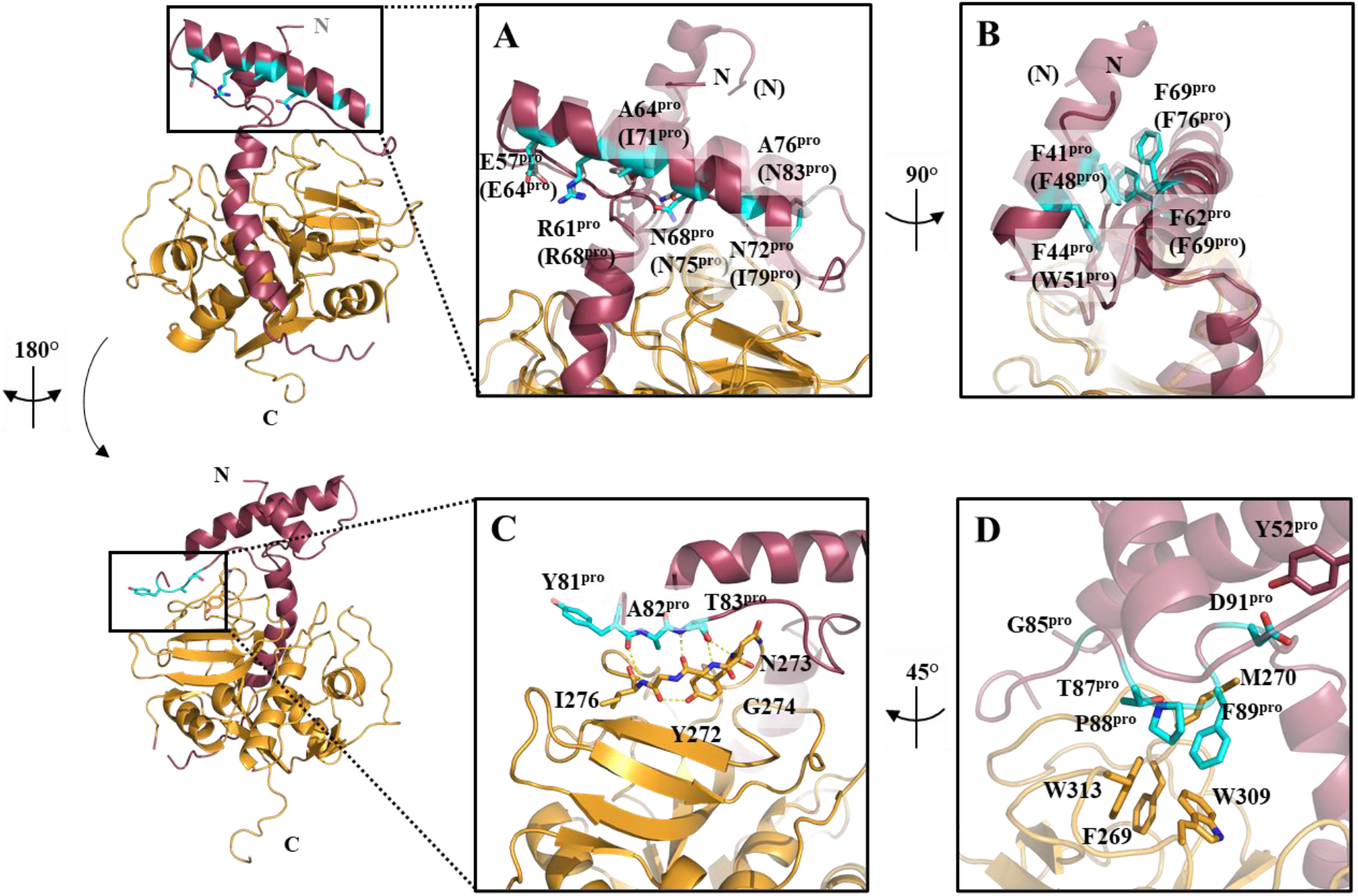
Conserved motifs within CathL-like proteases. Structure alignment of pro-rhodesain (solid) and pro-papain (transparent, PDB: 3TNX).^[31]^ The amino acid numbers for both enzymes are based on their uniprot entries (ID:Q95PM0, pro-rhodesain; ID:P00784, pro-papain). Pro-papain residues are displayed in brackets. A) Amino acids of the conserved ER(A/V)FNAA (cyan) or ER(A/V)FNIN motifs (grey) within the pro-peptide (red) are shown as sticks. B) Conserved aromatic π-stacking between residues from H1 and H2 of the pro-peptide. C) The pro-peptide binding loop (PBL) forms a β-sheet with a loop from the R sub-domain of the core protease. Y272 stabilizes a sharp kink by forming an H-bridge to the backbone of I276 with its aromatic OH-group. D) Interactions of the GNFD-motif. D91^pro^ interacts via an H-bridge with Y52^pro^ of the first pro-domain loop, while P88^pro^ and F89^pro^ are buried in a hydrophobic pocket formed by F269, W309 and W313 in the catalytic domain.

In most zymogens, H2 is longer than in our pro-rhodesain structure and we do not observe density for the C-terminal end of this region (residues 75 ^pro^ −76 ^pro^). In the pro-papain structure (PDB: 3TNX), ^[31]^ this region has been modeled as part of the H2/H3 linker and consequently, H2 is of similar length compared to what we observe for pro-rhodesain. This, together with the elevated B-factor values for this region, indicates that the C-terminal end of H2 may be flexible in the pro-domain of pro-rhodesain as has been stated for hsCathL.^[37]^

H2 connects to Helix 3 (H3, residues 96 ^pro^ −113 ^pro^) via a long linker. This linker partially forms an antiparallel β-sheet (residues 80 ^pro^ −83 ^pro^) with the pro-peptide binding loop (PBL) (residues 271-275 (Fig. 3C), a loop extending from the R sub-domain of the catalytic core domain. Notably, both, the short β-sheet in the H2/H3 linker of the pro-domain and the interacting PBL in the protease domain display very low B-values indicating high rigidity (Fig. 2C, D). In cathepsin pro-domains, the β-strand in the H2/H3 linker is typically followed by a GNFD motif with the GxNxFxD (x = any amino acid) consensus sequence, which is required for proper protein folding and autoactivation.^[38]^ In pro-rhodesain, this sequence corresponds to the GNFD-like motif G_85_VT_87_PF_89_SD_91_. In our structure, residues from this motif form multiple contacts within both the pro-domain and to the core protease domain. From the pro-domain, residues G85^pro^ together with Y81^pro^ and T83^pro^ within the H2/H3 loop’s β-strand interact via backbone hydrogen bonds with the backbone of residues I276 and G274 in the catalytic domain (Fig. 3C, D). In addition, the side chain of T83^pro^ forms a hydrogen bond to the backbone amide of G274 and the side chain carbonyl group of N273 interacts with the backbone amide of G85 ^pro^. Simultaneously, the side chain amide group of N273 is involved in a hydrogen bond with the backbone carbonyl of its preceding amino acid, Y272, which introduces a sharp kink in the peptide chain. This brings the hydroxyl group of the sidechain of Y272 in close proximity to the backbone carbonyl group of G275 to form another H-bond while the aromatic ring of Y272 stacks with the peptide bond between N273 and G274, thus providing further stabilizing interactions. This intricate interaction network may explain why the core protease domain’s PBL β-strand harbors a di-glycine motif (G274/G275) and provides some insights why this region could be crucial for protease folding^[36]^, since larger amino acid side chains may sterically interfere with these interactions. Indeed, residues Y271, G272 and G275 are highly conserved among CathL proteases (Fig. S6).

An interaction network involving mostly hydrophobic and aromatic residues in the R domain of the core protease domain positions the remainder of the pro-domain H2/H3 linker. Notably, the side chains of P88^pro^ and F89^pro^ dip into a hydrophobic basket formed by the sidechains of F269, M270, W309 and W313 in the R-domain of the core protease (Fig. 3D). In addition, the backbone carbonyl of M270 forms a hydrogen bond with the side chain of T87^pro^ while its sidechain forms a hydrophobic interaction with F97^pro^ in the first turn of H3, the long α-helix covering the entire length of the cleft between the L- and R-domains of the catalytic domain. In all available structures of other cathepsin pro-enzymes, this region is mainly unstructured (see below) (Fig. 4A). This α-helix therefore presents the most notable structural difference between pro-rhodesain and closely related non-trypanosomal proteases.

**Fig. 4:**
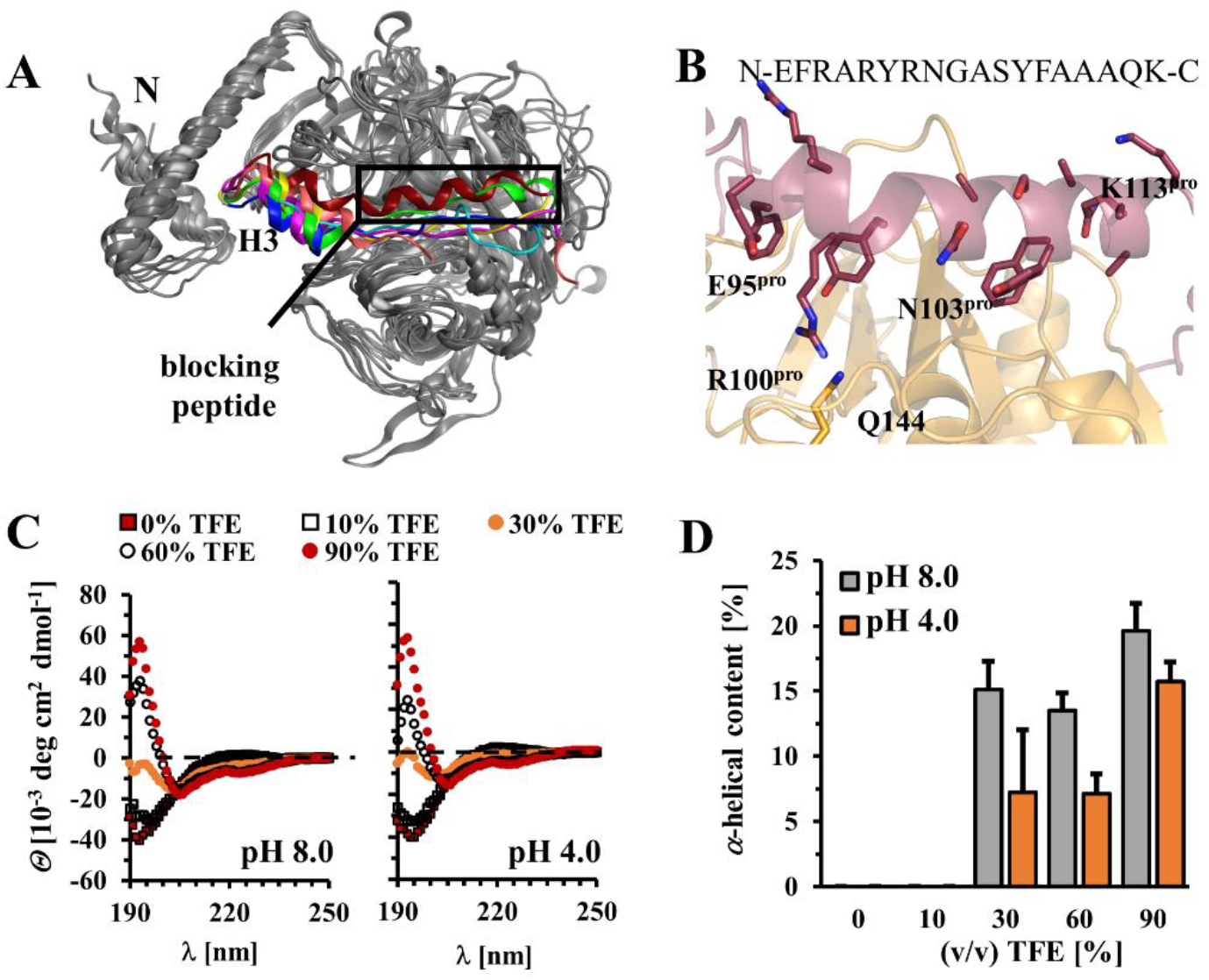
The trypanosomal pro-CathL blocking peptide adopts an α-helical structure so far not observed in non-parasitic CathL zymogens. A) Comparison of blocking peptide in pro-rhodesain and other CathL zymogens (green = *F. hepatica*, PDB: 2O6X; blue = *H.sapiens*, PDB: 1CJL; pink = *C. papaya*, PDB: 1PCI; beige = *T. molitor*, PDB: 3QT4; turquoise = *H. sapiens* (Pro-CathX), PDB: 1DEU; yellow = *H. sapiens* (Pro-CathK), PDB: 1BY8). B) Zoom of helix 3 of the pro-rhodesain pro-peptide (red) covering the active site cleft of the core protease domain (gold). The amino acid sequence of H3 is shown on top and side chains are presented as sticks. H3 harbors a kink between R102^pro^ (residue not visible in this orientation) and N103^pro^. The side chains of R100^pro^ and Q144 (within the catalytic domain, gold) interact. C) CD-spectra of the trypanosomal blocking peptide at various TFE concentrations and different pH values (for CD spectra of the human CathL blocking peptide see Fig. S7). D) The α-helical content of the blocking peptides calculated with the online tool BeStSel (http://bestsel.elte.hu/index.php).^[29]^

### Helix H3 in pro-rhodesain differs from that of other non-parasitic pro-cathepsins

The residues in H3 facing the catalytic domain are involved in numerous, mostly hydrophobic interactions with the catalytic domain, thereby efficiently blocking the rhodesain active site (Fig. 4A). In agreement with structures from multiple CathL pro-enzymes, the pro-rhodesain H3 helix starts out as a regular α-helix. However, and so far unique to pro-rhodesain, a bulge forms around residues N103^pro^ and G104^pro^, whose backbone amides are not involved in hydrogen bonds. The hydrogen bond acceptor group for the amide group of G104^pro^ in the context of a regular α-helix would be the oxygen of the backbone carbonyl of R100^Pro^. This group, however, is 5.1 Å away from the G104^pro^ amide group thus precluding formation of a hydrogen bond at this site and instead locally distorting the α-helix in proximity to the active site of the core protease (Fig 4B). Importantly, this bending of the α-helix enables the guanidino group of R100 ^pro^ to interact with the side chain of Q144 in the core protease domain. Notably, Q144 is conserved across all CathL proteases while R100^pro^ is conserved in the majority of homologous trypansomal CathL pro-proteases (Fig. S6). Below the bulge, H3 continues as a regular α-helix up to residue K113^pro^ from where the pro-domain leads into the R-domain of the catalytic core domain via a C-terminal 15 amino acid unstructured linker (Fig. 2, Fig. 4B). The electron density in our X-ray structure allows tracing the backbone of the entire H3 α-helix unambiguously for all residues in H3 (E95^pro^ −K113^pro^) (Fig. S5B). The B-factor values for the two N-terminal helical turns of H3 are among the highest in the pro-domain while in the entire remainder of this helix following the bulge, the B-factors are among the lowest (Fig 2C, D). The sequence of the N-terminal region of H3 preceding the bulge, which is α-helical across all available CathL pro-domain structures, is generally well conserved across CathL enzymes (Fig. 4A, S6). However, and in contrast to the continuous α-helix observed for H3 in pro-rhodesain, the structures of all other available, non-trypanosomal pro-CathL pro-proteases display a mostly unstructured peptide in the C-terminal end of H3 (Fig. 4A). Despite the differences in secondary structure within this blocking peptide, its overall position relative to the cleft of the core protease domain is nonetheless highly similar. Furthermore, this region shows no sequence similarity between *T. brucei* rhodesain and other, non-parasitic pro-CathL enzymes, but extremely high sequence identity across trypanosomal pro-cathepsins (Fig. S6).

The observation that the region of the pro-domain blocking the protease active site is α-helical in pro-rhodesain, but not in any of the other pro-Cath(L) structures determined to date (Fig. 4A), prompted us to test whether the intrinsic propensity of this region to form an α-helix is higher in pro-rhodesain compared to related proteases. We used the isolated blocking peptides from *T. b. rhodesiense* rhodesain (residues R102^pro^ − K114^pro^) and human CathL (residues N112^pro^ − R120^pro^) to record CD spectra at pH 8 and pH 4 and with increasing amounts of trifluoroethanol (TFE) (Fig. 4C, D, S7). TFE induces secondary structure formation by promoting hydrogen bond formation.^[39,40]^ Both peptides are mainly unstructured in the absence of TFE. For the peptide from human CathL, even high TFE concentrations (up to 90% v/v) do not induce significant secondary structure content (Fig. S7). This is in agreement with the highly disordered protein region observed in the human pro-CathL crystal structure (PDB: 1CJL).^[36]^ In contrast, the isolated rhodesain pro-domain peptide becomes α-helical already at low TFE concentrations (Fig. 4 C, D). The variation in pH had no significant effect on secondary structure propensity of either blocking peptide. Likewise, the CD spectra of the purified pro-rhodesain C150A construct are also virtually identical at high and low pH (Fig. S8).

### Pro-peptide binding site interactions resemble the binding mode of the K11777 inhibitor

Proteases recognize their substrates by engaging with the substrate amino acid side chains in specific pockets. These pockets are numbered outwards from the cleavage site towards the substrate’s N- (binding sites S1-Sn) and C-terminus (prime sites S1’-Sn’).^[41]^ In pro-cathepsins, the N- to C-terminal orientation of the pro-domain blocking peptide is inverted with respect to the interaction of a putative peptide substrate (Fig. 5A). In the pro-rhodesain structure, the side chains of F108^pro^ and Y107^pro^ occupy the S2 and S3 pockets, while the S’ and S1’ sites are partially occupied by the side chains of Y82^pro^ and R81^pro^. A glycerol molecule sits in the center of the substrate binding site (Fig. 5A, B). The side chains of the pro-domain residues Y107^pro^ and F108^pro^ in the S3 and S2 pockets also overlap with the binding sites of the phenyl- and *N*-methylpiperazine groups of the K11777 inhibitor as observed in the crystal structure (PDB: 2P7U) (Fig. 5C, D).^[25]^ We also looked at the position of the peptidic substrate Z-Phe-Arg-AMC used for activity assays by molecular docking and observed similar occupations of the substrate binding pockets of rhodesain (Fig. 5E, F).

**Fig. 5:**
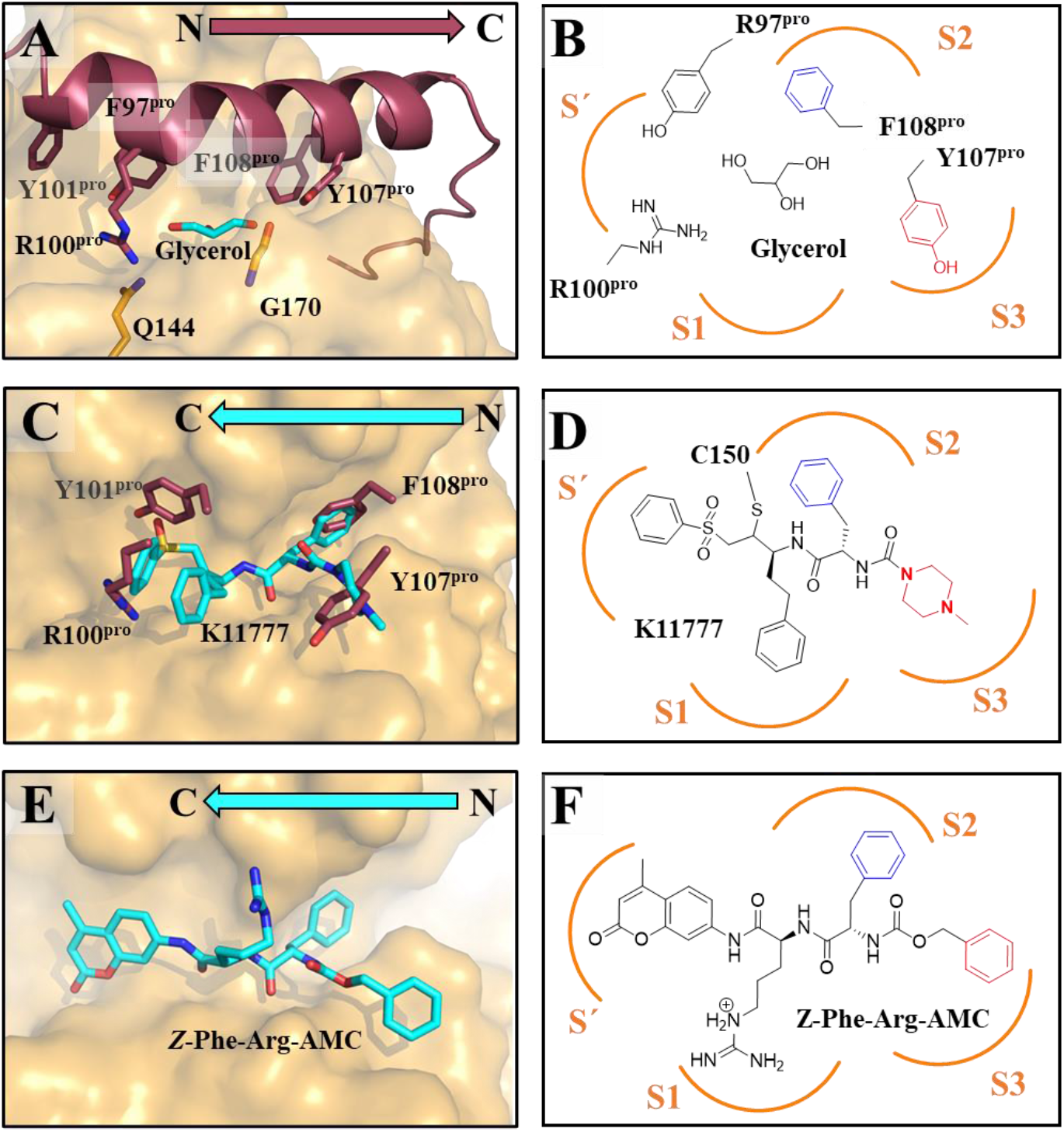
Side chain interactions of H3 with the protease active site cleft are similar to those of the peptide-based inhibitor K11777. A) Interactions between the active site cleft and pro-peptide side chains. B) 2D-presentation of the H3 side chains within the active site cleft. C) Crystallographic binding mode of K11777 (cyan carbon atoms) in complex with rhodesain (gold surface) in superposition with pro-peptide residues occupying S1’, S2 and S3 (red). D) 2D-presentation of K11777 in the active site based on the crystal structure 2P7U. Orientation of the inhibitor within the active site leads to a comparable binding mode as H3 in B). E, F) Binding mode and 2D-presentation of Z-Phe-Arg-AMC docked into the binding pocket of rhodesain (FlexX score −23.98 kJ/mol).

### pH-dependent auto-cleavage of pro-rhodesain can occur intra- and intermolecularly

For the physiological activation of rhodesain and other CathL-like proteases, the pro-domain has to undergo cleavage at low pH, and it has been suggested that this process can occur inter- or intramolecularly (or both).^[21,42,43]^ In our hands, folded pro-rhodesain harboring the active site mutation C150A purified from *E. coli* is not cleaved, ruling out a role for other *E. coli* proteases in the cleavage process of heterologously expressed pro-rhodesain. Lowering the pH for purified wildtype pro-rhodesain leads to cleavage indicating that intramolecular autocleavage plays at least some role in pro-rhodesain processing (Fig. S4). However, this does not exclude the possibility that cleavage also occurs intermolecularly. Intermolecular protease cleavage can be easily probed by mixing catalytically inactivated enzymes with a wildtype protease.^[44]^ Incubating an excess of purified, catalytically inactive pro-rhodesain C150A with catalytically active, mature, wildtype rhodesain led to a continuous decrease of the high molecular band corresponding to pro-rhodesain in a gel-shift assay (Fig. 6A, B). Importantly, this process is pH dependent and lower pH values result in more efficient autocleavage.

**Fig. 6:**
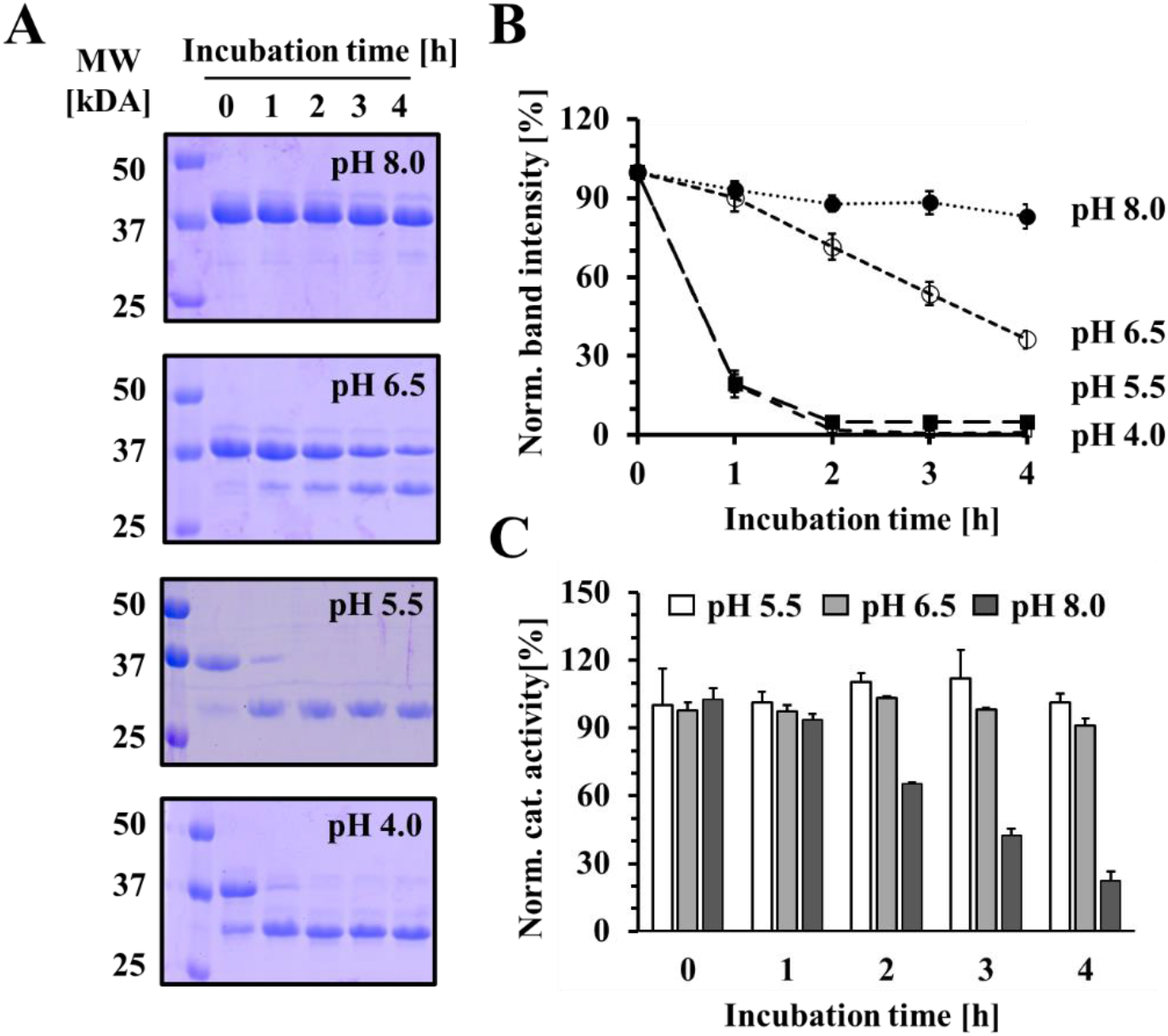
pH-dependent auto-cleavage of pro-rhodesain. A) Co-incubation of pro-rhodesain C150A with sub-stoichiometric amounts of mature rhodesain (100:1 mol:mol) at different pH values leads to cleavage of the 37 kDa pro-enzyme to the mature form at 26 kDa as observed on SDS-PAGE. Lower pH values lead to more efficient cleavage. B) Densitometric analysis of gels in (A). Shown is the decay in the bands at 37 kDa corresponding to pro-rhodesain C150A. Error bars depict SDE from three independent measurements. C) Activity of wildtype rhodesain probed by a fluorescence peptide cleavage assays under the conditions used in (A). Shown are values normalized to rhodesain activity determined at pH 5.5 with a fluorogenic peptide.

More efficient pro-rhodesain C150A cleavage at lower pH values could be due to a higher intrinsic activity of the mature wt rhodesain and/or changes in the structural dynamics of the pro-enzyme and consequently a higher accessibility of its cleavage site. To probe the first possibility, we tested the residual activity of mature rhodesain at different pH values by pre-incubating it with the fluorogenic peptide-based substrate (Fig. 6C). While the activity of rhodesain slowly decreases at pH 8 over the course of multiple hours, this drop in activity is not sufficient to explain the near-absence of pro-domain cleavage within the same time-frame and the differences in pro-rhodesain C150A cleavage at pH 6.5, pH 5.5 and pH 4 (Fig. 6). We therefore investigated pro-rhodesain dynamics at high and low pH values, in particular the relative motion of the pro-domain to the core protease domain.

### Dynamics of pro-rhodesain probed by PET-FCS and MD simulations

To investigate the intramolecular dynamics of the pro-domain in relation to the core protease domain, we used photoinduced electron transfer fluorescence correlation spectroscopy (PET-FCS).^[27]^ Here, the dynamics of a protein can be probed via contact-induced fluorescence quenching, i.e. a photoinduced electron transfer (PET) reaction of an extrinsic fluorophore attached to a cysteine side chain with the side chain of the natural amino acid tryptophan.^[45]^ Fluorophore and indole group form a van der Waals contact at distances below 1 nm which can be disrupted or formed by conformational changes of a protein. Such distance changes are thus translated into fluorescence fluctuations and the combination of the PET fluorescence reporters with fluorescence correlation spectroscopy facilitates the single-molecule detection of fast protein dynamics with nanosecond time resolution.^[27,46]^ We have also used this approach to probe the binding kinetics of mature rhodesain with its inhibitors (Johe et al., in prep.). Pro-rhodesain contains eight native cysteine residues, the three cysteine pairs forming disulfide bonds in the core protease domain (Fig. S5), the active site residue C150 and a cysteine in the pro-domain (C22). To avoid unspecific labeling of the enzyme with a fluorophore, we thus mutated both the active site cysteine (C150A) to stabilize the pro-enzyme as well as C22 (C22S).

For the pro-rhodesain C22S^pro^, C150A double mutant, protein modified with an AttoOxa11-dye was hardly detected, showing that the remaining six cysteine residues were protected against fluorophore labeling due to their involvement in disulfide bridges. Subsequently, two new cysteine residues were introduced in the C22S^pro^, C150A background of pro-rhodesain, either at the C-terminal end of pro-domain helix 1 (V51C^pro^) or right above the bulge in helix 3 in the vicinity of the active site (A99C^pro^). For PET fluorescence measurements, a tryptophan mutation at position Q146 in the L domain of the core protease domain was introduced in the vicinity of V51C^pro^ and A99C^pro^ (Fig. 7). Neither the cysteine mutations nor the additional tryptophan mutation perturbed the secondary structure of pro-rhodesain and all constructs could be successfully labeled with an AttoOxa11-dye as assessed by steady-state fluorescence spectroscopy (Fig. 7, 8A, B). As expected, due to the large distance from the fluorophore as apparent from our structure, the eight native tryptophan residues in pro-rhodesain did not lead to fluorescence quenching of dyes attached to residues V51C^pro^ or A99C^pro^, while introduction of Q146W did. The AttoOxa11-labeled V51C^pro^/Q146W and A99C^pro^/Q146W pairs can thus be used as sensitive fluorescence reporters for the relative motion of the pro-domain with regard to the protease domain of pro-rhodesain.

**Fig. 7:**
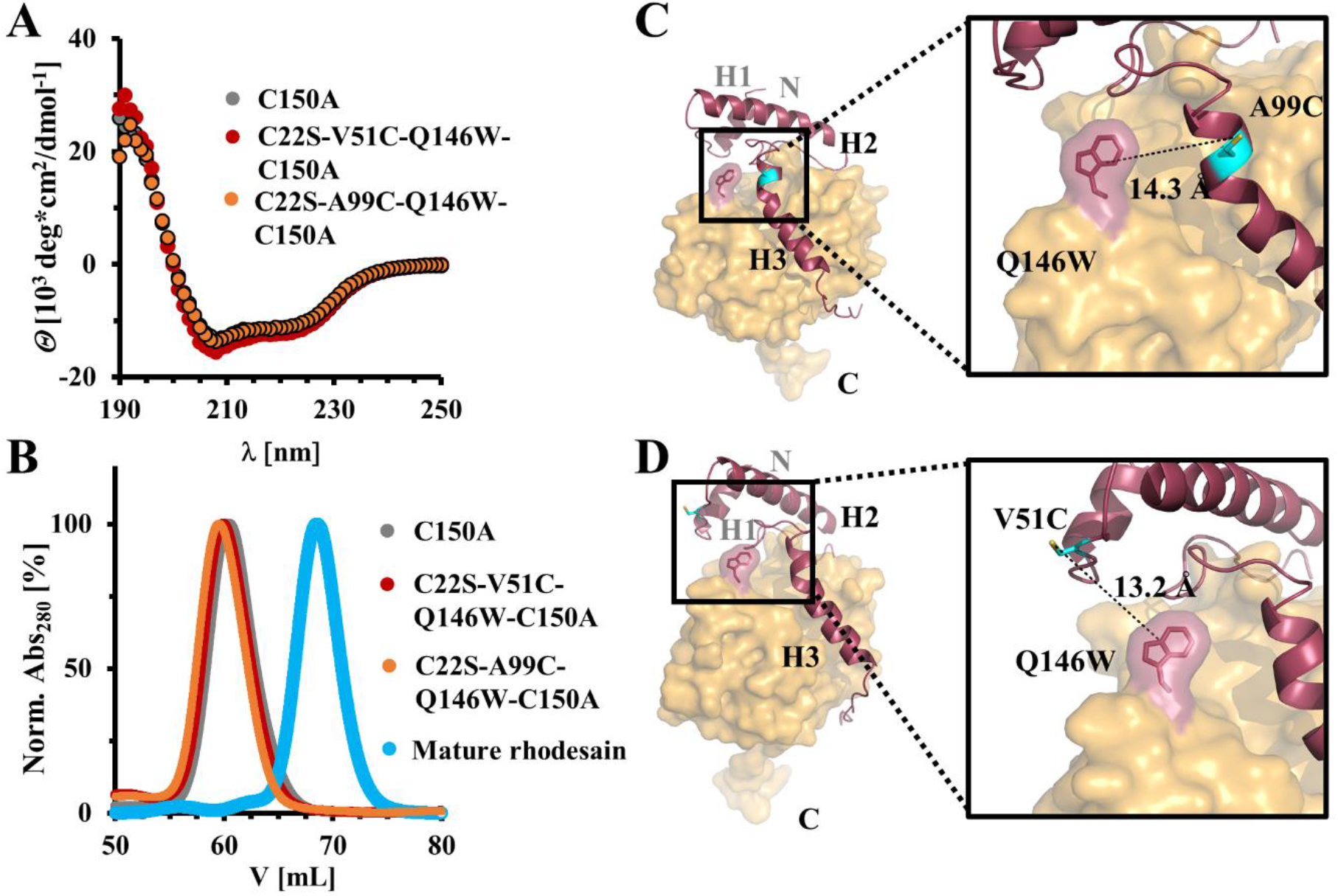
Pro-rhodesain mutants used for PET-FCS measurements. Introduction of the required Trp and Cys mutations does not change the secondary structure as assessed by A) CD spectroscopy and B) size exclusion chromatography of rhodesain variants. C, D) Position of the Trp/Cys pairs used for PET-FCS to probe interdomain dynamics between pro-domain and core protease domain.

**Fig. 8:**
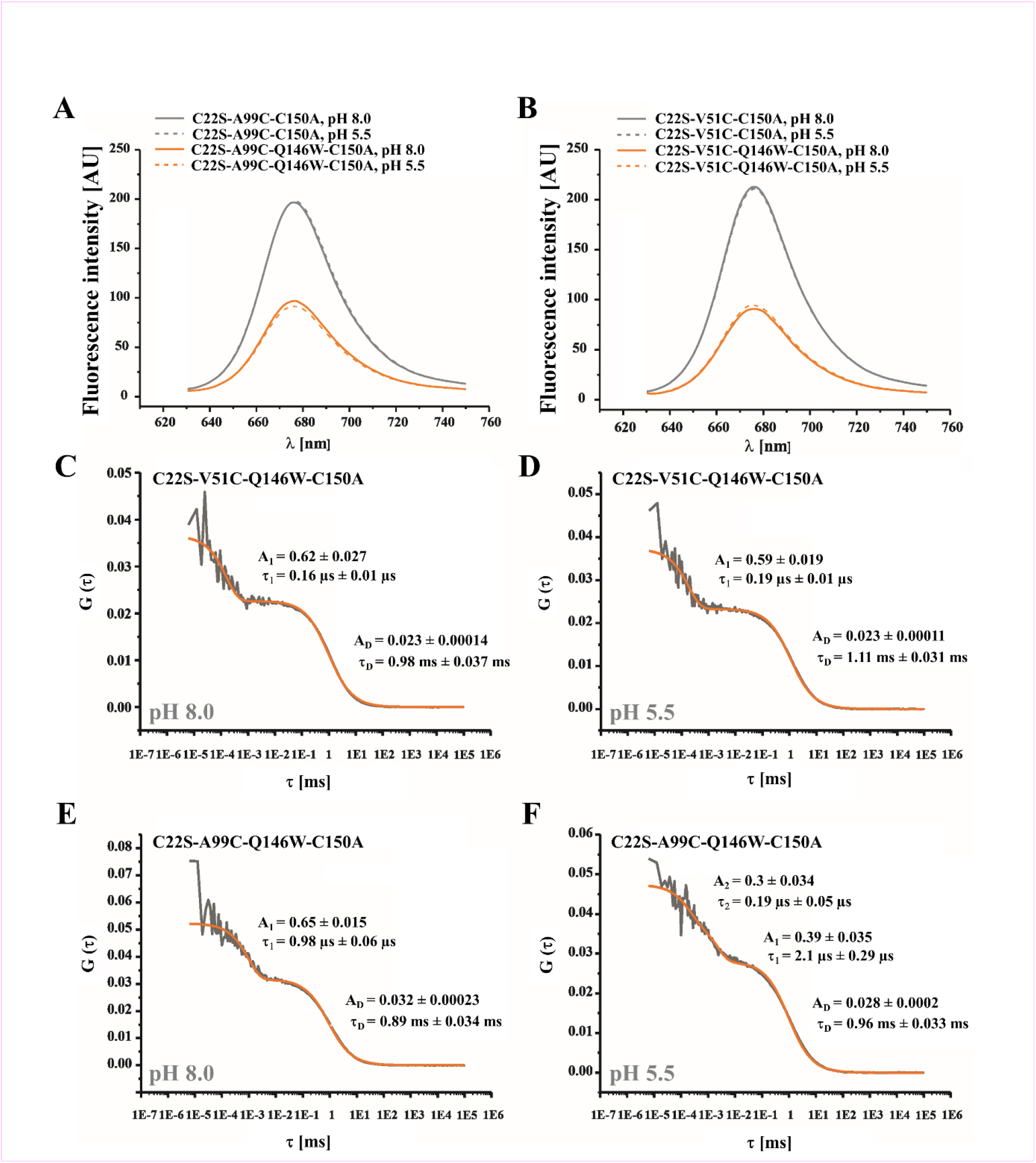
PET-FCS on pro-rhodesain. A, B) Steady-state fluorescence spectra of pro-rhodesain constructs, labeled at position A99C^pro^ or V51C^pro^ using AttoOxa11, show efficient quenching of the label by the Trp residue Q146W in the core protease domain while pH has a negligible effect on the spectra. C-F) FCS autocorrelation functions (G(τ)) measured from pro-rhodesain constructs at 37 °C under solution conditions specified in panels. Data shown in panels C-E were fit using a model for molecular diffusion containing a single exponential relaxation (orange line). Data shown in panel F were fit using a model for molecular diffusion containing two single-exponential relaxations (orange line). Amplitudes (A_n_) and time constants (τ_n_) extracted from fits are shown in panels.

No changes in fluorescence intensity upon reducing the pH are observed in our fluorescence spectra of pro-rhodesain (Fig. 8A, B). To probe whether there are nonetheless dynamic ramifications for pro-rhodesain in response to pH changes, we additonally carried out PET-FCS measurements with the dye/Trp pair containing proteins at pH 8 and pH 5.5 (Fig. 8C-F). Here, the drop in pH led to a small increase of the diffusion time constant τ_D_ of all tested mutants indicative of a larger hydrodynamic radius under acidic conditions. Since this behavior is observed for all proteins, it might be interpreted as a global structural rearrangement of pro-rhodesain or “loosening” of the overall structure.

Interestingly, the pro-rhodesain construct with the A99C^pro^/Q146W dye/quencher pair also showed a pH-dependent change in sub-ms fluorescence fluctuations (Fig. 8E, F). Fluorophores attached to residue A99C^pro^, within the “bulge” of H3 covering the protease active site, yield a monoexponential autocorrelation curve with a 1 μs time constant at pH 8. At pH 5.5, the same construct shows a biexponential trace with time constants of 0.21 and 2.1 μs (Fig. 8E, F). The amplitudes are the same in both cases, showing that the total fluorescence quenching is similar at high and low pH, but that the local dynamics of the system are more heterogeneous under acidic conditions. In contrast, the pro-rhodesain construct harboring the V51C^pro^/Q146W dye/quencher pair, with V51C^pro^ at the C-terminal end of H1 in the pro-domain shows only a modest response to the change in pH (Fig. 8C, D).

To shed additional light on the pH-dependent autocleavage process, we carried out molecular dynamic (MD) simulations with the pro-enzyme at pH 4 and 8, respectively. While large conformational changes are beyond the time scale of MD simulations (20 ns), “loosening” of the pro-domain may be indicated by an increased flexibility and the reduction of interactions between residues sensitive to pH changes.^[37]^ All simulations showed a high stability within the given time frame (Fig. S9A). The root mean square fluctuation (RMSF, Fig. S9B) and deviation (RMSD, Fig. S9C) analysis indicated the same regions for high flexibility as found by B-factor analysis (Fig. 2D). The evaluation of the interaction energies between the pro-domain and main domain indicate significantly reduced interactions between the two domains at pH 4 compared to pH 8 while higher interaction energies were found within the main domain at pH 4 (Table S1).

Since pro-rhodesain is cleaved efficiently at low pH and our PET-FCS data indicate that the dynamics of H3 increase under acidic conditions, we wondered whether protonatable groups mediating interactions between H3 and the core protease play a role in the cleavage process. We thus performed a detailed analysis of the H-bond interaction profile for the MD simulations with focus on titratable groups (Tables S2-S7). This revealed some changes in the interaction patterns of specific amino acids for different protonation states. A salt bridge connecting the C-terminal end of H3 (R116^Pro^) with the core protease (D194 and D242) was identified as especially interesting. For MD simulations at pH 4, D194 was protonated and therefore able to interact with D242 via a hydrogen bond (64-74% of the simulation time). When D194 was deprotonated during simulations at pH 8, this interaction was no longer possible and D194 and D242 were able to interact with the side chains of R114^Pro^ and R116^Pro^ *via* ionic interactions (Table 2). This provides a hint for putative stabilizing interactions between pro- and main domain at higher pH values that can be replaced by intra-domain interactions at acidic pH.

**Table 2:**
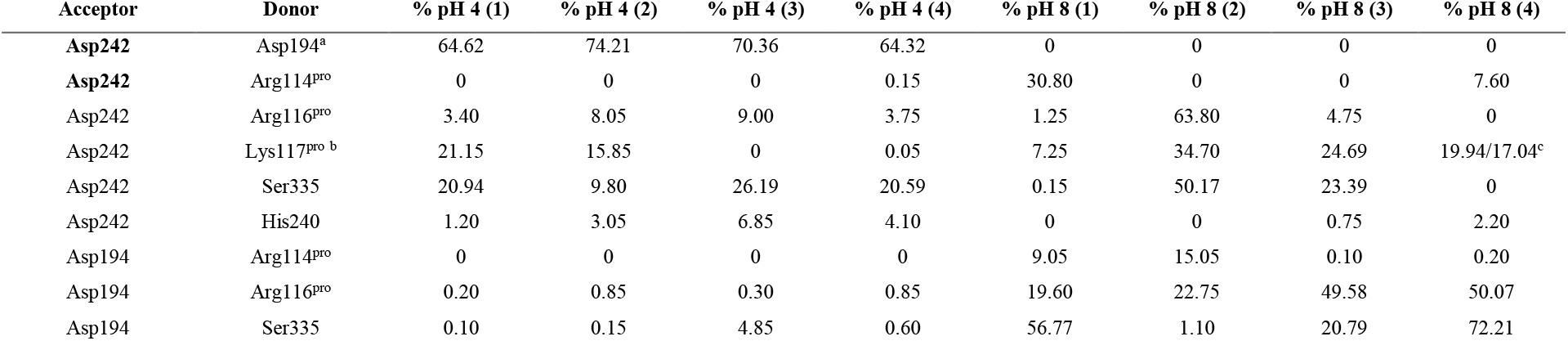
Relative occurrence of side-chain hydrogen bonds between selected pH-sensitive residues. Four respective MD simulations at pH 4 or pH 8 were carried out. ^a^ denotes donor functionality only in protonated state (pH 4). ^b^Interaction with NH backbone. ^c^Interaction with side chain amine.

Incidentally, the residues involved in this putative interdomain “lock” at pH 8 are highly conserved across trypanosomal pro-CathL proteins (Fig. 9A, S6). Despite the lack of overall sequence similarity, this interaction is even replicated in non-trypanosomal CathL proteins (Fig. 9A). In *C. papaya* pro-caricain, the positions of the corresponding basic and acidic residues are interchanged further underscoring the fact that these amino acids play an important role and may co-evolve.

**Fig. 9:**
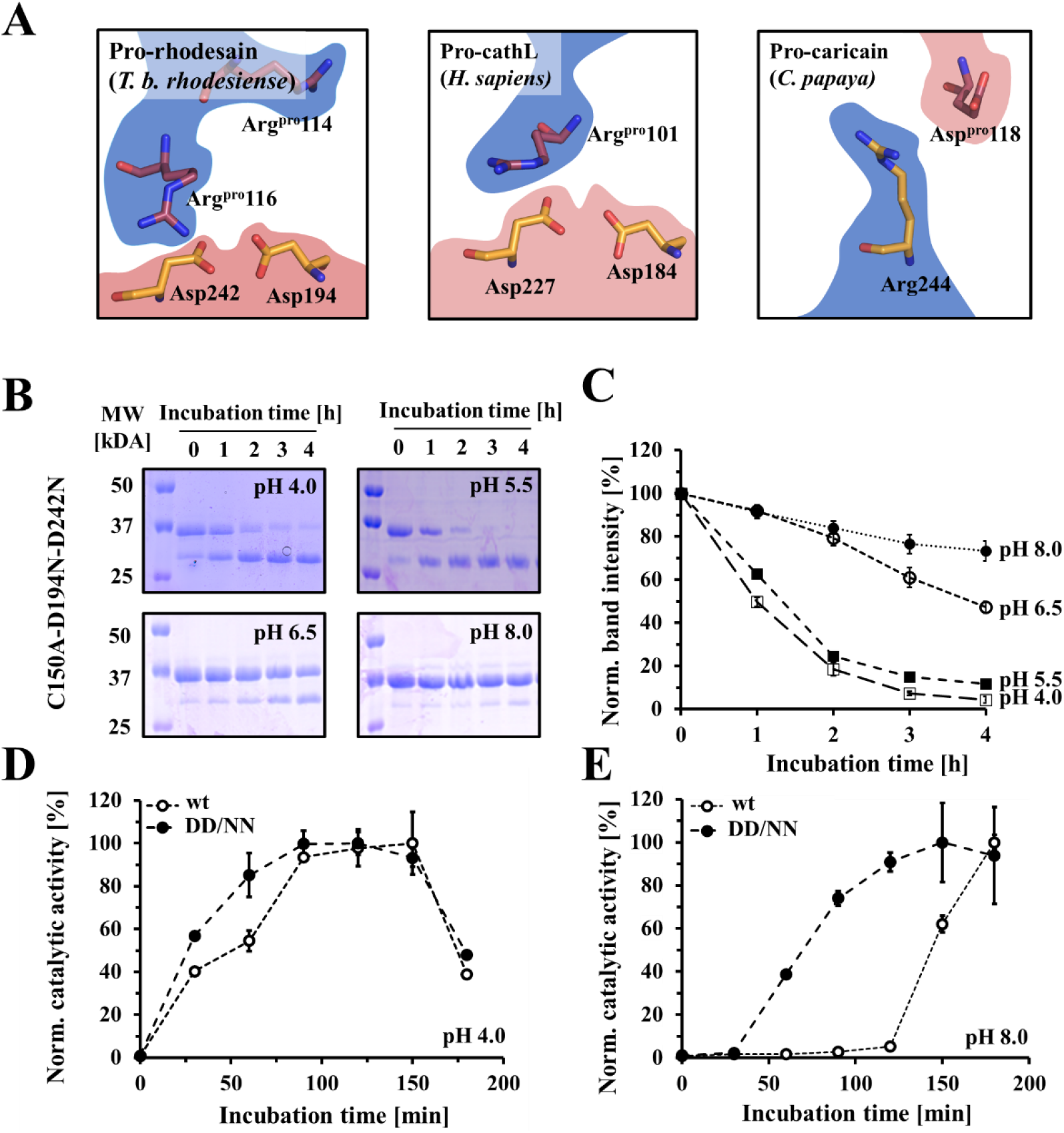
An interdomain salt bridge influences the pH sensitivity of pro-rhodesain autoactivation. A) Conserved salt bridges between C-termini of blocking peptides and catalytic domains in crystal structures from pro-CathL proteases from different species (*T. b. rhodesiense*, PDB: 7AVM; *H. sapiens*, PDB: 1CJL; *C. papaya*, PDB: 1PCI). The amino acids of the pro-domain (red) and the catalytic domain (orange) are shown as sticks and highlighted by their charge (blue = positive, red = negative). In the case of *C. papaya* pro-caricain, a polarity switch occurs. B) pH-dependent intermolecular (*trans*) cleavage of inactive pro-rhodesain C150A DD/NN was probed by incubating the purified protein with a sub-stoichiometric amount of mature rhodesain (100:1 mol:mol) at pH 4.0, 5.5, 6.5 or 8.0 at 37 °C. Samples were taken at indicated time points and run on 15% SDS-PAGE. The data show that pro-rhodesain C150A DD/NN can be digested intermolecularly by mature rhodesain. C) Intensities of the pro-rhodesain band at ~37 kDa were determined with ImageJ and relative values compared to t0 plotted against the incubation time. D, E) Purified wt pro-rhodesain and the pro-rhodesain DD/NN-mutant, importantly both harboring the active site cysteine to enable autoactivation, were incubated at pH 4.0 (D) and pH 8.0 (E) at 37 °C. Inhibition with PMSF was removed by adding DTT and the catalytic activities of each sample were determined with a fluorescence based enzyme assay to probe the respective autoactivation of both proteins at thhe indicated time points. The ratio of the measured activity at a given time point to the maximum values was plotted against the respective time point.

To probe the effect of this putative interaction on pro-rhodesain autocleavage, we mutated the two aspartic acids into non-protonatable asparagine residues in the C150A background yielding pro-rhodesain C150A, D194N, D242N (C150A DD/NN). Adding mature wt rhodesain to this mutant, we probed its digestion via the same gel-shift assay used previously to establish intermolecular cleavage (Fig.9B, C).

The inactive pro-rhodesain C150A DD/NN mutant was efficiently cleaved at low pH values, although it displayed a slightly decreased cleavage behavior compared to the construct harboring only the C150A active site mutation (Fig. 6). The interdomain interaction at this site therefore does not significantly affect the intermolecular cleavage process. To probe whether it may instead play a role in the intramolecular cleavage, both wt pro-rhodesain and pro-rhodesain DD/NN without the active site mutation were expressed in *E. coli* and obtained in their pro-forms by purifying them at pH > 7 and in the presence of the protease inhibitor phenylmethylsulfonyl fluoride (PMSF) to avoid autocleavage.^[47]^ After purification, the PMSF inhibition was reversed by the addition of DTT and the pH was adjusted to either pH 8 or pH 4. Subsequently, the enzymes’ auto-activation was monitored using the fluorescence-based peptide cleavage assay (Fig. 9D, E). At pH 4, the enzymes did not show differences regarding their activation behavior, however, at higher pH values, the DD/NN mutant showed a significant enhancement of autoactivation.

## Discussion

Typanosomes are responsible for devastating diseases affecting both humans and their lifestock. For survival and propagation in the host, these parasites rely on proteases such as rhodesain. Similar to related cathepsin proteases, rhodesain is expressed as a zymogen with a pro-domain that maintains its inactive state until the protein is correctly delivered into the acidic environment of the lysosome. Here, we present the first trypanosomal pro-CathL structure, *T. brucei rhodesiense* pro-rhodesain, as an important intermediate of the protease maturation process.

While the overall architecture of pro-rhodesain agrees with related pro-enzymes including human cathepsin L or *C. papaya* papain,^[36,48]^ pro-rhodesain displays an unique extended α-helix covering the cleft between the L- and R-lobes of the core protease domain (Fig. 2). A pronounced bulge in the region covering the protease active site disrupts the extended α-helix in H3 of the pro-domain. In the structures of currently available homologous, non-trypanosomal pro-proteases, H3 becomes unstructured at this position (Fig. 4A). It thus seems possible that deviations from a regular α-helix are a unifying structural characteristic of pro-CathL proteins although the consequences for protease maturation and cleavage remain to be investigated. As a word of caution, all available structures of pro-CathL proteases including pro-rhodesain were obtained by mutating the active site cysteine. This removes a thiolate anion from the active site and may thus locally distort the structure of the pro-domain. In addition, we observe a glycerol molecule in the active site that could lead to deviations in the side chain orientations of residues within pro-domain peptide interacting with the active site. On the other hand, our crystal structure was determined at pH ~4 where we also observed differences in local dynamics and an increase in the hydrodynamic radius compared to basic pH (see below). Changes in pH may thus affect the substrate binding pocket occupancy by pro-domain amino acid side chains and could be associated with reordering and loosening of the relative interactions between pro-domain and protease domain.

In the α-helical region following the bulge in H3, trypanosomal CathL proteases display very high sequence identity to each other but not to CathL enzymes from other organisms. The extended H3 helix of the pro-domain covering the protease active site may thus be a defining feature of trypanosomal CathL proteases and as such present a structural template to further improve the affinity or selectivity of rhodesain inhibitors, including K11777, a promising drug candidate against protozoan diseases.^[49]^ We found that the blocking peptide of the rhodesain pro-domain partially resembles K11777 binding by occupying the S2 and S3 substrate binding pockets with its aromatic and piperazine groups. In hindsight, this nicely rationalizes the original screening based design of the K11777 ^[50]^ and may provide an opportunity for further inhibitor optimizations.

In agreement with the literature,^[51]^ our data show that pro-rhodesain can be activated by pH-dependent intra- and intermolecular cleavage processes. This activation was not contingent on other proteases, although it cannot be ruled out that heterospecific cleavage events occur in the context of the cell.

To gauge whether the pro-domain undergoes molecular motions relative to the core domain, we introduced dye/tryptophan pairs for PET-FCS measurements. Globally, the diffusion times of pro-rhodesain increase at lower pH, indicating that low pH “loosens” the pro-rhodesain structure, presumably by either partial unfolding or via domain reorientations. A hypothesis for an increase in intermolecular cleavage of pro-rhodesain at low pH might be the partial detachment of the H3 helix from the core protease domain thus rendering it more accessible to other proteases. This would also lead to an increase in the hydrodynamic radius and thus the observed reduced diffusion times. Our MD simulations also suggest that the interactions between pro- and core protease domain are significantly reduced at lower pH values. Additionally, the acid-induced heterogeneiety in conformational dynamics observed for the reporter pair reporting on motions between core protease and H3 may be interpreted as local folding/unfolding events of H3 at low pH values and high crystallographic B factors indicate an increased flexibility in this region. However globally, the CD spectra of both the isolated blocking peptide and the entire pro-enzyme do not show pH-dependent secondary structure changes. Interestingly, a second PET-FCS reporter pair informing on the relative motions between H1 from the pro-peptide and the core protease domain did not exhibit changes in dynamics under all conditions tested thus suggesting that the degree of pH-dependent structural dynamics differ within the pro-domain and that H3 may be particularly sensitive to pH changes.

Looking for protonatable groups that may guide pH-dependent structural responses in pro-rhodesain, we focused on a putative salt bridge connecting the C-terminal end of H3 to the core protease domain. The residues involved are highly conserved across pro-cathepsins. When mutated, the auto-activation of the DD/NN mutant pro-protease at high pH values was significantly enhanced compared to the wt enzyme. In contrast, at low pH values, no difference in the activation rate between mutant and wt was observed. This indicates that these residues act as a pH-dependent protective lock, presumably to prevent premature protease activation at higher pH values, i.e. before the protein reaches the acidic lumen of the lysosome.

In addition, the rhodesain activation curve at higher pH values features a very shallow initial slope (Fig. 9E) indicating that very few activated protomers are present at this stage, but over time, the amount of active rhodesain rises exponentially. In agreement with models put forward for the related human CathL protease^[21]^ as well as more recently for rhodesain itself^[51]^, activiation of pro-rhodesain could follow a two-step mechanism. An initial, slow intramolecular cleavage step is delayed by the protective salt bridge between the pro-domain and the core protease as well as strong interdomain interactions that preclude premature autoactivation at higher pH values. This is followed by a faster intermolecular cleavage step. The typical pK_a_ of free aspartic acid is ~ 4 although this value can be significantly lower in the context of a salt bridge in the interior of a protein.^[52]^ In pro-rhodesain, the salt bridge is solvent exposed, a feature that may be necessary to fine-tune pH-dependent autoactivation of pro-rhodesain in a physiological pH range. An elevated pK_a_ of the two aspartic acid residues involved in the intradomain salt bridge is likely because they form a “handshake” interaction (Fig. 9).^[53]^ As the pro-protease progresses along the cellular trafficking routes from the ER via the Golgi to the lysosome and thus experiences continuously more acidic environments, their protonation can already occur at pH values around ~5.5, which would eliminate the salt bridge to the arginine residue within the protease domain and thus loosen this structural element, in agreement with our observation of altered conformational dynamics. The resulting structural and dynamic changes of H3 in relation to the core protease domain presumably enable the intramolecular autocleavage of individual pro-rhodesain molecules. Once liberated from the inhibitory pro-domain, the mature protease is now also available to activate other pro-rhodesain protomers in an intermolecular fashion, thus speeding up the maturation process of the protease population significantly. This simple model may explain how the physiological environment triggers rhodesain cleavage while simultaneously preventing premature activation. Due to the high structural similarities among pro-CathL enzymes and the additional high sequence similarity of H3 among trypansomatid proteases, our structure in combination with the presented delayed activation model may potentially act as a blueprint for these important enzymes in trypanosomatids and beyond.

## Experimental procedures

### *Rhodesain Expression and Purification from* P. pastoris *and* E. coli

The expression of rhodesain from *P. pastoris* was performed as described previously.^[9]^ For purification from *E. coli*, the *T.b. rhodesiense* rhodesain sequence (uniprot-ID.: Q95PM0) followed by a TEV cleavage site and a hexahistidine-tag was cloned between NdeI and BamHI restriction sites of a pET-11a vector by Genscript with a codon optimization. The plasmid was amplified in XL-10 cells. A GFP sequence was additionally introduced between the TEV site and the 6xHis-tag by Gibson assembly following extensive construct and purification optimization (see extended Materials and Methods for details). The rhodesain plasmid and the GFP gene were amplified in a PCR using following primers 5’-caccaccatcaccaccactaagg-3’ and 5’-gctaccttgaaaatacagattctccggg-3’ for the vector and 5’-ctgtattttcaaggtagcgtgagcaagggcgaggagc-3’as well as 5’-ggtggtgatggtggtgcttgtacagctcgtccatgccg-3’ for the insert. The plasmid was transformed into Rosetta 2(DE3)pLysS cells (Novagen).

### *Purification of pro-rhodesain C150A from* E. coli

The active site C150A mutation was introduced by site direct mutagenesis using the primers 5’-GCGGTAGCGCGTGGGCGTTCAGCACC-3’ and 5’-CGCCCACGCGCTACCGCATTGGCCCTG-3’. Transformation, protein expression, cell lysis, purification by Ni-NTA and TEV-cleavage were performed as in extended Materials and Methods. In short, upon digestion of IMAC-purified pro-rhodesain with TEV protease, the cleavage products were loaded on 8 mL Ni-NTA gravity flow column preincubated with 50 mM Tris buffer (pH 8.5, 100 mM NaCl, 10 mM imidazole) to remove the TEV protease and GFP. Pro-rhodesain was eluted with 50 mM Tris buffer (pH 8.5, 100 mM NaCl, 10 mM imidazole) and combined with the flow through. After concentration by spin filtration to 4-5 mL pro-rhodesain was loaded onto a HiLoad 16/600 Superdex 75 pg column (GE Healthcare), and purified by SEC (0.5 mL/min, 20 mM Tris, pH 8.0, 200 mM NaCl). The protein was dialyzed against H2O at 4 °C, lyophilized and stored at −80 °C upon further usage.

### Circular Dichroism spectroscopy

CD spectra were measured using a Jasco-815 CD-spectrometer between 190 — 260 nm wavelength with 5 nm bandwidth, an interval of 1 nm and a scanning speed of 50 nm/min. Measurements of mature rhodesain from *P. pastoris* and *E. coli* (5 μM in 3 mM sodium phosphate buffer, pH 6.8, 0.5 mM TCEP, 5 mM NaCl) and inactive pro-rhodesain C150A (5 μM in 10 mM Tris, pH 8.0, 3 mM NaCl) were performed in a 1 mm quartz curvet at 25 °C. Spectra were derived from the average of three measurements after subtraction of the baseline.

The obtained ellipticities *θ* were converted to the mean residue ellipticities [*θ*]*_mrw,λ_* by the equation

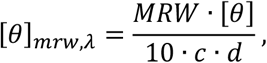

where *c* equals the concentration of the protein, *d* the pathlength through the sample and *MRW* for the mean residue weight, which is calculated by

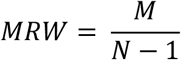

*M* corresponds to the molecular weight and *N* to the number of amnio acids in the protein.

### Circular Dichroism Spectroscopy of Pro-domain CathL Peptides

Peptides were purchased from *peptides&elefants* (Hennigsdorf, Germany). CD spectra were measured between 190 and 250 nm in sodium acetate (3 mM NaOAc, pH 4.0, 5 mM NaCl) or Tris buffer (3 mM Tris, pH 8.0, 5 mM NaCl) with additional TFE (0, 10, 30, 60 and 90% (v/v)) at a peptide concentration of 25 μM. Instrument settings, parameters and data transformations were performed equally to the whole protein CD-measurements as described above. Proportions of secondary structures were calculated by the online tool BeStSel (http://bestsel.elte.hu/index.php).

### Size Exclusion Chromatography

SEC runs were performed with a HiLoad 16/600 Superdex 75 pg column (GE Healthcare) at a flow rate of 0.5 mL/min at 4 °C. 4.5 mL samples were loaded on a 5 mL loop and analyzed using a 20 mM Tris (pH 8.0, 200 mM NaCl) as running buffer. Absorption was measured at a wavelength of 280 nm and values were normalized to the absorption maximum.

### Pro-rhodesain Autocleavage Assay

Inactive pro-rhodesain C150A was dissolved in the respective buffers: (a) 50 mM NaOAc pH 4.0, 235 mM NaCl, 5 mM EDTA, 5 mM DTT, 0.005% Brij35; (b) 50 mM NaOAc pH 5.5, 200 mM NaCl, 5 mM EDTA, 5 mM DTT, 0.005% Brij35; (c) 50 mM Bis-Tris pH 6.5, 229 mM NaCl, 5 mM EDTA, 5 mM DTT, 0.005% Brij35 or (d) 50 mM Tris pH 8.0, 221 mM NaCl, 5 mM EDTA, 5 mM DTT, 0.005% Brij35) to a concentration of 0.5 mg/mL. Mature, active rhodesain was pre-incubated in 50 mM sodium acetate buffer (pH 5.5, 200 mM NaCl, 5 mM EDTA, 5 mM DTT, 33 μg/mL) at room temperature for 10 min. 11 μL of the active rhodesain solution were added to 100 μL of the different pro-rhodesain solutions each and the mixture was kept at 37 °C. Samples were taken at the given time points, denatured immediately and analyzed by 15% SDS-PAGE. Band intensities were determined by integration with ImageJ (Version 1.52q) and normalized to the sample of the wild type protease at pH 5.5.

### X-ray crystallography

For crystallization, 1 μL of reservoir buffer (40 mM sodium citrate pH 3.5, 30% PEG-6000 (v/v), Pb(OAc)_2 (saturated)_) was added to 1 μL of a pro-rhodesain C150A solution (0,7 mg/mL in 10 mM sodium citrate, pH 5). Crystals were grown at room temperature by the hanging drop method within 2 weeks. The crystals were measured under cryo conditions. Thus, the crystals were preincubated in reservoir buffer with additional 10% (v/v) glycerol for 20 s before flash freezing in a 100 K N_2_ cryo stream. Diffraction data were collected with a Bruker AXS Microstar-H generator and a MAR-scanner 345 mm image-plate detector in a distance of 350 mm and a X-ray wavelength of 1.5417 Å in 1°-steps after 10 min exposition. Data processing was performed by the program XDS in space group H1.^[54]^ For structure determination, the previously determined structure of mature rhodesain in complex with a macrolactam inhibitor (PDB: 6EX8) was used as a model for molecular replacement with PhaserMR of the phenix work suites after removal of the ligand and all water molecules.^[24,54–56]^ The prodomain was then built into the remaining electron density and final refinement was done with WinCoot.^[57]^

### MD simulations

MD simulations were performed with the crystal structure of pro-rhodesain described in this manuscript but containing an active site cysteine residue instead of alanine. All amino acids and crystallographic water molecules were kept, while one glycerol molecule was removed from the structure. Missing residues within the structure were added and the structures were built using tleap of AmberTools17.^[58,59]^ For MDs resembling pH of 8, aspartate and glutamate residues were deprotonated and the catalytic diad was defined to form an ion pair of C150^-^ (thiolate, CYM) and H287^+^ (imidazolium, HIP). Additionally, residie H231 was protonated and H240 deprotonated (HIE). Protonation states of pH 4 were determined using predicted pK_a_-values from MOE, resulting in protonation of all histidine residues, as well as protonated D143, 185, 194, 271, 289, 296 (ASH) and E38, 58, 66, 95, 144, 175, 232, 256, 283, 322 and 343 (GLH). Again, the catalytic C150 was deprotonated to form the active ion pair. The protein structures were energetically minimized with sander over 200 time steps. Counter ions (7 Cl^-^ for pH 4 and 10 Na^+^ for pH 8) to neutralize the system and a TIP3P waterbox exceeding the protein by 10 Å in every dimension were added. System equilibration was performed over 500 ns with gradually releasing constraints and 500 ns without constraint while heating from 100 to 300 K in an NVT ensemble.^[60]^ Subsequent production runs were performed in an NPT ensemble with periodic boundary conditions and a van der Waals cutoff of 14.0 Å over 20 ns. All simulations were performed with the AMBER forcefield (ff14SB) on the high performance computing (HPC) cluster MOGON of the University of Mainz with NAMD-2.12.^[61,62]^ Simulations were performed 4 times for each pH and results were analyzed using VMD-1.9.3 and related scripts.^[63]^ For the analysis of hydrogen bonds a cut-off distance of 3.0 Å and angle of 20° were selected.

### Z-Phe-Arg-AMC docking

The predicted binding mode of the assay substrate in complex with rhodesain (Fig. 5) was generated by molecular docking. The receptor was prepared within LeadIT-2.3.2 (*BioSolveIT GmbH*, Sankt Augustin, Germany, 2017, www.biosolveit.de/LeadIT) using the crystal structure of the K11777-rhodesain complex, PDB-ID 2P7U.^[25,64]^ The covalent ligand was untethered from the enzyme using MOE (Molecular Operating Environment, 2018.01; *Chemical Computing Group ULC*: Montreal, QC, Canada, 2018). For docking the receptor was protonated using the Protoss module within LeadIT and the catalytic dyad was manually set to form the C25 thiolate and H162 imidazolium ion pair (corresponding to C150 and H287 in the pro-enzyme).^[65]^ Water molecule 512 was kept as part of the binding site. Ligands were energetically minimized using omega2 (OMEGA 3.1.0.3, *OpenEye Scientific Software*, Santa Fe, NM. http://www.eyesopen.com) and the Merck molecular force field (MMFF94).^[66,67]^ The receptor was validated by redocking of K11777 (FlexX score −24.92 kJ/mol, RMSD of 1.7 Å). The substrate ligand Z-Phe-Arg-AMC was subsequently docked into the binding site.

### PET fluorescence experiments

Steady-state fluorescence emission spectra were recorded using a Jasco FP-6500 spectrofluorometer. Sample temperature was set to 37 °C using a Peltier thermocouple. A sample concentration of 100 nM modified pro-rhodesain in a 0.5 ml quartz glass cuvette (Hellma) was used. Spectra were recorded in either 50 mM phosphate, pH 8.0, with the ionic strength adjusted to 200 mM using sodium chloride, or in 50 mM acetate, pH 5.5, with the ionic strength adjusted to 200 mM using sodium chloride. Both buffers contained 5 mM EDTA; and 0.3 mg/mL bovine serum albumine and 0.05% Tween-20 to prevent sticking of rhodesain to the glass surface.

PET-FCS was performed on a custom-built confocal fluorescence microscope setup that consists of an inverse microscope body (Zeiss Axiovert 100 TV) equipped with a diode laser emitting at 637 nm as excitation source (Coherent Cube). The laser beam is coupled into an oil-immersion objective lens (Zeiss Plan Apochromat, 63x, NA 1.4) via a dichroic beam splitter (Omega Optics 645DLRP). The average laser power was adjusted to 400 μW before entering the back aperture of the microscope using an optical density filter. The fluorescence signal was collected by the same objective, filtered by a band-pass filter (Omega Optics 675RDF50), and imaged onto the active area of two fibre-coupled avalanche photodiode detectors (APDs; Perkin Elmer, SPCM-AQRH-15-FC) using a cubic non-polarizing beam-splitter (Linos) and multi-mode optical fibres of 100 μm diameter. The signals of the APDs were recorded in the cross-correlation mode using a digital hard-ware correlator device (ALV 5000/60X0 multiple tau digital real correlator) to bypass detector dead-time and after-pulsing effects. Fluorescently modified pro-rhodesain was diluted to 1 nM samples in the buffered solutions specified above for steady-state fluorescence spectroscopy. Samples were filtered through a 0.2 μm syringe filter before measurement, transferred onto a microscope slide and covered by a cover slip. Sample temperature was set to 37 °C using a custom-built objective heater. For each sample the accumulated measurement time of the autocorrelation function, G(τ), was 15 min. G(τ) was fitted using an analytical model for translational diffusion of a molecule that exhibits chemical relaxations.^[27]^ ACFs fitted well to a model for diffusion of a globule with either one- or two single-exponential decays:

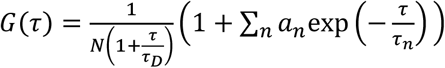

*τ* is the lag time, *N* is the average number of molecules in the detection focus, *τ*_D_ is the diffusion time constant, *a*_n_ is the amplitude of the *n*th relaxation, and *τ*_n_ is the according time constants. The application of a model for diffusion in two dimensions was of sufficient accuracy because the two horizontal dimensions (*x, y*) of the detection focus were much smaller than the lateral dimension (*z*) in the applied setup.^[27]^ Errors are s.e. from regression analyses.

## Acknowledgements

We thank Prof. Hermann Schindelin, Universität Würzburg, for the generous gift of the *P. pastoris* cells expressing rhodesain. We gratefully acknowledge the advisory services offered and the computing time granted on the supercomputer Mogon at Johannes Gutenberg University Mainz (hpc.uni-mainz.de), which is a member of the AHRP (Alliance for High Performance Computing in Rhineland Palatinate, www.ahrp.info) and the Gauss Alliance e.V. We further thank Openeye Scientific for free academic software licenses. Research was supported by the Carl-Zeiss Foundation (to UAH) and the Centre for Biomolecular Magnetic Resonance (BMRZ), Goethe-University Frankfurt, funded by the state of Hesse.

The X-ray structure of *Trypanosoma brucei rhodesiense* pro-rhodesain C150A has been deposited in the PDB under the accession number 7AVM.

## Author contributions

JP cloned constructs, established *E. coli* expression and purification protocols, purified proteins and carried out CD spectroscopy and SEC; JP, EJ and CK carried out X-ray crystallography; CK carried out MD simulations; JP and HN carried out PET-FCS measurements; JP, HN, TS CK and UAH analyzed data; JP, TS and UAH designed the project; TS and UAH supervised the project; JP, CK and UAH wrote the paper with contributions from all authors.

