## Supporting Information for "Structure, interdomain dynamics and pH-dependent autoactivation of pro-rhodesain, the main lysosomal cysteine protease from African trypanosomes"

### *Extended Materials and Methods*

### *Optimization of purification of pro- and mature rhodesain from E. coli*

Initially, the pro-rhodesain construct used for heterologous expression in *E. coli* contained the N-terminal propeptide, the catalytic domain, a tobacco etch virus (TEV)-protease cleavage sequence and a C-terminal 6xHis tag (Fig. S1A). This construct differs from the wildtype protein by the lack of the first 20 amino acids constituting the signal peptide and a C-terminal domain of unknown function. While protein expression could be induced by IPTG, multiple bands were observed on SDS-PAGE in the expected size-range for (pro-)rhodesain (~37/~24 kDa) (Fig. S1B). Although western blot analysis with anti-$His_6$-HPR revealed that the 6xHis-tag remained attached to the protein, no interaction with a $Ni^{2+}$-NTA matrix could be achieved (Fig. S1C, D).

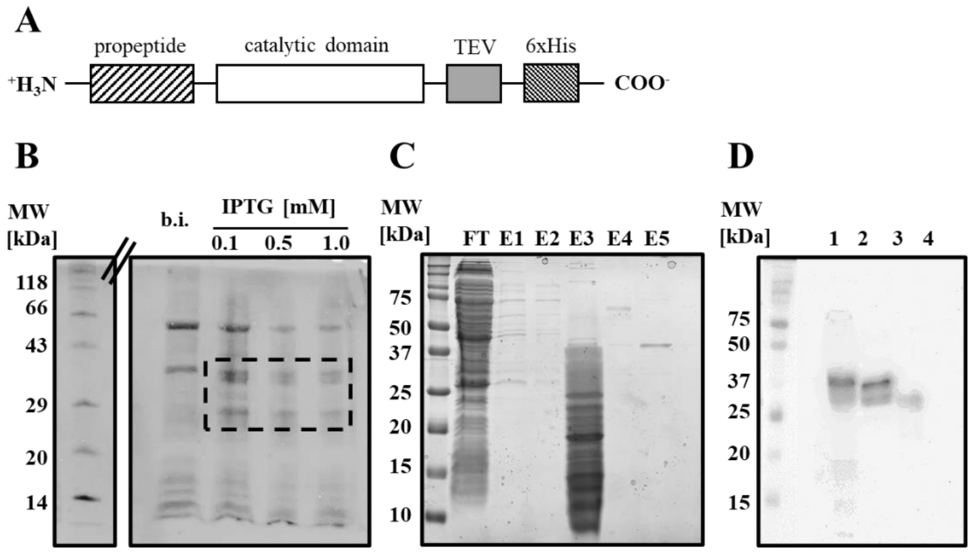

**Figure S1:** Expression and purification optimization of *T. b. rhodesiense* rhodesain from *E. coli*. A) Initial pro-rhodesain construct design. B) 15% SDS-PAGE of whole *E. coli* cell lysate before (b.i.) and after induction with various IPTG concentrations. C) 15% SDS-PAGE of flow-through (FT) and different $Ni^{2+}$-NTA elution fractions when trying to purify pro-rhodesain via IMAC. D) Western blot with anti-$His_6$-antibody of the soluble fraction of *E. coli* cells shows that the His-tag was attached to the protein. (Cells were lysed in the presence of different additives (1: lysis buffer; 2: lysis buffer + 0.5% (v/v) TritonX; 3: lysis buffer + 1 M urea; 4: lysis buffer cells without IPTG induction).

To tackle this issue, we incorporated a green fluorescent protein (GFP) between the TEV cleavage sequence and the 6xHis-tag to increase the distance between the tag and the protein, to improve solubility and to be able to follow the purification progress more easily (Fig. S2A). The DNA and peptide sequences of the rhodesain-GFP construct are shown in Figure S3. Indeed, insertion of the GFP led to a construct that was efficiently expressed and could be purified with $Ni^{2+}$-NTA beads (Fig. S2B, see below for details). TEV protease could efficiently cleave off the GFP-His6 tag, however and unexpectedly, the separation of pro-rhodesain and the GFP by an additional $Ni^{2+}$-NTA based IMAC purification step failed because both fragments were found in the flow through. For the inactive pro-rhodesain C150A constructs, this was not the case and these constructs could be purified efficiently by combining an IMAC purification step with a TEV digest, a second "reverse" IMAC and a SEC step (see main paper for details). This indicates that loss of the GFP-6xHis fragment's ability to interact with the $Ni^{2+}$-NTA matrix may be related to an inherent, weak rhodesain activity.

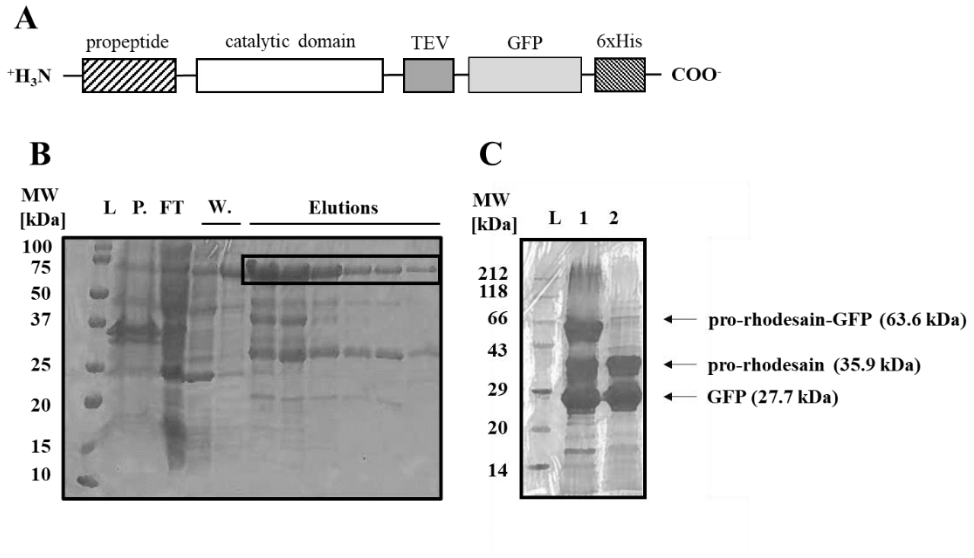

**Figure S2:** Insertion of GFP in the (pro-)rhodesain construct enables purification by immobilized metal affinity chromatography (IMAC). A) Construct design. B) 15% SDS-PAGE of different elution fraction from the purification of pro-rhodesain-GFP via IMAC. Protein marker (L.), insoluble fraction (P.), flow through (FT), wash fraction (at 20 mM imidazole, W.) and elution fractions (at 100 mM imidazole) are marked. The bands of the pro-rhodesain-GFP fusion protein are boxed. C) 15% SDS-PAGE of the $Ni^{2+}$-NTA elution fraction before (1) and after (2) cleavage by TEV protease.

Furthermore, because the GFP-His6-tag and the protease could not be separated efficiently by IEX or SEC after TEV cleavage, the mature wildtype protease had to be purified taking advantage of its auto-activation behavior. A drop in pH was used to activate the autocleavage of the enzyme which simultaneously led to the digestion as well as the slow denaturation of the released GFP and thereby to the vanishing of the green color. The precipitated GFP was removed by centrifugation. Although the rhodesain activation step contributed to purification of the protease itself, it needed to be carefully controlled, due to the occurrence of autoproteolysis products which occur after the enzyme reaches a short plateau of maximum catalytic activity (Fig. S4A, B). As described in the main paper, autoproteolysis can be prevented via the reversible inhibition of the enzyme with PMSF. Once the propeptide was removed, the released mature rhodesain could be purified from the remaining peptide fragments by SEC. Alternatively final purification could also be achieved by anion exchange chromatography, due to the very low pI value of 3.94 for mature rhodesain.

***Detailed expression and purification protocol of pro-rhodesain:*** Cells transformed with the plasmid encoding the GFP-tagged pro-rhodesain were grown at 37 °C in 100 mL LB medium in the presence of 100 µg/mL ampicillin and 50 µg/mL chloramphenicol overnight. 3 L main culture (LB medium, 100 µg/mL ampicillin, 50 µg/mL chloramphenicol) were inoculated with 2 Vol% of pre-culture and incubated at 37 °C. When the culture reached an $OD_{600}$ of ~ 0.5 after 5 h, IPTG was added to a final

ATGGCTTGTCTAGCATCAGTAGCTCTAGGGAGTTTACACGTTGAAGAATCACTAGAAATGCGTTTT

M A C L A S V A L G S L H V E E S L E M R F

GCGGCGTTCAAGAAGAAATACGGTAAGGTGTACAAAGACGCGAAGGAAGAGGCGTTCCGTTTTCGT

A A F K K K Y G K V Y K D A K E E A F R F R

GCGTTCGAGGAAAACATGGAGCAAGCGAAAATCCAAGCGGCGGCGAACCCGTATGCGACCTTCGGC

A F E E N M E Q A K I Q A A A N P Y A T F G

GTTACCCCGTTTAGCGATATGACCCGTGAGGAATTCCGTGCGCGTTACCGTAACGGTGCGAGCTAT

V T P F S D M T R E E F R A R Y R N G A S Y

TTTGCGGCGGCGCAAAAACGTCTGCGTAAGACCGTGAACGTTACCACCGGTCGTGCGCCGGCGGCG

F A A A Q K R L R K T V N V T T G R A P A A

GTGGACTGGCGTGAAAAGGGTGCGGTGACCCCGGTTAAGGATCAGGGCCAATGCGGTAGC**TGC**TGG

V D W R E K G A V T P V K D Q G Q C G S **C** W

GCGTTCAGCACCATCGGCAACATTGAGGGCCAGTGGCAAGTGGCGGGCAACCCGCTGGTTAGCCTG

A F S T I G N I E G Q W Q V A G N P L V S L

AGCGAACAGATGCTGGTGAGCTGCGACACCATCGATTTCGGTTGCGGTGGCGGTCTGATGGACAAC

S E Q M L V S C D T I D F G C G G G L M D N

GCGTTTAACTGGATTGTGAACAGCAACGGCGGTAACGTTTTTACCGAGGCGAGCTACCCGTATGTT

A F N W I V N S N G G N V F T E A S Y P Y V

AGCGGCAACGGCGAGCAGCCGCAATGCCAGATGAACGGCCACGAAATCGGTGCGGCGATTACCGAC

S G N G E Q P Q C Q M N G H E I G A A I T D

CACGTGGATCTGCCGCAAGACGAGGATGCGATTGCGGCGTACCTGGCGGAAAACGGTCCGCTGGCG

H V D L P Q D E D A I A A Y L A E N G P L A

ATTGCGGTTGATGCGACCAGCTTTATGGATTATAACGGCGGTATTCTGACCAGCTGCACCAGCGAA

I A V D A T S F M D Y N G G I L T S C T S E

CAGCTGGACCACGGCGTGCTGCTGGTTGGTTACAACGATGCGAGCAACCCGCCGTATTGGATCATT

Q L D H G V L L V G Y N D A S N P P Y W I I

AAAAACAGCTGGAGCAACATGTGGGGCGAGGATGGTTACATCCGTATTGAAAAGGGCACCAACCAA

K N S W S N M W G E D G Y I R I E K G T N Q

TGCCTGATGAACCAGGCGGTTAGCAGCGCGGTTGTTGGCGGCCCGGAGAATCTGTATTTTCAAGGT

C L M N Q A V S S A V V G G P *E N L Y F Q* ǁ*G*

AGCGTGAGCAAGGGCGAGGAGCTGTTCACCGGGGTGGTGCCCATCCTGGTCGAGCTGGACGGCGAC

S V S K G E E L F T G V V P I L V E L D G D

GTAAACGGCCACAAGTTCAGCGTGTCCGGCGAGGGCGAGGGCGATGCCACCTACGGCAAGCTGACC

V N G H K F S V S G E G E G D A T Y G K L T

CTGAAGTTCATCTGCACCACCGGCAAGCTGCCCGTGCCCTGGCCCACCCTCGTGACCACCCTGACC

L K F I C T T G K L P V P W P T L V T T L T

TACGGCGTGCAGTGCTTCAGCCGCTACCCCGACCACATGAAGCAGCACGACTTCTTCAAGTCCGCC

Y G V Q C F S R Y P D H M K Q H D F F K S A

ATGCCCGAAGGCTACGTCCAGGAGCGCACCATCTTCTTCAAGGACGACGGCAACTACAAGACCCGC

M P E G Y V Q E R T I F F K D D G N Y K T R

GCCGAGGTGAAGTTCGAGGGCGACACCCTGGTGAACCGCATCGAGCTGAAGGGCATCGACTTCAAG

A E V K F E G D T L V N R I E L K G I D F K

GAGGACGGCAACATCCTGGGGCACAAGCTGGAGTACAACTACAACAGCCACAACGTCTATATCATG

E D G N I L G H K L E Y N Y N S H N V Y I M

GCCGACAAGCAGAAGAACGGCATCAAGGTGAACTTCAAGATCCGCCACAACATCGAGGACGGCAGC

A D K Q K N G I K V N F K I R H N I E D G S

GTGCAGCTCGCCGACCACTACCAGCAGAACACCCCCATCGGCGACGGCCCCGTGCTGCTGCCCGAC

V Q L A D H Y Q Q N T P I G D G P V L L P D

AACCACTACCTGAGCACCCAGTCCGCCCTGAGCAAAGACCCCAACGAGAAGCGCGATCACATGGTC

N H Y L S T Q S A L S K D P N E K R D H M V

CTGCTGGAGTTCGTGACCGCCGCCGGGATCACTCTCGGCATGGACGAGCTGTACAAGCACCACCAT

L L E F V T A A G I T L G M D E L Y K H H H

CACCACCACTAA

H H H ***

**Figure S3:** DNA and peptide sequences of the rhodesain-GFP construct. The stop codon (***), propeptide (underlined), C150A mutation (underlined), TEV cleavage site (ǁ) and GFP (underlined) are highlighted.

concentration of 0.5 mM for induction. The cells were cultivated at 19.5 °C for 18 h, harvested by centrifugation at 15,000 *g* at 4 °C and stored at -80 °C.

The cell pellet was resuspended in 100 mL lysis buffer containing 50 mM Tris pH 8.5, 100 mM NaCl, 10 mM imidazole and 1 mM benzamidine. The cells were incubated for 15 min on ice in presence of lysozyme and DNase I. After sonication on ice, the non-soluble cell fragments were sedimented by centrifugation at 25,000 *g* for 30 min at 4 °C and the green supernatant was loaded onto a Ni-NTA column with gravity flow. After a first washing step with 20 mM imidazole the green rhodesain-GFP construct was eluted at 100 mM imidazol concentration. 10% TEV-protease (n/n) was added to the combined fractions containing the desired protein to cleave the cysteine protease from its fusion tag. The solution was dialyzed against 50 mM Tris pH 8, 0.5 mM EDTA, 3 mM Glutathion red., 0.3 mM Glutathion ox., 10 mM NaCl over night at 4 °C. The TEV-protease was removed by anion exchange chromatography (HiTrap Q 5 mL, Buffer A: 50 mM Tris pH 8 Buffer B: 50 mM Tris pH 8, 500 mM NaCl). However, a separation of pro-rhodesain and GFP could not be achieved at this step. The maturation of the rhodesain zymogen was induced by a pH drop. DTT was added to a final concentration of 2 mM prior to adding 200 mM citrate buffer (pH 3) dropwise until a final pH value of 3.5 was reached. The enzyme was incubated at room temperature and its activity monitored by a fluorometric enzyme assay until a maximum of activity was observed. DTT was eliminated using KTT (5 mM final concentration) and the enzyme was inhibited by adding 2 mM PMSF to prevent autohydrolysis. The resulting pro-rhodesain was concentrated by spin filtration and further purified by SEC (HiLoad 16/600 Superdex 75 pg column (GE Healthcare), 0.5 mL/min, 20 mM sodium citrate, pH 5.0, 200 mM NaCl). The protein was dialyzed against $H_2O$ at 4 °C for 4 h, lyophilized and stored at -80 °C upon further use.

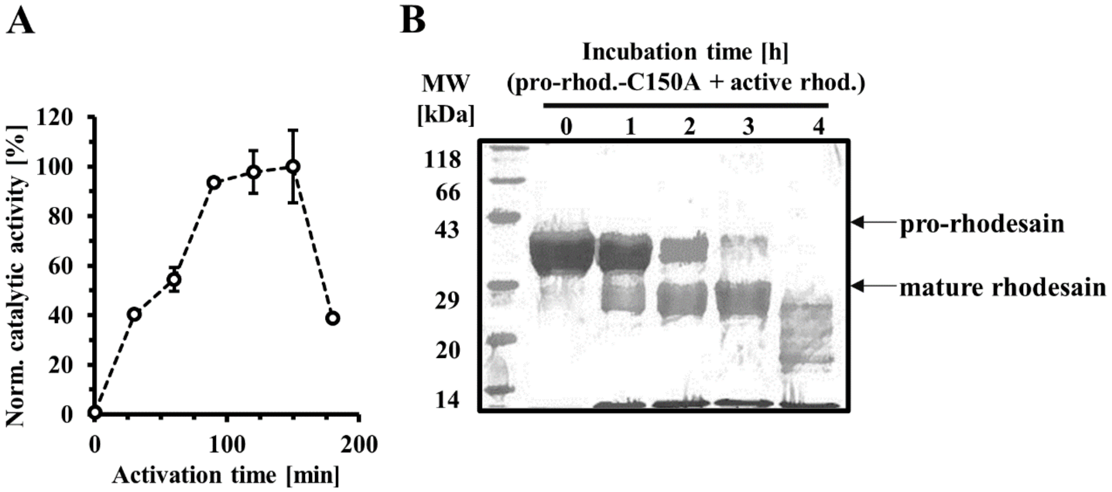

**Figure S4:** Autoactivation of rhodesain can lead to an unspecific autoproteolysis and an irreversible destruction of the enzyme. A) Autoactivation of pro-rhodesain was induced by a pH drop. Samples were taken at given time points and the catalytic activity determined via a fluorescence based enzyme assay with the peptidic substrate Z-Phe-Arg-AMC. The slope of the increasing fluorescence over time (which corresponds to the catalytic activity) was plotted against the pre-incubation time in the acidic milieu. B) SDS-PAGE of inactive pro-rhodesain C150A with active mature rhodesain shows complete digestion of enzymes at prolonged time points.

***Site directed mutagenesis***

Site directed mutagenesis was performed by quik-change PCR with the KAPA-polymerase kit (KAPA HiFi HotStart PCR Kit, *Roche*, Mannheim, Germany). Primers were purchased from *Biomers* (Ulm, Deutschland) or IDT (*Integrated DNA Technologies*, Iowa, USA). The following primers were used to introduce point mutations into pro-rhodesain:

Q146W: 5´-CAGGGCTGGTGCGGTAGCTGCTGGGCGTTCAG-3´, 5´-CCGCACCAGCCCTGATCCTTAAC-CGGGGTCAC-3´; C150A: 5´-GCGGTAGCGCGTGGGCGTTCAGCACC-3´, 5'-CGCCCACGCGCTACCG-CATTGGCCCTG-3'; C22S: 5´-GGCTAGCCTAGCATCAGTAGCTCTAGGGAG-3´, 5´-CTGATGCTAGG-CTAGCCATATGTATATCTCCTTC-3´; A79C: 5´-GAATTCCGTTGCCGTTACCGTAACGGTGCG-3´, 5´-GTAACGGCAACGGAATTCCTCACGGGTCATATCGC-3´; V51C: 5´-CGGTAAGTGCTACAAAGA-CGCGAAGGAAGAGG-3´, 5´-CTTTGTAGCACTTACCGTATTTCTTCTTGAACGCCG-3´; D194N: 5´-CGGTCTGATGAACAACGCGTTTAACTGG-3´, 5´-GTTGTTCATCAGACCGCCACCGCAAC-3´; D242N: 5´-CCACGTGAACCTGCCGCAAGACGAGGATGC-3´, 5´-CGGCAGTTGCACGTGGTCGGTA-ATCGCCG-3´; Q146W (in C150A background): 5´-GATCAGGGCTGGTGCGGTAGCTGCTGGG-3´ and 5´-CGCACCAGCC-CTGATCCTTAACCGGGG-3´.

***Fluorescence based cleavage assay***

Catalytic activity of rhodesain was measured at a fluorimeter F2000 (Tecan) by detecting the hydrolysis of the fluorogenic substrate Z-Phe-Arg-AMC (Bachem). The enzyme (90 ng/mL) was incubated in 195 µL assay buffer (50 mM NaOAc pH 5.5, 200 mM NaCl, 5 mM EDTA, 5 mM DTT, 0.005% Brij35 (w/w)) at room temperature for 10 min before the reaction was started by adding 5 µL of substrate (varying concentrations in DMSO). The reaction was kept at 25 °C and the fluorescence of the released AMC was measured at a wavelength of 460 nm after excitation at a wavelength of 380 nm. Since the fluorescence intensity is proportional to the released AMC, the catalytic activity was derived from the slope of the line. Enzyme kinetic values $K_M$ and $v_{max}$ were calculated by GraFit® (5.0.13, Erithacus Software) and the fitting equation

$$v = \frac{v_{max}\,[S]}{K_M + [S]}$$

were v equals the catalytic activity at a given substrate concentration [S] and $K_M$ is the Michaelis-Menten constant.

Comparison of the catalytic activities of mature rhodesain purified from *E. coli* or *P. pastoris* was probed using the fluorescence assay described above using the peptide Z-Phe-Arg-AMC (Bachem). The resulting $K_M$ and $v_{max}$ values for rhodesain expressed in *P. pastoris* are 5.78 ± 0.5 µM and 6.88 ± 0.7 $s^{-1}$ and agree with those previously published[1,2]. Likewise, the values obtained for rhodesain purified from *E. coli* are 4.62 ± 0.42 µM and 5.47 ± 0.45 $s^{-1}$ and thus show that the different expression host does not affect rhodesain functional integrity.

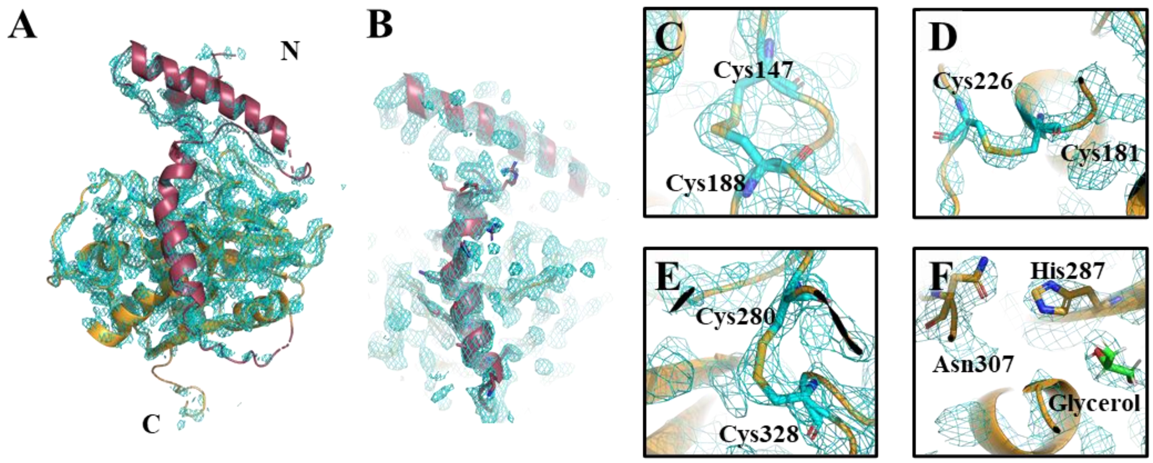

**Figure S5:** Electron density map of the pro-rhodesain crystal structure. The electron density map for the whole structure (A) and the pro-peptide (B) is colored in cyan at a sigma level of 2.0. The zoom-ins show the regions of the three disulfide bridges (C-E) and the active site (F).

### *Alignment of CathL zymogenes from different species*

```
Rhodesain       -----------ACLASVALGSLHVEESLEMRFAAFKKKYGKVYKDAKEEAFRFRAFEENM    69
Cruzain         ------------CLVPAATASLHAEETLTSQFAEFKQKHGRVYESAAEEAFRLSVFRENL    66
hsCathL         --------------------TLTFDHSLEAQWTKWKAMHNRLYGM-NEEGWRRAVWEKNM    56
Caricain        LFVHMSVSFGDFSIVGYSQDDLTSTERLIQLFNSWMLNHNKFYENVDEKLYRFEIFKDNL    76
Papain          --VYMGLSFGDFSIVGYSQNDLTSTERLIQLFESWMLKHNKIYKNIDEKIYRFEIFKDNL    76
                                     *   . *   :  :   :.:.*    *: :*   :..*:

                                                    blocking peptide
Rhodesain       EQAKIQAAAN----PYATFGVTPFSDMTREEFRARYRNGASYFA-AAQKRLRKTVNVTTG    124
Cruzain         FLARLHAAAN----PHATFGVTPFSDLTREEFRSRYHNGAAHFA-AAQERARVPVKVEVV    121
hsCathL         KMIELHNQEYREGKHSFTMAMNAFGDMTSEEFRQVMNGFQNRK----PRKGKVFQEPLFY    112
Caricain        NYIDETNKKN----NSYWLGLNEFADLSNDEFNEKYVGSLIDATIE-QSYDEEFINEDTV    131
Papain          KYIDETNKKN----NSYWLGLNVFADMSNDEFKEKYTGSIAGNYTTTELSYEEVLNDGDV    132
                                  :.:. *.*:: :**.    .             .   :

Rhodesain       RAPAAVDWREKGAVTPVKDQGQCGSCWAFSTIGNIEGQWQVAGNPLVSLSEQMLVSCDT-    183
Cruzain         GAPAAVDWRARGAVTAVKDQGQCGSCWAFSAIGNVECQWFLAGHPLTNLSEQMLVSCDK-    180
hsCathL         EAPRSVDWREKGYVTPVKNQGQCGSCWAFSATGALEGQMFRKTGRLISLSEQNLVDCSGP    172
Caricain        NLPENVDWRKKGAVTPVRHQGSCGSCWAFSAVATVEGINKIRTGKLVELSEQELVDCER-    190
Papain          NIPEYVDWRQKGAVTPVKNQGSCGSCWAFSAVVTIEGIIKIRTGNLNEYSEQELLDCDR-    191
                  *  **** :* ** *:.**.********:   :*         * . *** *:.*.

Rhodesain       -IDFGCGGGLMDNAFNWIVNSNGGNVFTEASYPYVSGNGEQPQCQMNGHEIGAAITDHVD    242
Cruzain         -TDSGCSGGLMNNAFEWIVQENNGAVYTEDSYPYASGEGISPPCTTSGHTVGATITGHVE    239
hsCathL         QGNEGCNGGLMDYAFQYVQ--DNGGLDSEESYPYEATE---ESCKYNPKYSVANDTGFVD    227
Caricain        -RSHGCKGGYPPYALEYVAK---NGIHLRSKYPYKAKQGTCRAKQVGGPIVKTSGVG-RV    245
Papain          -RSYGCNGGYPWSALQLVAQ---YGIHYRNTYPYEGVQRYCRSREKGPYAAKTDGVR-QV    246
                  . ** **    *:: :       :  . .*** . :        .     :  .

Rhodesain       LPQDEDAIAAYLAENGPLAIAVDA--TSFMDYNGGIL--TSCTSEQLDHGVLLVGYN---    295
Cruzain         LPQDEAQIAAWLAVNGPVAVAVDA--SSWMTYTGGVM--TSCVSEQLDHGVLLVGYN---    292
hsCathL         IPKQEKALMKAVATVGPISVAIDAGHESFLFYKEGIYFEPDCSSEDMDHGVLVVGYGFES    287
Caricain        QPNNEGNLLN-AIAKQPVSVVVESKGRPFQLYKGGIFEGP--CGTKVDHAVTAVGYG---    299
Papain          QPYNEGALLY-SIANQPVSVVLEAAGKDFQLYRGGIFVGP--CGNKVDHAVAAVGYG---    298
                 * :*  :        *:::.:::    :  *  *:       . .:**.*  ***.

Rhodesain       -DSSNPPYWIIKNSWSNMWGEDGYIRIEKGT-NQ---CLMNQAVSSAVVGGPTPPPPPP-    349
Cruzain         -DSAAVPYWIIKNSWTTQWGEEGYIRIAKGS-NQ---CLVKEEASSAVVGGPGPTPEPTT    347
hsCathL         TESDNNKYWLVKNSWGEEWGMGGYVKMAKDRRNH---CGIASAASYPTV-----------    333
Caricain        -KSGGKGYILIKNSWGTAWGEKGYIRIKRAPGNSPGVCGLYKSSYYPTKN----------    348
Papain          -----PNYILIKNSWGTGWGENGYIRIKRGTGNSYGVCGLYTSSFYPVKN----------    345
                       * ::****   **  **::: :   *    *  :       .

Rhodesain       -------PPPSATFTQDFCEGKGCTKGCSHATFPTGECVQTTGVGSVIATCGASNLTQII    402
Cruzain         TTTTSAPGPSPSYFVQMSCTDAACIVGCENVTLPTGQCLLTTSGVSAIVTCGAETLTEEV    406
hsCathL         ------------------------------------------------------------    333
Caricain        ------------------------------------------------------------    348
Papain          ------------------------------------------------------------    345

Rhodesain       YPLSRSCSGLSVPITVPLDKCIPILIGSVEYHCSTNPPTKAARLVPHQ------------    450
Cruzain         FLTSTHCSGPSVRSSVPLNKCNRLLRGSVEFFCGSSSSGRLADVDRQRRHQPYHSRHRRL    467
hsCathL         ------------------------------------------------------------    333
Caricain        ------------------------------------------------------------    348
Papain          ------------------------------------------------------------    345
```

**Figure S6:** Alignment of cathepsin L-like proteases from different organisms. Amino acid sequences of *T. b. rhodesiense* pro-rhodesain (uniprot-ID.: Q95PM0), *T. cruzi* pro-cruzain (uniprot-ID.: P25779), human pro-cathL (uniprot-ID.: P07711), *C. papaya* caricain and papain (uniprot-IDs: P10056, P00784) without the N-terminal signal peptides were aligned using the Clustal Ω multiple sequence alignment tool (version 1.2.4). X: conserved aromatic amino acids, cyan: ER(I/V)FNIN/ ER(I/V)FNAA motif; yellow: GNFD motif, underlined: trypanosomal blocking peptide; green: β-sheet within the pro-peptide binding loop (PBL).

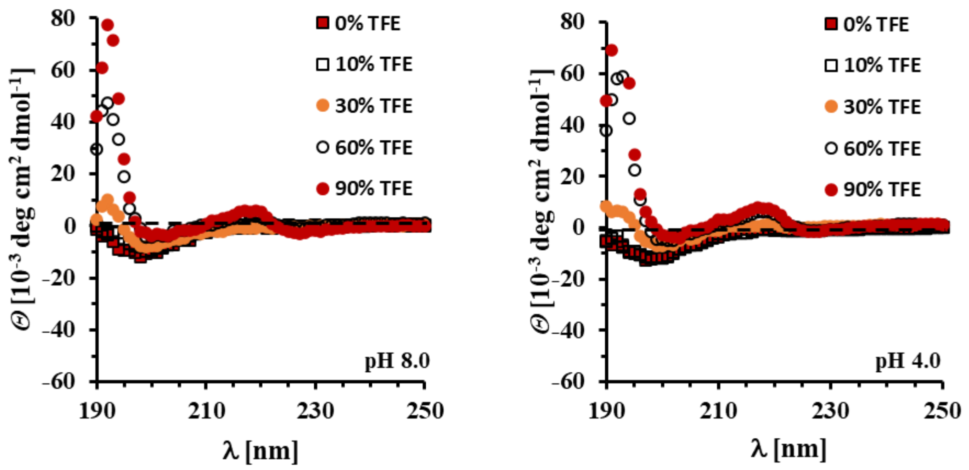

**Figure S7**: CD-spectra of the blocking peptide from human pro-CathL at various TFE concentrations and different pH values.

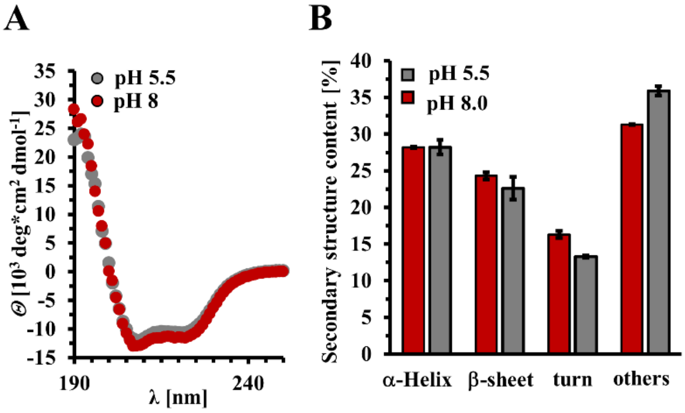

**Figure S8:** Comparison of secondary structure of pro-rhodesain at pH 8 and pH 5.5 (A) CD spectra of pro-rhodesain C150A at pH 8 and pH 5.5. (B) Calculated secondary structure content based on CD spectra using the BsStSel server.[3]

### *Molecular dynamics simulations*

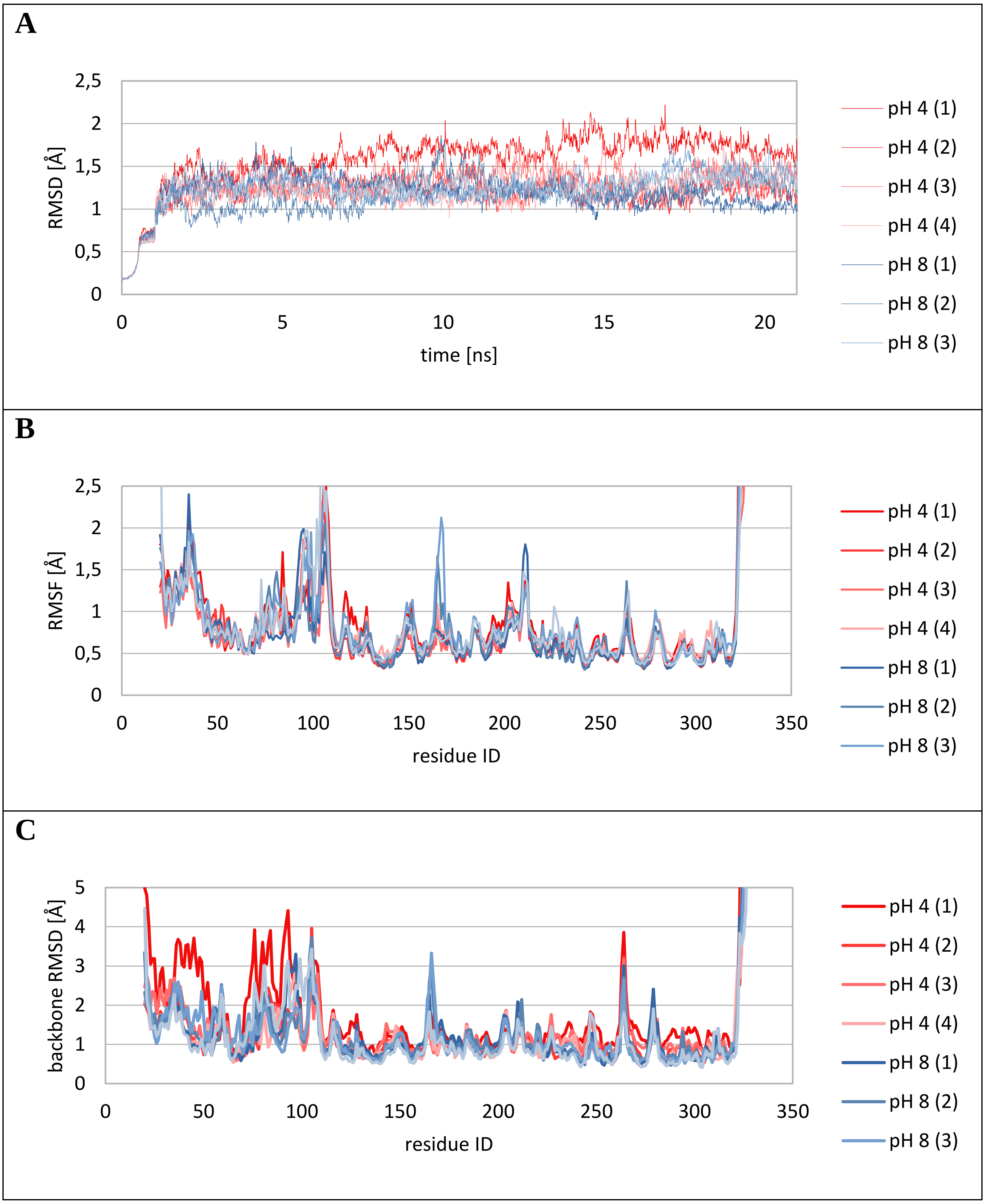

**Figure S9:** MD simulations on pro-rhodesain. A) 1-D RMSD traces of 8 MD simulations over 1 ns of equilibration and 20 ns production at pH 4 (red traces) and 8 (blue traces), respectively for backbone atoms excluding highly flexible C-terminal residues 338-348. B) $C_\alpha$-RMSF per residue over 20 ns MD production run. C) RMSD per residue backbone compared to starting structure.

**Table S1: Average intra- and interdomain electrostatic ($E_{ele}$) and van der Waals ($E_{vdW}$) energies from MD simulations** within the pro-domain (pro), within the main domain (main), between pro-domain and main domain (pro-main), between pro-domain and solvent (pro-solv) and main domain and solvent (main-solv) calculated from MD simulations at pH 4 and 8, respectively. All values are in kcal/mol.

| Simulation | $E_{ele}$ (pro-main) | $E_{vdW}$ (pro-main) | $E_{ele}$ (pro) | $E_{vdW}$ (pro) | $E_{ele}$ (main) | $E_{vdW}$ (main) | $E_{ele}$ (pro-solv) | $E_{vdW}$ (pro-solv) | $E_{ele}$ (main-solv) | $E_{vdW}$ (main-solv) |
|---|---|---|---|---|---|---|---|---|---|---|
| **pH 4 (1)** | -422.8 | -226.5 | -2697.9 | -209.6 | -5718.1 | -900.0 | -2703.8 | -237.3 | -3335.3 | -340.8 |
| **pH 4 (2)** | -359.4 | -198.0 | -2683.6 | -213.5 | -5701.7 | -918.2 | -2830.7 | -255.6 | -3482.1 | -342.4 |
| **pH 4 (3)** | -308.1 | -190.4 | -2654.3 | -213.8 | -5717.0 | -919.4 | -2915.3 | -261.7 | -3409.8 | -324.8 |
| **pH 4 (4)** | -264.5 | -188.0 | -2698.7 | -220.1 | -5654.9 | -917.5 | -2861.6 | -256.3 | -3715.7 | -346.5 |
| **pH 8 (1)** | -585.3 | -198.3 | -2731.4 | -205.0 | -5039.3 | -894.7 | -2933.7 | -238.8 | -4524.1 | -294.2 |
| **pH 8 (2)** | -706.4 | -181.7 | -2681.2 | -214.1 | -5042.2 | -901.4 | -2903.2 | -241.8 | -4605.8 | -285.3 |
| **pH 8 (3)** | -665.8 | -191.4 | -2811.4 | -201.7 | -5020.6 | -886.7 | -2676.1 | -246.1 | -4680.5 | -302.4 |
| **pH 8 (4)** | -678.2 | -196.8 | -2662.5 | -206.6 | -5092.5 | -900.6 | -2936.8 | -234.5 | -4563.2 | -283.9 |

**Table S2a: Relative occurrence of hydrogen bonds within the pro-domain.** Four respective MD simulations at pH 4 (pH4_1 and pH4_2, see Table S2b for pH4_3 and pH4_4) were carried out. `Main' refers to interactions of the backbone. 'Side' indicates interactions of side chains. Occurrences below 5% were left out for clarity.

| | | | | | | | | | | | | | |
|---|---|---|---|---|---|---|---|---|---|---|---|---|---|
| pH4_1 | | | | | | | pH4_2 | | | | | | |
| | | | | | | ’ | | | | | | | |
| donor | | | acceptor | | | occupancy | donor | | | acceptor | | | occupancy |
| TYR | 52 | Side | ASP | 91 | Side | 70,00% | ARG | 61 | Side | GLU | 96 | Side | 86,00% |
| ARG | 61 | Side | GLU | 96 | Side | 66,45% | TYR | 52 | Side | ASP | 91 | Side | 68,90% |
| GLN | 75 | Main | GLN | 71 | Main | 50,65% | ARG | 61 | Side | ASP | 91 | Main | 50,55% |
| ARG | 61 | Main | GLU | 57 | Main | 49,55% | ARG | 61 | Main | GLU | 57 | Main | 48,05% |
| ARG | 61 | Side | ASP | 91 | Main | 45,45% | LYS | 45 | Main | PHE | 41 | Main | 47,30% |
| LYS | 45 | Main | PHE | 41 | Main | 45,35% | ARG | 102 | Main | ARG | 98 | Main | 47,00% |
| ASN | 79 | Side | GLN | 75 | Main | 45,05% | GLN | 75 | Main | GLN | 71 | Main | 46,90% |
| GLH | 66 | Main | PHE | 62 | Main | 38,75% | ARG | 61 | Side | GLU | 57 | Side | 44,50% |
| GLH | 58 | Main | ASP | 54 | Main | 36,25% | ASN | 79 | Side | GLN | 75 | Main | 41,10% |
| PHE | 108 | Main | GLY | 104 | Main | 36,15% | GLH | 66 | Main | PHE | 62 | Main | 38,10% |
| ARG | 61 | Side | GLU | 57 | Side | 33,35% | LYS | 50 | Side | ASP | 91 | Side | 35,00% |
| LYS | 47 | Main | ALA | 43 | Main | 33,20% | THR | 93 | Side | GLU | 96 | Side | 34,65% |
| ALA | 105 | Main | TYR | 101 | Main | 32,75% | THR | 93 | Main | GLU | 96 | Side | 34,50% |
| ALA | 64 | Main | PHE | 60 | Main | 31,20% | ALA | 99 | Main | GLH | 95 | Main | 33,80% |
| THR | 87 | Side | SER | 90 | Side | 31,05% | GLH | 58 | Main | ASP | 54 | Main | 32,70% |
| ARG | 100 | Side | GLU | 96 | Side | 29,90% | ALA | 64 | Main | PHE | 60 | Main | 32,60% |
| ARG | 98 | Main | ARG | 94 | Main | 28,95% | ALA | 42 | Main | GLH | 38 | Main | 32,10% |
| ARG | 100 | Main | GLU | 96 | Main | 28,80% | LYS | 47 | Main | ALA | 43 | Main | 31,65% |
| ALA | 42 | Main | GLH | 38 | Main | 28,55% | MET | 92 | Main | PHE | 89 | Main | 31,20% |
| ASN | 68 | Side | SER | 90 | Main | 28,30% | ARG | 94 | Side | GLU | 67 | Side | 30,10% |
| PHE | 97 | Main | THR | 93 | Main | 26,70% | PHE | 97 | Main | THR | 93 | Main | 28,35% |
| MET | 69 | Main | PHE | 65 | Main | 25,90% | GLN | 71 | Main | GLU | 67 | Main | 27,70% |

| | | | | | | | | | | | | | |
|---|---|---|---|---|---|---|---|---|---|---|---|---|---|
| ARG | 114 | Main | ALA | 110 | Main | 25,40% | ALA | 72 | Main | ASN | 68 | Main | 26,95% |
| PHE | 65 | Main | ARG | 61 | Main | 25,30% | LYS | 56 | Side | ASP | 54 | Side | 26,40% |
| ASN | 68 | Side | MET | 92 | Main | 24,75% | MET | 69 | Main | PHE | 65 | Main | 24,60% |
| ARG | 63 | Main | ALA | 59 | Main | 23,95% | LYS | 56 | Main | ASP | 54 | Side | 23,60% |
| TYR | 101 | Main | PHE | 97 | Main | 23,85% | THR | 87 | Main | SER | 90 | Side | 23,25% |
| PHE | 62 | Main | GLH | 58 | Main | 23,35% | PHE | 65 | Main | ARG | 61 | Main | 22,30% |
| PHE | 44 | Main | ARG | 40 | Main | 21,70% | ASN | 68 | Side | MET | 92 | Main | 21,45% |
| LYS | 46 | Main | ALA | 42 | Main | 21,45% | LYS | 46 | Main | ALA | 42 | Main | 21,15% |
| THR | 93 | Side | GLU | 67 | Side | 20,70% | PHE | 44 | Main | ARG | 40 | Main | 20,20% |
| ALA | 76 | Main | ALA | 72 | Main | 19,45% | ARG | 63 | Main | ALA | 59 | Main | 19,95% |
| ALA | 43 | Main | MET | 39 | Main | 16,75% | PHE | 62 | Main | GLH | 58 | Main | 18,90% |
| LYS | 50 | Side | ASP | 91 | Side | 16,55% | ILE | 74 | Main | GLU | 70 | Main | 18,40% |
| LYS | 73 | Side | GLU | 70 | Side | 16,55% | ALA | 109 | Main | ALA | 105 | Main | 17,50% |
| GLN | 71 | Main | GLU | 67 | Main | 16,25% | ASN | 68 | Side | SER | 90 | Main | 17,30% |
| ALA | 72 | Main | ASN | 68 | Main | 15,95% | ARG | 114 | Main | ALA | 110 | Main | 16,90% |
| LYS | 56 | Main | ASP | 54 | Side | 15,80% | ALA | 76 | Main | ALA | 72 | Main | 16,55% |
| ALA | 59 | Main | ALA | 55 | Main | 14,75% | ALA | 43 | Main | MET | 39 | Main | 16,50% |
| LYS | 50 | Side | PRO | 88 | Main | 14,45% | GLU | 67 | Main | ARG | 63 | Main | 16,25% |
| ALA | 109 | Main | ALA | 105 | Main | 13,45% | ALA | 59 | Main | ALA | 55 | Main | 15,60% |
| GLH | 58 | Side | GLH | 38 | Side | 12,85% | ALA | 110 | Main | SER | 106 | Main | 15,25% |
| ARG | 94 | Side | GLU | 67 | Side | 12,45% | ARG | 98 | Main | ARG | 94 | Main | 14,40% |
| PHE | 60 | Main | LYS | 56 | Main | 12,35% | LYS | 73 | Side | GLU | 70 | Side | 13,55% |
| GLU | 70 | Main | GLH | 66 | Main | 12,30% | PHE | 60 | Main | LYS | 56 | Main | 13,30% |
| LYS | 56 | Side | ASP | 54 | Side | 12,25% | ASN | 68 | Main | ALA | 64 | Main | 12,20% |
| TYR | 48 | Main | PHE | 44 | Main | 11,65% | TYR | 48 | Main | PHE | 44 | Main | 11,95% |
| GLU | 57 | Main | ASP | 54 | Side | 11,30% | ALA | 77 | Main | LYS | 73 | Main | 11,20% |
| ASN | 68 | Main | ALA | 64 | Main | 10,50% | ALA | 111 | Main | TYR | 107 | Main | 11,20% |
| GLU | 67 | Main | ARG | 63 | Main | 10,40% | ALA | 78 | Main | ILE | 74 | Main | 11,10% |
| LYS | 45 | Side | GLH | 58 | Side | 9,50% | ASN | 68 | Side | ALA | 64 | Main | 10,70% |
| ILE | 74 | Main | GLU | 70 | Main | 9,25% | ALA | 82 | Main | ASN | 79 | Main | 10,15% |
| TYR | 107 | Main | ASN | 103 | Main | 7,85% | LYS | 50 | Main | LYS | 45 | Main | 9,95% |
| LYS | 53 | Side | ASP | 54 | Side | 7,55% | GLN | 112 | Main | PHE | 108 | Main | 9,70% |
| THR | 93 | Side | ASN | 68 | Side | 7,50% | TYR | 101 | Main | ARG | 98 | Main | 9,55% |
| LYS | 50 | Main | LYS | 45 | Main | 6,45% | LYS | 45 | Side | GLH | 58 | Side | 8,30% |
| SER | 90 | Main | THR | 87 | Main | 6,45% | ALA | 105 | Main | ARG | 102 | Main | 7,90% |
| LYS | 113 | Main | ALA | 109 | Main | 5,80% | GLU | 57 | Main | ASP | 54 | Side | 7,30% |
| ALA | 77 | Main | LYS | 73 | Main | 5,40% | GLY | 104 | Main | TYR | 101 | Main | 7,20% |
| | | | | | | | SER | 106 | Side | ASN | 103 | Main | 7,00% |
| | | | | | | | ARG | 100 | Main | GLU | 96 | Main | 6,55% |
| | | | | | | | GLU | 96 | Main | THR | 93 | Side | 6,25% |
| | | | | | | | THR | 93 | Side | GLU | 67 | Side | 6,10% |

**Table S2b: Relative occurrence of hydrogen bonds within the pro-domain.** Four respective MD simulations at pH 4 (pH4_3 and pH4_4, see Table S2a for pH4_1 and pH4_2) were carried out. `Main' refers to interactions of the backbone. 'Side' indicates interactions of side chains. Occurrences below 5% were left out for clarity.

| pH 4_3 | | | | | | | pH 4_4 | | | | | | |
|---|---|---|---|---|---|---|---|---|---|---|---|---|---|
| donor | | | acceptor | | | occupancy | donor | | | acceptor | | | occupancy |
| ARG | 61 | Side | GLU | 96 | Side | 88,31% | ARG | 61 | Side | GLU | 96 | Side | 87,01% |
| TYR | 52 | Side | ASP | 91 | Side | 76,31% | TYR | 52 | Side | ASP | 91 | Side | 77,06% |
| ARG | 61 | Side | ASP | 91 | Main | 52,82% | ARG | 61 | Side | GLU | 57 | Side | 52,92% |
| LYS | 45 | Main | PHE | 41 | Main | 50,52% | THR | 93 | Side | GLU | 96 | Side | 49,78% |
| ARG | 61 | Side | GLU | 57 | Side | 49,08% | LYS | 45 | Main | PHE | 41 | Main | 49,53% |
| ARG | 61 | Main | GLU | 57 | Main | 44,18% | ARG | 61 | Side | ASP | 91 | Main | 48,38% |
| THR | 93 | Side | GLU | 96 | Side | 42,18% | THR | 93 | Main | GLU | 96 | Side | 45,63% |
| GLH | 66 | Main | PHE | 62 | Main | 41,43% | ARG | 61 | Main | GLU | 57 | Main | 43,48% |
| GLN | 75 | Main | GLN | 71 | Main | 40,23% | GLH | 66 | Main | PHE | 62 | Main | 41,23% |
| ASN | 79 | Side | GLN | 75 | Main | 39,48% | ARG | 102 | Main | ARG | 98 | Main | 40,73% |
| GLN | 71 | Main | GLU | 67 | Main | 38,18% | ASN | 68 | Side | SER | 90 | Main | 38,58% |
| MET | 69 | Main | PHE | 65 | Main | 36,53% | ASN | 68 | Side | MET | 92 | Main | 38,08% |
| THR | 93 | Main | GLU | 96 | Side | 35,78% | GLN | 75 | Main | GLN | 71 | Main | 36,18% |
| ASN | 68 | Side | SER | 90 | Main | 35,13% | TYR | 101 | Main | PHE | 97 | Main | 34,63% |
| ASN | 68 | Side | MET | 92 | Main | 33,58% | ALA | 42 | Main | GLH | 38 | Main | 33,68% |
| ALA | 42 | Main | GLH | 38 | Main | 33,53% | LYS | 47 | Main | ALA | 43 | Main | 33,13% |
| PHE | 97 | Main | THR | 93 | Main | 32,68% | ASN | 79 | Side | GLN | 75 | Main | 31,48% |
| LYS | 50 | Side | ASP | 91 | Side | 30,68% | ALA | 99 | Main | GLH | 95 | Main | 30,28% |
| LYS | 47 | Main | ALA | 43 | Main | 30,03% | MET | 69 | Main | PHE | 65 | Main | 29,29% |
| ALA | 99 | Main | GLH | 95 | Main | 29,24% | ARG | 98 | Main | ARG | 94 | Main | 27,44% |
| GLH | 58 | Main | ASP | 54 | Main | 28,84% | GLH | 58 | Main | ASP | 54 | Main | 26,34% |
| SER | 106 | Side | ASN | 103 | Main | 27,64% | PHE | 65 | Main | ARG | 61 | Main | 26,29% |
| ALA | 64 | Main | PHE | 60 | Main | 25,94% | GLN | 71 | Main | GLU | 67 | Main | 26,29% |
| THR | 87 | Main | SER | 90 | Side | 23,89% | GLU | 57 | Main | ASP | 54 | Side | 25,64% |
| PHE | 65 | Main | ARG | 61 | Main | 23,84% | ARG | 94 | Side | GLU | 67 | Side | 25,24% |
| ALA | 72 | Main | ASN | 68 | Main | 23,49% | THR | 87 | Main | SER | 90 | Side | 25,04% |
| LYS | 46 | Main | ALA | 42 | Main | 23,24% | ALA | 105 | Main | TYR | 101 | Main | 24,94% |
| MET | 92 | Main | PHE | 89 | Main | 22,54% | ARG | 63 | Main | ALA | 59 | Main | 24,74% |
| PHE | 62 | Main | GLH | 58 | Main | 22,44% | LYS | 50 | Side | ASP | 91 | Side | 24,39% |
| ARG | 63 | Main | ALA | 59 | Main | 22,19% | ALA | 64 | Main | PHE | 60 | Main | 23,89% |
| ASN | 68 | Main | ALA | 64 | Main | 21,99% | MET | 92 | Main | PHE | 89 | Main | 22,99% |
| ALA | 105 | Main | TYR | 101 | Main | 21,84% | ALA | 72 | Main | ASN | 68 | Main | 21,34% |
| GLN | 71 | Side | GLU | 67 | Side | 21,24% | LYS | 46 | Main | ALA | 42 | Main | 21,09% |
| ALA | 76 | Main | ALA | 72 | Main | 20,84% | ALA | 76 | Main | ALA | 72 | Main | 20,59% |
| ILE | 74 | Main | GLU | 70 | Main | 20,14% | PHE | 44 | Main | ARG | 40 | Main | 18,69% |
| PHE | 44 | Main | ARG | 40 | Main | 20,09% | PHE | 62 | Main | GLH | 58 | Main | 18,29% |
| SER | 90 | Side | THR | 87 | Main | 19,24% | LYS | 73 | Side | GLU | 70 | Side | 17,79% |
| ALA | 43 | Main | MET | 39 | Main | 18,54% | PHE | 60 | Main | LYS | 56 | Main | 17,39% |
| ARG | 98 | Main | ARG | 94 | Main | 18,19% | ASN | 68 | Main | ALA | 64 | Main | 17,14% |
| GLU | 57 | Main | ASP | 54 | Side | 15,79% | ALA | 43 | Main | MET | 39 | Main | 16,99% |
| ALA | 59 | Main | ALA | 55 | Main | 15,54% | LYS | 113 | Main | ALA | 109 | Main | 16,69% |

| | | | | | | | | | | | | | |
|---|---|---|---|---|---|---|---|---|---|---|---|---|---|
| ALA | 109 | Main | ALA | 105 | Main | 15,19% | ALA | 59 | Main | ALA | 55 | Main | 15,74% |
| ARG | 94 | Side | GLU | 67 | Side | 14,74% | GLN | 71 | Side | GLU | 67 | Side | 14,99% |
| PHE | 60 | Main | LYS | 56 | Main | 13,44% | ILE | 74 | Main | GLU | 70 | Main | 14,29% |
| ALA | 111 | Main | TYR | 107 | Main | 13,44% | GLU | 67 | Main | ARG | 63 | Main | 12,64% |
| LYS | 73 | Side | GLU | 70 | Side | 13,34% | PHE | 97 | Main | THR | 93 | Main | 12,09% |
| GLN | 112 | Main | PHE | 108 | Main | 13,14% | ALA | 77 | Main | LYS | 73 | Main | 11,49% |
| ALA | 110 | Main | SER | 106 | Main | 13,14% | LYS | 50 | Main | LYS | 45 | Main | 11,24% |
| LYS | 56 | Main | ASP | 54 | Side | 12,74% | LYS | 53 | Side | ASP | 54 | Side | 10,69% |
| ALA | 77 | Main | LYS | 73 | Main | 12,44% | ALA | 109 | Main | ALA | 105 | Main | 10,59% |
| LYS | 50 | Main | LYS | 45 | Main | 11,74% | GLY | 104 | Main | ARG | 100 | Main | 9,90% |
| GLU | 67 | Main | ARG | 63 | Main | 11,59% | TYR | 48 | Main | PHE | 44 | Main | 9,45% |
| PHE | 108 | Main | GLY | 104 | Main | 10,34% | LYS | 45 | Side | GLH | 58 | Side | 9,30% |
| ARG | 114 | Main | ALA | 110 | Main | 9,95% | SER | 106 | Side | ASN | 103 | Main | 9,25% |
| LYS | 45 | Side | GLH | 58 | Side | 9,40% | ARG | 114 | Main | ALA | 110 | Main | 8,95% |
| GLU | 96 | Main | THR | 93 | Side | 9,00% | THR | 123 | Side | VAL | 121 | Main | 7,65% |
| GLN | 75 | Side | GLN | 71 | Main | 8,90% | LYS | 56 | Main | ASP | 54 | Side | 7,50% |
| TYR | 48 | Main | PHE | 44 | Main | 8,45% | GLN | 112 | Main | PHE | 108 | Main | 6,80% |
| ARG | 100 | Main | GLU | 96 | Main | 8,20% | PHE | 108 | Main | GLY | 104 | Main | 6,45% |
| THR | 123 | Side | VAL | 121 | Main | 5,45% | ALA | 82 | Main | ASN | 79 | Main | 6,30% |
| LYS | 50 | Side | PRO | 88 | Main | 5,00% | ALA | 78 | Main | ILE | 74 | Main | 6,20% |
| | | | | | | | ALA | 111 | Main | PHE | 108 | Main | 5,85% |
| | | | | | | | SER | 106 | Side | ARG | 102 | Main | 5,05% |

**Table S3a: Relative occurrence of hydrogen bonds within the pro-domain.** Four respective MD simulations at pH 8 (pH8_1 and pH8_2, see Table S3b for pH8_3 and pH8_4) were carried out. `Main' refers to interactions of the backbone. 'Side' indicates interactions of side chains. Occurrences below 5% were left out for clarity.

| pH8_1 | | | | | | | pH8_2 | | | | | | |
|---|---|---|---|---|---|---|---|---|---|---|---|---|---|
| Found | | | 153 | | | hbonds, | Found | | | 165 | | | hbonds, |
| donor | | | acceptor | | | occupancy | donor | | | acceptor | | | occupancy |
| ARG | 102 | Side | GLU | 95 | Side | 71,90% | ARG | 61 | Side | GLU | 96 | Side | 81,65% |
| TYR | 52 | Side | ASP | 91 | Side | 67,00% | TYR | 52 | Side | ASP | 91 | Side | 76,05% |
| ASN | 68 | Side | GLY | 85 | Main | 50,45% | THR | 93 | Side | GLU | 96 | Side | 52,45% |
| GLN | 71 | Main | GLU | 67 | Main | 49,05% | ARG | 61 | Side | ASP | 91 | Main | 49,90% |
| LYS | 45 | Side | GLU | 58 | Side | 44,20% | ASN | 68 | Side | SER | 90 | Main | 49,65% |
| LYS | 45 | Main | PHE | 41 | Main | 41,55% | LYS | 45 | Main | PHE | 41 | Main | 47,80% |
| ARG | 61 | Main | GLU | 57 | Main | 38,45% | LYS | 45 | Side | GLU | 58 | Side | 45,60% |
| ARG | 40 | Side | GLU | 66 | Side | 37,80% | ARG | 61 | Main | GLU | 57 | Main | 43,30% |
| ARG | 61 | Side | ASP | 91 | Main | 36,50% | ARG | 61 | Side | GLU | 57 | Side | 37,65% |
| ASN | 79 | Side | GLN | 75 | Main | 34,70% | THR | 93 | Main | GLU | 96 | Side | 35,80% |
| GLU | 58 | Main | ASP | 54 | Main | 34,35% | ALA | 42 | Main | GLU | 38 | Main | 35,40% |
| ARG | 98 | Main | ARG | 94 | Main | 34,35% | MET | 69 | Main | PHE | 65 | Main | 35,15% |
| ARG | 61 | Side | GLU | 96 | Side | 33,85% | ASN | 68 | Side | MET | 92 | Main | 35,05% |
| LYS | 47 | Main | ALA | 43 | Main | 33,65% | GLN | 75 | Main | GLN | 71 | Main | 34,60% |
| ALA | 43 | Main | MET | 39 | Main | 33,10% | GLU | 66 | Main | PHE | 62 | Main | 34,25% |
| LYS | 73 | Side | GLU | 70 | Side | 31,40% | ARG | 102 | Side | GLU | 95 | Side | 34,05% |

| | | | | | | | | | | | | | |
|---|---|---|---|---|---|---|---|---|---|---|---|---|---|
| SER | 106 | Side | ASN | 103 | Main | 30,75% | ASN | 79 | Side | GLN | 75 | Main | 33,45% |
| GLU | 66 | Main | PHE | 62 | Main | 28,85% | THR | 87 | Main | SER | 90 | Side | 32,60% |
| ARG | 61 | Side | GLU | 57 | Side | 28,75% | LYS | 47 | Main | ALA | 43 | Main | 29,20% |
| PHE | 62 | Main | GLU | 58 | Main | 27,80% | PHE | 62 | Main | GLU | 58 | Main | 29,10% |
| LYS | 46 | Main | ALA | 42 | Main | 26,45% | PHE | 65 | Main | ARG | 61 | Main | 28,40% |
| ALA | 76 | Main | ALA | 72 | Main | 26,20% | LYS | 50 | Side | ASP | 91 | Side | 28,25% |
| ARG | 94 | Side | GLU | 67 | Side | 26,05% | ALA | 105 | Main | TYR | 101 | Main | 28,05% |
| LYS | 50 | Side | ASP | 91 | Side | 25,45% | ALA | 43 | Main | MET | 39 | Main | 27,95% |
| ALA | 42 | Main | GLU | 38 | Main | 25,40% | ARG | 98 | Main | ARG | 94 | Main | 27,35% |
| THR | 87 | Main | SER | 90 | Side | 23,85% | ALA | 64 | Main | PHE | 60 | Main | 25,10% |
| ARG | 94 | Side | GLN | 71 | Side | 23,55% | MET | 92 | Main | PHE | 89 | Main | 23,40% |
| THR | 93 | Side | GLU | 96 | Side | 23,30% | LYS | 46 | Main | ALA | 42 | Main | 22,75% |
| PHE | 65 | Main | ARG | 61 | Main | 22,90% | GLU | 58 | Main | ASP | 54 | Main | 22,50% |
| ALA | 99 | Main | GLU | 95 | Main | 22,40% | SER | 106 | Side | ASN | 103 | Main | 22,25% |
| PHE | 97 | Main | THR | 93 | Main | 18,35% | ALA | 99 | Main | GLU | 95 | Main | 21,65% |
| ARG | 63 | Main | ALA | 59 | Main | 17,60% | LYS | 73 | Side | GLU | 70 | Side | 20,90% |
| ALA | 72 | Main | ASN | 68 | Main | 16,90% | ALA | 111 | Main | TYR | 107 | Main | 19,65% |
| MET | 92 | Main | PHE | 89 | Main | 15,40% | PHE | 108 | Main | GLY | 104 | Main | 19,00% |
| ARG | 114 | Main | ALA | 110 | Main | 15,15% | PHE | 97 | Main | THR | 93 | Main | 17,80% |
| ALA | 77 | Main | LYS | 73 | Main | 15,10% | PHE | 44 | Main | ARG | 40 | Main | 17,05% |
| THR | 93 | Main | GLU | 96 | Side | 14,70% | ALA | 72 | Main | ASN | 68 | Main | 16,85% |
| SER | 90 | Side | ASN | 68 | Side | 14,30% | ARG | 63 | Main | ALA | 59 | Main | 16,05% |
| PHE | 108 | Main | GLY | 104 | Main | 14,25% | ALA | 76 | Main | ALA | 72 | Main | 15,80% |
| LYS | 50 | Side | PRO | 88 | Main | 13,25% | ALA | 59 | Main | ALA | 55 | Main | 14,05% |
| GLN | 75 | Side | GLN | 71 | Main | 12,60% | GLN | 71 | Main | GLU | 67 | Main | 12,85% |
| ALA | 111 | Main | TYR | 107 | Main | 12,30% | ARG | 116 | Side | ALA | 111 | Main | 12,60% |
| ASN | 68 | Main | ALA | 64 | Main | 11,35% | ALA | 77 | Main | LYS | 73 | Main | 12,25% |
| LYS | 56 | Main | ASP | 54 | Side | 11,25% | ARG | 102 | Main | ARG | 98 | Main | 12,20% |
| LYS | 50 | Main | LYS | 45 | Main | 11,15% | ARG | 94 | Side | GLU | 67 | Side | 11,90% |
| ALA | 64 | Main | PHE | 60 | Main | 10,50% | GLU | 57 | Main | ASP | 54 | Side | 11,70% |
| THR | 123 | Side | VAL | 121 | Main | 10,40% | LYS | 50 | Main | LYS | 45 | Main | 11,60% |
| MET | 69 | Main | PHE | 65 | Main | 10,35% | ASN | 68 | Main | ALA | 64 | Main | 10,45% |
| ALA | 59 | Main | ALA | 55 | Main | 9,95% | TYR | 48 | Main | PHE | 44 | Main | 9,50% |
| TYR | 48 | Main | PHE | 44 | Main | 9,30% | ARG | 100 | Side | ARG | 100 | Main | 9,50% |
| GLU | 57 | Main | ASP | 54 | Side | 9,20% | SER | 106 | Side | ARG | 102 | Main | 9,20% |
| ARG | 63 | Side | GLU | 67 | Side | 8,50% | ALA | 78 | Main | ILE | 74 | Main | 8,90% |
| ARG | 40 | Main | GLU | 38 | Side | 8,40% | ALA | 109 | Main | ALA | 105 | Main | 8,35% |
| PHE | 44 | Main | ARG | 40 | Main | 8,30% | LYS | 53 | Main | GLU | 57 | Side | 8,30% |
| ARG | 61 | Side | ASP | 91 | Side | 8,20% | PHE | 60 | Main | LYS | 56 | Main | 7,60% |
| ALA | 82 | Main | ASN | 79 | Main | 7,75% | ALA | 82 | Main | ASN | 79 | Main | 6,75% |
| LYS | 113 | Main | ALA | 109 | Main | 7,65% | ILE | 74 | Main | GLU | 70 | Main | 6,45% |
| LYS | 56 | Side | ASP | 54 | Side | 7,00% | ARG | 63 | Side | GLU | 67 | Side | 6,40% |
| ALA | 105 | Main | TYR | 101 | Main | 6,75% | ARG | 40 | Main | GLU | 38 | Side | 6,35% |
| ALA | 109 | Main | ALA | 105 | Main | 6,70% | TYR | 107 | Main | ASN | 103 | Main | 5,30% |
| GLN | 112 | Side | PHE | 108 | Main | 6,00% | GLN | 112 | Side | ALA | 109 | Main | 5,10% |
| GLN | 75 | Main | GLN | 71 | Main | 5,60% | GLU | 67 | Main | ARG | 63 | Main | 5,05% |
| ALA | 78 | Main | ILE | 74 | Main | 5,55% | | | | | | | |

| | | | | | | |
|---|---|---|---|---|---|---|
| GLU | 67 | Main | ARG | 63 | Main | 5,00% |

**Table S3b: Relative occurrence of hydrogen bonds within the pro-domain.** Four respective MD simulations at pH 8 (pH8_3 and pH8_4, see Table S3a for pH8_1 and pH8_2) were carried out. \`Main' refers to interactions of the backbone. 'Side' indicates interactions of side chains. Occurrences below 5% were left out for clarity.

| pH_8_3 | | | | | | | pH8_4 | | | | | | |
|---|---|---|---|---|---|---|---|---|---|---|---|---|---|
| Found | | | 166 | | | hbonds, | Found | | | 162 | | | hbonds, |
| donor | | | acceptor | | | occupancy | donor | | | acceptor | | | occupancy |
| TYR | 52 | Side | ASP | 91 | Side | 74,56% | ARG | 61 | Side | GLU | 96 | Side | 85,41% |
| ARG | 61 | Side | GLU | 96 | Side | 72,86% | TYR | 52 | Side | ASP | 91 | Side | 78,61% |
| SER | 90 | Side | ASN | 68 | Side | 66,12% | SER | 90 | Side | ASN | 68 | Side | 58,27% |
| THR | 93 | Side | GLU | 96 | Side | 62,82% | LYS | 45 | Main | PHE | 41 | Main | 46,18% |
| THR | 93 | Main | GLU | 96 | Side | 53,17% | ARG | 61 | Side | ASP | 91 | Main | 45,53% |
| ARG | 94 | Side | GLU | 67 | Side | 44,48% | GLN | 75 | Main | GLN | 71 | Main | 45,13% |
| ARG | 61 | Main | GLU | 57 | Main | 43,53% | LYS | 45 | Side | GLU | 58 | Side | 43,53% |
| LYS | 45 | Side | GLU | 58 | Side | 42,33% | ARG | 40 | Side | GLU | 66 | Side | 41,63% |
| ALA | 76 | Main | ALA | 72 | Main | 41,18% | ARG | 61 | Main | GLU | 57 | Main | 40,43% |
| ARG | 98 | Side | GLU | 95 | Side | 38,08% | ARG | 61 | Side | GLU | 57 | Side | 39,63% |
| LYS | 45 | Main | PHE | 41 | Main | 35,28% | ARG | 98 | Main | ARG | 94 | Main | 36,33% |
| LYS | 47 | Main | ALA | 43 | Main | 34,58% | ASN | 79 | Side | GLN | 75 | Main | 34,23% |
| ALA | 42 | Main | GLU | 38 | Main | 32,03% | GLU | 66 | Main | PHE | 62 | Main | 32,98% |
| ARG | 63 | Side | GLU | 67 | Side | 31,73% | GLN | 71 | Main | GLU | 67 | Main | 32,83% |
| LYS | 46 | Main | ALA | 42 | Main | 31,08% | ALA | 72 | Main | ASN | 68 | Main | 32,03% |
| ARG | 40 | Side | GLU | 66 | Side | 30,73% | GLU | 58 | Main | ASP | 54 | Main | 31,93% |
| GLU | 58 | Main | ASP | 54 | Main | 30,58% | THR | 93 | Side | GLU | 96 | Side | 31,78% |
| ARG | 61 | Side | ASP | 91 | Main | 29,49% | LYS | 47 | Main | ALA | 43 | Main | 30,58% |
| ARG | 63 | Side | ARG | 63 | Main | 29,14% | THR | 93 | Main | GLU | 96 | Side | 29,69% |
| ARG | 94 | Main | GLU | 67 | Side | 28,69% | ASN | 68 | Side | GLY | 85 | Main | 28,19% |
| PHE | 65 | Main | ARG | 61 | Main | 28,54% | PHE | 62 | Main | GLU | 58 | Main | 28,04% |
| LYS | 50 | Side | ASP | 91 | Side | 27,24% | PHE | 65 | Main | ARG | 61 | Main | 27,59% |
| ARG | 61 | Side | GLU | 57 | Side | 26,94% | ARG | 98 | Side | GLU | 95 | Side | 26,79% |
| ALA | 43 | Main | MET | 39 | Main | 25,74% | LYS | 46 | Main | ALA | 42 | Main | 25,84% |
| ASN | 79 | Side | GLN | 75 | Main | 25,04% | LYS | 50 | Side | ASP | 91 | Side | 24,94% |
| PHE | 97 | Main | THR | 93 | Main | 23,39% | SER | 90 | Main | THR | 87 | Main | 23,84% |
| ARG | 98 | Main | ARG | 94 | Main | 21,44% | MET | 69 | Main | PHE | 65 | Main | 22,54% |
| ALA | 99 | Main | GLU | 95 | Main | 20,99% | LYS | 73 | Side | GLU | 70 | Side | 20,49% |
| ALA | 105 | Main | TYR | 101 | Main | 19,94% | TYR | 48 | Main | PHE | 44 | Main | 19,59% |
| LYS | 56 | Main | ASP | 54 | Side | 19,44% | PHE | 97 | Main | THR | 93 | Main | 19,14% |
| THR | 87 | Main | SER | 90 | Side | 17,54% | TYR | 101 | Main | PHE | 97 | Main | 18,54% |
| MET | 92 | Main | PHE | 89 | Main | 17,34% | ALA | 43 | Main | MET | 39 | Main | 17,19% |
| ALA | 111 | Main | TYR | 107 | Main | 17,24% | ALA | 76 | Main | ALA | 72 | Main | 16,89% |
| LYS | 56 | Side | ASP | 54 | Side | 16,99% | ARG | 63 | Main | ALA | 59 | Main | 14,84% |
| LYS | 73 | Side | GLU | 70 | Side | 16,99% | GLY | 104 | Main | TYR | 101 | Main | 14,84% |
| ARG | 102 | Side | GLU | 95 | Side | 16,29% | MET | 92 | Main | PHE | 89 | Main | 14,64% |
| GLU | 67 | Main | ALA | 64 | Main | 15,64% | ALA | 59 | Main | ALA | 55 | Main | 14,44% |

| | | | | | | | | | | | | | |
|---|---|---|---|---|---|---|---|---|---|---|---|---|---|
| SER | 106 | Side | ASN | 103 | Main | 14,69% | ARG | 114 | Main | ALA | 110 | Main | 13,89% |
| PHE | 62 | Main | GLU | 58 | Main | 13,84% | ALA | 64 | Main | PHE | 60 | Main | 12,94% |
| ALA | 110 | Main | SER | 106 | Main | 13,54% | LYS | 50 | Main | LYS | 45 | Main | 12,84% |
| GLN | 75 | Main | GLN | 71 | Main | 12,69% | ARG | 102 | Main | ARG | 98 | Main | 11,99% |
| LYS | 50 | Main | LYS | 45 | Main | 12,34% | THR | 87 | Main | SER | 90 | Side | 11,29% |
| SER | 90 | Main | THR | 87 | Main | 12,19% | ALA | 77 | Main | LYS | 73 | Main | 11,04% |
| PHE | 108 | Main | GLY | 104 | Main | 11,14% | LYS | 56 | Main | ASP | 54 | Side | 11,04% |
| ARG | 102 | Main | ARG | 98 | Main | 10,94% | ALA | 78 | Main | ILE | 74 | Main | 10,99% |
| ARG | 102 | Side | ARG | 98 | Main | 10,69% | LYS | 56 | Side | ASP | 54 | Side | 10,39% |
| ALA | 109 | Main | ALA | 105 | Main | 10,19% | ALA | 105 | Main | ARG | 102 | Main | 9,35% |
| ALA | 59 | Main | ALA | 55 | Main | 10,04% | PHE | 44 | Main | ARG | 40 | Main | 9,05% |
| GLN | 71 | Main | GLU | 67 | Main | 9,45% | ALA | 99 | Main | GLU | 95 | Main | 8,95% |
| GLN | 112 | Main | PHE | 108 | Main | 9,25% | ALA | 42 | Main | GLU | 38 | Main | 8,90% |
| PHE | 44 | Main | ARG | 40 | Main | 9,20% | LYS | 113 | Main | ALA | 109 | Main | 8,65% |
| ALA | 72 | Main | ASN | 68 | Main | 9,15% | ALA | 109 | Main | ALA | 105 | Main | 8,20% |
| GLU | 70 | Main | GLU | 66 | Main | 9,15% | ALA | 82 | Main | ASN | 79 | Main | 6,95% |
| GLN | 112 | Side | PHE | 108 | Main | 9,15% | ARG | 94 | Side | GLU | 67 | Side | 6,50% |
| THR | 123 | Side | THR | 122 | Main | 8,65% | PHE | 108 | Main | GLY | 104 | Main | 6,35% |
| LYS | 45 | Side | GLU | 38 | Side | 8,10% | SER | 106 | Side | ASN | 103 | Main | 6,25% |
| LYS | 73 | Side | GLU | 70 | Main | 7,40% | ARG | 116 | Side | ALA | 111 | Main | 5,75% |
| ALA | 64 | Main | PHE | 60 | Main | 6,90% | GLU | 57 | Main | ASP | 54 | Side | 5,70% |
| GLU | 66 | Main | PHE | 62 | Main | 6,65% | GLN | 112 | Main | ALA | 109 | Main | 5,65% |
| ALA | 78 | Main | ILE | 74 | Main | 6,50% | ASN | 79 | Main | GLN | 75 | Main | 5,50% |
| ARG | 94 | Side | GLU | 70 | Side | 6,40% | PHE | 60 | Main | LYS | 56 | Main | 5,40% |
| ARG | 114 | Main | ALA | 111 | Main | 6,15% | | | | | | | |
| TYR | 48 | Main | PHE | 44 | Main | 5,70% | | | | | | | |
| ALA | 77 | Main | LYS | 73 | Main | 5,55% | | | | | | | |
| GLY | 104 | Main | TYR | 101 | Main | 5,50% | | | | | | | |
| ARG | 40 | Main | GLU | 38 | Side | 5,25% | | | | | | | |
| ARG | 114 | Main | ALA | 110 | Main | 5,20% | | | | | | | |
| ASN | 79 | Main | GLN | 75 | Main | 5,20% | | | | | | | |
| TYR | 101 | Main | ARG | 98 | Main | 5,00% | | | | | | | |

**Table S4a: Relative occurrence of hydrogen bonds within the catalytic domain.** Four respective MD simulations at pH 4 (pH4_1 and pH4_2, see Table S4b for pH4_3 and pH4_4) were carried out. `Main' refers to interactions of the backbone. 'Side' indicates interactions of side chains. Occurrences below 5% were left out for clarity.

| **pH4_1** | | | | | | | **pH4_2** | | | | | | |
|---|---|---|---|---|---|---|---|---|---|---|---|---|---|
| **donor** | | | acceptor | | | occupancy | donor | | | acceptor | | | occupancy |
| **ASH** | 185 | Side | ASP | 182 | Side | 86,05% | ASH | 185 | Side | ASP | 182 | Side | 78,20% |
| **THR** | 267 | Side | ASP | 265 | Side | 77,00% | THR | 267 | Side | ASP | 265 | Side | 78,00% |
| **GLH** | 175 | Side | GLU | 211 | Side | 76,25% | ASH | 194 | Side | ASP | 242 | Side | 74,20% |
| **THR** | 155 | Side | TRP | 151 | Main | 75,65% | THR | 155 | Side | TRP | 151 | Main | 72,20% |
| **TYR** | 272 | Side | GLY | 275 | Main | 71,00% | TYR | 272 | Side | GLY | 275 | Main | 69,40% |
| **TRP** | 303 | Main | ILE | 321 | Main | 69,20% | TRP | 303 | Main | ILE | 321 | Main | 68,70% |
| **SER** | 174 | Side | TYR | 214 | Main | 67,40% | TYR | 294 | Side | PRO | 127 | Main | 66,35% |

| | | | | | | | | | | | | | |
|---|---|---|---|---|---|---|---|---|---|---|---|---|---|
| **ASH** | 194 | Side | ASP | 242 | Side | 64,60% | ARG | 320 | Side | GLU | 315 | Side | 65,25% |
| **LYS** | 306 | Main | LEU | 290 | Main | 61,10% | GLH | 175 | Side | GLU | 211 | Side | 62,45% |
| **THR** | 139 | Side | GLU | 160 | Side | 61,00% | GLH | 283 | Main | ASP | 265 | Side | 60,75% |
| **VAL** | 241 | Main | SER | 336 | Main | 60,00% | VAL | 241 | Main | SER | 336 | Main | 57,20% |
| **GLH** | 283 | Main | ASP | 265 | Side | 58,75% | PHE | 197 | Main | MET | 193 | Main | 57,05% |
| **SER** | 218 | Side | GLU | 222 | Main | 58,15% | LYS | 306 | Main | LEU | 290 | Main | 56,75% |
| **TYR** | 216 | Side | GLN | 144 | Main | 56,95% | ILE | 156 | Main | ALA | 152 | Main | 54,90% |
| **PHE** | 197 | Main | MET | 193 | Main | 54,80% | SER | 174 | Side | TYR | 214 | Main | 54,90% |
| **ARG** | 320 | Side | GLU | 315 | Side | 54,60% | SER | 174 | Main | PHE | 209 | Main | 53,95% |
| **GLN** | 162 | Main | ASN | 158 | Main | 53,70% | TRP | 132 | Main | VAL | 292 | Main | 53,90% |
| **ILE** | 156 | Main | ALA | 152 | Main | 52,45% | ASH | 296 | Main | ALA | 128 | Main | 53,60% |
| **SER** | 154 | Side | VAL | 289 | Main | 52,20% | SER | 218 | Side | GLU | 222 | Main | 53,20% |
| **ARG** | 133 | Side | ASP | 131 | Side | 52,10% | GLN | 144 | Side | CYX | 147 | Main | 51,70% |
| **SER** | 154 | Main | CYM | 150 | Main | 51,60% | VAL | 217 | Main | GLN | 176 | Side | 51,65% |
| **LEU** | 173 | Main | GLU | 160 | Side | 51,30% | CYX | 280 | Main | GLN | 327 | Side | 51,30% |
| **SER** | 174 | Main | PHE | 209 | Main | 50,40% | HIP | 287 | Side | ASN | 307 | Side | 50,40% |
| **GLU** | 160 | Main | ILE | 156 | Main | 50,15% | TYR | 216 | Side | GLN | 144 | Main | 50,15% |
| **GLN** | 144 | Side | CYX | 147 | Main | 49,90% | GLN | 162 | Main | ASN | 158 | Main | 49,90% |
| **HIP** | 287 | Side | ASN | 307 | Side | 48,95% | GLN | 284 | Main | ASP | 265 | Side | 48,60% |
| **ALA** | 266 | Main | ASH | 286 | Side | 48,60% | LYS | 323 | Main | PRO | 301 | Main | 46,85% |
| **VAL** | 217 | Main | GLN | 176 | Side | 45,35% | GLH | 322 | Main | LEU | 277 | Main | 45,30% |
| **SER** | 308 | Side | GLN | 144 | Side | 44,85% | LEU | 173 | Main | GLU | 160 | Side | 44,65% |
| **SER** | 298 | Side | PRO | 300 | Main | 44,70% | THR | 238 | Main | VAL | 338 | Main | 43,95% |
| **SER** | 180 | Main | GLN | 176 | Main | 44,25% | GLU | 211 | Main | SER | 172 | Main | 43,95% |
| **GLH** | 322 | Main | LEU | 277 | Main | 44,20% | ASN | 168 | Side | TRP | 163 | Main | 43,75% |
| **ASH** | 296 | Main | ALA | 128 | Main | 43,75% | GLN | 162 | Side | ASN | 257 | Main | 43,05% |
| **TRP** | 132 | Main | VAL | 292 | Main | 43,15% | SER | 213 | Side | THR | 210 | Side | 43,05% |
| **VAL** | 264 | Main | HIP | 287 | Main | 42,55% | SER | 298 | Side | PRO | 300 | Main | 42,80% |
| **GLN** | 176 | Side | SER | 218 | Side | 42,20% | SER | 180 | Main | GLN | 176 | Main | 41,15% |
| **LEU** | 291 | Main | LEU | 260 | Main | 41,85% | GLU | 160 | Main | ILE | 156 | Main | 40,60% |
| **VAL** | 289 | Main | ILE | 262 | Main | 41,40% | SER | 154 | Main | CYM | 150 | Main | 40,10% |
| **LYS** | 323 | Main | PRO | 301 | Main | 41,35% | GLY | 234 | Main | VAL | 208 | Main | 39,95% |
| **SER** | 213 | Side | THR | 210 | Side | 41,20% | ASN | 202 | Main | ASN | 198 | Main | 39,90% |
| **SER** | 282 | Side | CYX | 280 | Main | 40,35% | GLY | 275 | Main | GLU | 315 | Side | 39,75% |
| **ALA** | 337 | Main | ASN | 158 | Side | 40,15% | ASN | 158 | Side | SER | 336 | Side | 39,60% |
| **ILE** | 237 | Main | GLY | 206 | Main | 39,80% | LEU | 291 | Main | LEU | 260 | Main | 39,45% |
| **CYX** | 280 | Main | GLN | 327 | Side | 39,75% | GLN | 176 | Side | SER | 218 | Side | 39,35% |
| **ASN** | 158 | Side | SER | 336 | Side | 39,45% | ILE | 237 | Main | GLY | 206 | Main | 39,05% |
| **GLY** | 275 | Main | GLU | 315 | Side | 37,95% | SER | 282 | Side | CYX | 280 | Main | 38,30% |
| **ASN** | 168 | Side | TRP | 163 | Main | 37,90% | VAL | 165 | Main | GLY | 161 | Main | 38,00% |
| **GLY** | 324 | Main | GLU | 247 | Side | 37,90% | LEU | 254 | Main | ILE | 250 | Main | 37,85% |
| **LEU** | 254 | Main | ILE | 250 | Main | 37,60% | ILE | 250 | Main | ASP | 246 | Main | 37,20% |
| **GLY** | 234 | Main | VAL | 208 | Main | 37,40% | ALA | 337 | Main | ASN | 158 | Side | 36,75% |
| **TRP** | 303 | Side | GLU | 247 | Side | 36,60% | THR | 139 | Side | GLU | 160 | Side | 36,45% |
| **VAL** | 165 | Main | GLY | 161 | Main | 36,45% | VAL | 264 | Main | HIP | 287 | Main | 35,20% |
| **ILE** | 304 | Main | GLY | 293 | Main | 36,25% | VAL | 289 | Main | ILE | 262 | Main | 35,05% |
| **GLU** | 211 | Main | SER | 172 | Main | 36,20% | TRP | 303 | Side | GLU | 247 | Side | 34,55% |

| | | | | | | | | | | | | | |
|---|---|---|---|---|---|---|---|---|---|---|---|---|---|
| **ASN** | 202 | Main | ASN | 198 | Main | 36,15% | VAL | 338 | Main | ASP | 239 | Main | 34,00% |
| **VAL** | 338 | Main | ASP | 239 | Main | 36,15% | ASN | 331 | Main | ASN | 326 | Side | 33,45% |
| **GLN** | 162 | Side | ASN | 257 | Main | 36,05% | HIP | 287 | Main | VAL | 264 | Main | 33,15% |
| **ILE** | 250 | Main | ASP | 246 | Main | 34,50% | GLN | 176 | Side | SER | 180 | Side | 32,20% |
| **GLN** | 284 | Main | ASP | 265 | Side | 33,55% | THR | 267 | Main | ASP | 265 | Side | 31,90% |
| **THR** | 267 | Main | ASP | 265 | Side | 33,00% | GLY | 324 | Main | GLU | 247 | Side | 31,70% |
| **ASN** | 331 | Main | ASN | 326 | Side | 32,45% | ASH | 286 | Main | VAL | 264 | Main | 31,65% |
| **THR** | 238 | Main | VAL | 338 | Main | 32,30% | HIP | 287 | Side | CYM | 150 | Side | 30,65% |
| **ILE** | 305 | Main | ILE | 319 | Main | 32,20% | ASN | 204 | Side | ASN | 207 | Main | 30,60% |
| **GLN** | 176 | Side | SER | 180 | Side | 31,85% | GLN | 144 | Main | SER | 308 | Main | 30,15% |
| **ASN** | 273 | Side | TYR | 272 | Main | 31,80% | ILE | 305 | Main | ILE | 319 | Main | 29,90% |
| **ASH** | 286 | Main | VAL | 264 | Main | 31,25% | ILE | 304 | Main | GLY | 293 | Main | 29,60% |
| **THR** | 238 | Side | GLY | 341 | Main | 30,40% | TRP | 163 | Main | ILE | 159 | Main | 27,65% |
| **ASN** | 257 | Main | TYR | 253 | Main | 30,00% | SER | 335 | Main | ALA | 261 | Main | 27,25% |
| **LYS** | 142 | Side | GLU | 160 | Side | 29,85% | ILE | 184 | Main | ASP | 182 | Side | 26,55% |
| **VAL** | 292 | Main | ILE | 304 | Main | 29,25% | VAL | 208 | Main | ALA | 235 | Main | 26,35% |
| **SER** | 335 | Main | ALA | 261 | Main | 28,90% | ASN | 326 | Main | GLU | 247 | Side | 26,25% |
| **LEU** | 277 | Main | ARG | 320 | Main | 28,85% | LEU | 260 | Main | LEU | 291 | Main | 26,20% |
| **SER** | 336 | Main | VAL | 241 | Main | 28,70% | VAL | 292 | Main | ILE | 304 | Main | 25,30% |
| **ASN** | 204 | Side | ASN | 207 | Main | 27,95% | ASN | 273 | Side | TYR | 272 | Main | 24,55% |
| **TRP** | 313 | Side | PHE | 269 | Main | 26,90% | GLN | 327 | Side | THR | 325 | Side | 24,10% |
| **SER** | 180 | Side | PRO | 224 | Main | 26,75% | SER | 336 | Main | VAL | 241 | Main | 23,55% |
| **GLN** | 327 | Side | THR | 325 | Side | 26,50% | ASN | 295 | Main | TYR | 302 | Main | 23,55% |
| **TRP** | 163 | Main | ILE | 159 | Main | 26,30% | ASN | 257 | Main | TYR | 253 | Main | 23,50% |
| **THR** | 210 | Main | GLH | 232 | Main | 25,70% | CYX | 188 | Main | GLN | 223 | Side | 22,05% |
| **LEU** | 243 | Main | VAL | 334 | Main | 24,75% | GLY | 340 | Main | ALA | 236 | Main | 21,80% |
| **VAL** | 208 | Main | ALA | 235 | Main | 24,70% | ALA | 137 | Main | TRP | 132 | Main | 20,95% |
| **ASN** | 326 | Main | GLU | 247 | Side | 24,45% | TRP | 313 | Side | PHE | 269 | Main | 20,90% |
| **ASN** | 307 | Main | GLY | 317 | Main | 24,45% | THR | 155 | Main | TRP | 151 | Main | 20,75% |
| **ILE** | 184 | Main | ASP | 182 | Side | 24,30% | LEU | 243 | Main | VAL | 334 | Main | 20,65% |
| **LEU** | 260 | Main | LEU | 291 | Main | 23,05% | SER | 180 | Side | PRO | 224 | Main | 20,50% |
| **ASN** | 295 | Main | TYR | 302 | Main | 22,85% | VAL | 130 | Main | TYR | 294 | Main | 20,25% |
| **HIP** | 287 | Side | CYM | 150 | Side | 21,75% | LEU | 290 | Main | LYS | 306 | Main | 20,15% |
| **TYR** | 294 | Main | VAL | 130 | Main | 21,15% | SER | 154 | Side | ILE | 262 | Main | 20,15% |
| **SER** | 335 | Side | ASP | 242 | Side | 20,95% | LEU | 277 | Main | ARG | 320 | Main | 20,05% |
| **PHE** | 347 | Main | GLH | 343 | Main | 20,85% | ALA | 252 | Main | ASP | 248 | Main | 19,65% |
| **SER** | 149 | Side | TYR | 216 | Side | 20,75% | TRP | 199 | Main | ASN | 195 | Main | 18,80% |
| **GLN** | 144 | Main | SER | 308 | Main | 20,35% | HIP | 231 | Side | GLN | 227 | Main | 18,70% |
| **GLY** | 221 | Main | SER | 218 | Main | 19,95% | GLN | 146 | Main | GLY | 219 | Main | 18,50% |
| **VAL** | 130 | Main | TYR | 294 | Main | 19,85% | GLY | 221 | Main | SER | 218 | Main | 18,35% |
| **ALA** | 252 | Main | ASP | 248 | Main | 19,50% | THR | 210 | Main | GLH | 232 | Main | 18,15% |
| **THR** | 155 | Main | TRP | 151 | Main | 19,15% | SER | 154 | Side | VAL | 289 | Main | 17,85% |
| **SER** | 336 | Side | ALA | 337 | Main | 18,95% | ASN | 307 | Main | GLY | 317 | Main | 17,70% |
| **TRP** | 151 | Side | GLY | 187 | Main | 18,55% | TRP | 151 | Side | GLY | 187 | Main | 17,40% |
| **CYX** | 188 | Main | GLN | 223 | Side | 18,45% | TRP | 132 | Side | LEU | 254 | Main | 17,10% |
| **VAL** | 201 | Main | PHE | 197 | Main | 18,45% | SER | 298 | Main | ASN | 295 | Side | 16,90% |
| **GLH** | 343 | Side | ASP | 239 | Side | 18,20% | TYR | 294 | Main | VAL | 130 | Main | 16,80% |

| | | | | | | | | | | | | | |
|---|---|---|---|---|---|---|---|---|---|---|---|---|---|
| **GLN** | 223 | Side | VAL | 179 | Main | 18,10% | SER | 149 | Side | TYR | 216 | Side | 16,40% |
| **LYS** | 142 | Side | GLH | 175 | Side | 17,85% | ASN | 158 | Side | PRO | 259 | Main | 15,95% |
| **GLU** | 315 | Main | TYR | 318 | Main | 17,75% | ALA | 266 | Main | ASH | 286 | Side | 15,95% |
| **GLY** | 157 | Main | PHE | 153 | Main | 17,40% | LEU | 192 | Main | ASH | 185 | Side | 15,85% |
| **LYS** | 142 | Side | GLU | 211 | Side | 17,20% | GLN | 162 | Side | PRO | 259 | Main | 15,45% |
| **LEU** | 192 | Main | ASH | 185 | Side | 16,65% | LYS | 142 | Side | GLU | 160 | Side | 15,30% |
| **ASN** | 158 | Side | PRO | 259 | Main | 16,65% | SER | 308 | Side | GLN | 144 | Side | 15,20% |
| **GLN** | 146 | Main | GLY | 219 | Main | 16,65% | LYS | 142 | Side | GLU | 211 | Side | 14,75% |
| **LEU** | 290 | Main | LYS | 306 | Main | 16,55% | PHE | 186 | Main | GLY | 190 | Main | 14,65% |
| **SER** | 298 | Main | ASN | 295 | Side | 16,10% | GLU | 315 | Main | TYR | 318 | Main | 14,35% |
| **GLN** | 162 | Side | PRO | 259 | Main | 15,60% | LYS | 306 | Side | THR | 139 | Main | 14,30% |
| **ARG** | 133 | Main | ASP | 131 | Side | 15,35% | VAL | 201 | Main | PHE | 197 | Main | 13,75% |
| **TRP** | 132 | Side | LEU | 254 | Main | 15,35% | SER | 336 | Side | ALA | 337 | Main | 13,55% |
| **ASH** | 185 | Main | ASP | 182 | Side | 15,05% | GLH | 232 | Main | SER | 213 | Side | 13,50% |
| **GLH** | 134 | Side | ASP | 131 | Side | 14,75% | ALA | 255 | Main | ALA | 251 | Main | 13,50% |
| **ILE** | 319 | Main | ILE | 305 | Main | 14,50% | GLN | 223 | Side | VAL | 179 | Main | 13,15% |
| **ASP** | 265 | Main | CYX | 328 | Main | 14,05% | ILE | 200 | Main | ALA | 196 | Main | 13,05% |
| **TRP** | 199 | Main | ASN | 195 | Main | 13,45% | TYR | 214 | Main | THR | 210 | Main | 12,50% |
| **PHE** | 269 | Main | ALA | 266 | Main | 13,40% | ALA | 196 | Main | LEU | 192 | Main | 12,20% |
| **ALA** | 255 | Main | ALA | 251 | Main | 13,30% | SER | 203 | Main | TRP | 199 | Main | 12,15% |
| **HIP** | 231 | Side | GLN | 227 | Main | 13,20% | ASN | 202 | Side | ASN | 198 | Main | 11,75% |
| **LYS** | 306 | Side | PRO | 140 | Main | 13,15% | ASP | 182 | Main | LEU | 178 | Main | 11,70% |
| **LYS** | 323 | Side | ASP | 248 | Side | 12,65% | GLY | 157 | Main | PHE | 153 | Main | 11,60% |
| **ASN** | 198 | Main | ASH | 194 | Main | 12,40% | ILE | 319 | Main | ILE | 305 | Main | 11,30% |
| **ILE** | 200 | Main | ALA | 196 | Main | 12,25% | ASH | 185 | Main | ASP | 182 | Side | 11,25% |
| **SER** | 172 | Side | GLU | 211 | Side | 12,15% | LYS | 306 | Side | PRO | 140 | Main | 11,15% |
| **GLH** | 232 | Main | SER | 213 | Side | 11,75% | ASN | 207 | Main | ASN | 204 | Main | 11,05% |
| **TYR** | 253 | Main | ALA | 249 | Main | 11,35% | LYS | 135 | Main | ASP | 131 | Main | 11,00% |
| **GLU** | 222 | Main | ASN | 220 | Side | 11,35% | GLY | 145 | Main | ASH | 143 | Side | 10,75% |
| **ASN** | 207 | Main | ASN | 204 | Main | 11,05% | GLH | 256 | Side | ALA | 252 | Main | 10,70% |
| **ASN** | 202 | Side | ASN | 198 | Main | 10,95% | TYR | 253 | Main | ALA | 249 | Main | 10,60% |
| **TYR** | 346 | Main | GLH | 343 | Main | 10,90% | ASP | 248 | Main | ASP | 246 | Side | 10,05% |
| **ALA** | 137 | Main | TRP | 132 | Main | 10,80% | ASN | 257 | Side | TYR | 253 | Main | 9,85% |
| **LEU** | 345 | Main | PRO | 342 | Main | 10,80% | ARG | 133 | Main | ASP | 131 | Side | 9,80% |
| **LYS** | 306 | Side | THR | 139 | Main | 10,00% | SER | 335 | Side | ASP | 242 | Side | 9,80% |
| **ASP** | 248 | Main | ASP | 246 | Side | 9,95% | ALA | 261 | Main | SER | 335 | Main | 9,75% |
| **GLY** | 189 | Main | PHE | 186 | Main | 9,90% | LYS | 323 | Side | ASP | 248 | Side | 9,65% |
| **GLY** | 317 | Main | TRP | 313 | Main | 9,80% | PHE | 269 | Main | ALA | 266 | Main | 9,50% |
| **ASP** | 182 | Main | LEU | 178 | Main | 9,75% | GLU | 222 | Main | ASN | 220 | Side | 9,45% |
| **HIP** | 287 | Main | VAL | 264 | Main | 9,70% | LYS | 142 | Side | GLH | 175 | Side | 9,35% |
| **TYR** | 302 | Main | ASN | 295 | Main | 9,60% | TYR | 302 | Main | ASN | 295 | Main | 9,30% |
| **TYR** | 214 | Main | THR | 210 | Main | 9,35% | ASN | 198 | Main | ASH | 194 | Main | 9,25% |
| **GLN** | 348 | Side | ASN | 344 | Main | 9,20% | PHE | 347 | Main | ASN | 344 | Main | 9,20% |
| **GLY** | 206 | Main | ILE | 200 | Main | 8,95% | ASP | 265 | Main | CYX | 328 | Main | 8,80% |
| **ASN** | 207 | Side | ASN | 204 | Main | 8,40% | THR | 281 | Main | SER | 268 | Side | 8,55% |
| **ASH** | 286 | Side | ASP | 265 | Side | 8,25% | TYR | 346 | Main | GLH | 343 | Main | 8,40% |
| **ASN** | 229 | Main | GLN | 227 | Side | 8,10% | ARG | 133 | Side | ASP | 131 | Side | 8,10% |

| | | | | | | | | | | | | | |
|---|---|---|---|---|---|---|---|---|---|---|---|---|---|
| **SER** | 203 | Main | TRP | 199 | Main | 7,95% | SER | 154 | Side | CYM | 150 | Main | 7,85% |
| **VAL** | 179 | Main | GLH | 175 | Main | 7,85% | TYR | 318 | Main | GLU | 315 | Main | 7,60% |
| **SER** | 154 | Side | ILE | 262 | Main | 7,85% | ASN | 344 | Side | ASN | 257 | Side | 7,50% |
| **GLH** | 343 | Side | HIP | 240 | Main | 7,80% | GLY | 189 | Main | PHE | 186 | Main | 7,40% |
| **GLN** | 245 | Side | ASN | 331 | Side | 7,70% | GLN | 227 | Side | ASN | 229 | Main | 7,35% |
| **TYR** | 318 | Main | GLU | 315 | Main | 7,30% | SER | 310 | Main | LYS | 142 | Main | 7,30% |
| **SER** | 310 | Side | TRP | 309 | Main | 7,05% | GLY | 317 | Main | TRP | 313 | Main | 7,30% |
| **GLN** | 164 | Side | ASN | 168 | Main | 6,90% | ALA | 251 | Main | GLU | 247 | Main | 7,20% |
| **GLY** | 288 | Main | CYM | 150 | Side | 6,90% | ASN | 195 | Side | ASH | 185 | Side | 7,15% |
| **PHE** | 186 | Main | GLY | 190 | Main | 6,85% | GLN | 348 | Side | ASN | 344 | Main | 7,15% |
| **LYS** | 323 | Side | ASH | 296 | Side | 6,80% | GLY | 206 | Main | ILE | 200 | Main | 7,10% |
| **SER** | 335 | Side | ALA | 261 | Main | 6,75% | GLN | 348 | Side | GLH | 256 | Main | 7,05% |
| **THR** | 281 | Main | SER | 268 | Side | 6,55% | GLN | 332 | Side | ASN | 331 | Side | 7,00% |
| **LYS** | 142 | Side | SER | 172 | Side | 6,50% | LYS | 142 | Side | SER | 172 | Side | 6,85% |
| **ASN** | 229 | Side | GLN | 227 | Side | 6,45% | GLN | 245 | Side | ASN | 331 | Side | 6,80% |
| **ALA** | 261 | Main | SER | 335 | Main | 6,35% | ASN | 207 | Side | ASN | 204 | Main | 6,65% |
| **ASN** | 331 | Side | GLU | 247 | Side | 6,35% | VAL | 179 | Main | GLH | 175 | Main | 6,50% |
| **THR** | 325 | Side | GLH | 322 | Side | 6,25% | SER | 218 | Main | GLN | 176 | Side | 6,45% |
| **TYR** | 346 | Side | ASN | 257 | Side | 6,25% | SER | 310 | Side | TRP | 309 | Main | 6,40% |
| **GLY** | 145 | Main | ASH | 143 | Side | 6,00% | GLN | 348 | Side | ASN | 257 | Side | 6,15% |
| **GLN** | 332 | Side | LEU | 329 | Main | 5,85% | TYR | 216 | Main | GLU | 211 | Side | 5,75% |
| **MET** | 330 | Main | GLN | 327 | Main | 5,80% | HIP | 231 | Side | SER | 213 | Main | 5,55% |
| **ASN** | 295 | Side | TYR | 302 | Side | 5,30% | LEU | 345 | Main | GLH | 343 | Side | 5,55% |
| **ALA** | 251 | Main | GLU | 247 | Main | 5,20% | LYS | 323 | Side | ASH | 296 | Side | 5,35% |
| **ASN** | 257 | Side | TYR | 253 | Main | 5,05% | SER | 180 | Side | GLN | 176 | Main | 5,35% |
| | | | | | | | ASN | 295 | Side | TYR | 302 | Side | 5,10% |
| | | | | | | | ALA | 333 | Main | MET | 330 | Main | 5,05% |
| | | | | | | | GLH | 134 | Main | ASP | 131 | Side | 5,05% |
| | | | | | | | GLN | 146 | Side | ASN | 220 | Main | 5,00% |

**Table S4b: Relative occurrence of hydrogen bonds within the catalytic domain.** Four respective MD simulations at pH 4 (pH4_3 and pH4_4, see Table S4a for pH4_1 and pH4_2) were carried out. `Main' refers to interactions of the backbone. 'Side' indicates interactions of side chains. Occurrences below 5% were left out for clarity.

| pH4_3 | | | | | | | pH4_4 | | | | | | |
|---|---|---|---|---|---|---|---|---|---|---|---|---|---|
| donor | | | acceptor | | | occupancy | donor | | | acceptor | | | occupancy |
| ASH | 185 | Side | ASP | 182 | Side | 77,36% | ASH | 222 | Side | ASP | 182 | Side | 82,91% |
| THR | 267 | Side | ASP | 265 | Side | 73,16% | THR | 304 | Side | ASP | 265 | Side | 78,21% |
| THR | 155 | Side | TRP | 151 | Main | 72,96% | TYR | 309 | Side | GLY | 275 | Main | 77,21% |
| ASH | 194 | Side | ASP | 242 | Side | 70,36% | ARG | 357 | Side | GLU | 315 | Side | 76,61% |
| TYR | 272 | Side | GLY | 275 | Main | 69,07% | THR | 192 | Side | TRP | 151 | Main | 72,51% |
| TRP | 303 | Main | ILE | 321 | Main | 68,62% | TRP | 340 | Main | ILE | 321 | Main | 67,42% |
| ARG | 320 | Side | GLU | 315 | Side | 64,77% | ASH | 231 | Side | ASP | 242 | Side | 64,32% |
| GLH | 283 | Main | ASP | 265 | Side | 61,02% | ALA | 303 | Main | ASH | 286 | Side | 60,57% |
| LYS | 306 | Main | LEU | 290 | Main | 60,77% | GLH | 320 | Main | ASP | 265 | Side | 56,92% |
| THR | 139 | Side | GLU | 160 | Side | 58,47% | LYS | 343 | Main | LEU | 290 | Main | 56,57% |
| ARG | 133 | Side | ASP | 131 | Side | 56,82% | ILE | 193 | Main | ALA | 152 | Main | 56,42% |

| | | | | | | | | | | | | | |
|---|---|---|---|---|---|---|---|---|---|---|---|---|---|
| PHE | 197 | Main | MET | 193 | Main | 56,47% | PHE | 234 | Main | MET | 193 | Main | 56,12% |
| VAL | 241 | Main | SER | 336 | Main | 56,22% | VAL | 278 | Main | SER | 336 | Main | 55,17% |
| TYR | 216 | Side | GLN | 144 | Main | 55,47% | SER | 211 | Side | TYR | 214 | Main | 53,82% |
| SER | 174 | Main | PHE | 209 | Main | 55,37% | ARG | 170 | Side | ASP | 131 | Side | 52,77% |
| GLH | 175 | Side | GLU | 211 | Side | 53,97% | TYR | 253 | Side | GLN | 144 | Main | 52,32% |
| SER | 174 | Side | TYR | 214 | Main | 53,12% | GLN | 199 | Main | ASN | 158 | Main | 52,22% |
| HIP | 287 | Side | ASN | 307 | Side | 52,77% | SER | 211 | Main | PHE | 209 | Main | 51,42% |
| ILE | 156 | Main | ALA | 152 | Main | 51,72% | TYR | 331 | Side | PRO | 127 | Main | 50,62% |
| SER | 298 | Side | PRO | 300 | Main | 51,27% | ASH | 333 | Main | ALA | 128 | Main | 50,27% |
| ALA | 337 | Main | ASN | 158 | Side | 51,22% | CYX | 317 | Main | GLN | 327 | Side | 50,22% |
| LEU | 173 | Main | GLU | 160 | Side | 49,08% | HIP | 324 | Side | ASN | 307 | Side | 50,12% |
| VAL | 217 | Main | GLN | 176 | Side | 48,28% | ASH | 323 | Side | ASP | 265 | Side | 49,08% |
| GLN | 144 | Side | CYX | 147 | Main | 48,13% | GLY | 312 | Main | GLU | 315 | Side | 48,53% |
| SER | 282 | Side | CYX | 280 | Main | 45,38% | SER | 335 | Side | PRO | 300 | Main | 48,48% |
| ASH | 296 | Main | ALA | 128 | Main | 45,28% | LEU | 328 | Main | LEU | 260 | Main | 48,23% |
| GLN | 162 | Main | ASN | 158 | Main | 44,13% | TRP | 169 | Main | VAL | 292 | Main | 47,23% |
| CYX | 280 | Main | GLN | 327 | Side | 43,88% | SER | 250 | Side | THR | 210 | Side | 46,58% |
| ASN | 168 | Side | TRP | 163 | Main | 43,83% | VAL | 254 | Main | GLN | 176 | Side | 46,28% |
| GLU | 160 | Main | ILE | 156 | Main | 43,68% | ILE | 274 | Main | GLY | 206 | Main | 45,53% |
| GLU | 211 | Main | SER | 172 | Main | 42,83% | VAL | 326 | Main | ILE | 262 | Main | 44,33% |
| LEU | 291 | Main | LEU | 260 | Main | 42,38% | ASN | 205 | Side | TRP | 163 | Main | 44,28% |
| ASH | 286 | Main | VAL | 264 | Main | 42,18% | GLU | 197 | Main | ILE | 156 | Main | 44,23% |
| THR | 238 | Main | VAL | 338 | Main | 42,03% | LYS | 360 | Main | PRO | 301 | Main | 43,98% |
| SER | 213 | Side | THR | 210 | Side | 41,83% | THR | 275 | Main | VAL | 338 | Main | 43,48% |
| GLY | 275 | Main | GLU | 315 | Side | 41,58% | ALA | 374 | Main | ASN | 158 | Side | 42,63% |
| ILE | 250 | Main | ASP | 246 | Main | 40,58% | SER | 255 | Side | GLU | 222 | Main | 41,93% |
| LYS | 323 | Main | PRO | 301 | Main | 40,48% | GLH | 359 | Main | LEU | 277 | Main | 41,93% |
| SER | 218 | Side | GLU | 222 | Main | 40,48% | GLN | 181 | Side | CYX | 147 | Main | 41,83% |
| ILE | 237 | Main | GLY | 206 | Main | 39,93% | VAL | 202 | Main | GLY | 161 | Main | 41,68% |
| GLH | 322 | Main | LEU | 277 | Main | 39,73% | GLY | 271 | Main | VAL | 208 | Main | 41,43% |
| GLY | 234 | Main | VAL | 208 | Main | 39,53% | ASN | 195 | Side | SER | 336 | Side | 39,18% |
| SER | 154 | Main | CYM | 150 | Main | 39,43% | ILE | 287 | Main | ASP | 246 | Main | 38,73% |
| GLY | 324 | Main | GLU | 247 | Side | 38,43% | LEU | 291 | Main | ILE | 250 | Main | 38,68% |
| SER | 180 | Main | GLN | 176 | Main | 37,93% | THR | 176 | Side | GLU | 160 | Side | 37,98% |
| ALA | 266 | Main | ASH | 286 | Side | 37,88% | SER | 345 | Side | GLN | 144 | Side | 37,38% |
| VAL | 289 | Main | ILE | 262 | Main | 37,28% | VAL | 301 | Main | HIP | 287 | Main | 36,98% |
| VAL | 264 | Main | HIP | 287 | Main | 36,98% | ASH | 323 | Main | VAL | 264 | Main | 36,93% |
| ASN | 202 | Main | ASN | 198 | Main | 36,93% | ASN | 239 | Main | ASN | 198 | Main | 36,13% |
| THR | 267 | Main | ASP | 265 | Side | 36,08% | GLN | 199 | Side | ASN | 257 | Main | 35,33% |
| TRP | 303 | Side | GLU | 247 | Side | 35,38% | LEU | 210 | Main | GLU | 160 | Side | 34,18% |
| LEU | 254 | Main | ILE | 250 | Main | 35,08% | ILE | 341 | Main | GLY | 293 | Main | 34,13% |
| ILE | 305 | Main | ILE | 319 | Main | 33,68% | SER | 191 | Main | CYM | 150 | Main | 34,03% |
| ILE | 304 | Main | GLY | 293 | Main | 32,63% | GLN | 213 | Side | SER | 218 | Side | 33,73% |
| ASN | 158 | Side | PRO | 259 | Main | 32,58% | SER | 217 | Main | GLN | 176 | Main | 33,53% |
| TRP | 132 | Main | VAL | 292 | Main | 32,23% | TRP | 200 | Main | ILE | 159 | Main | 32,63% |
| SER | 154 | Side | VAL | 289 | Main | 32,13% | TRP | 340 | Side | GLU | 247 | Side | 32,48% |
| SER | 335 | Main | ALA | 261 | Main | 31,18% | GLU | 248 | Main | SER | 172 | Main | 32,33% |

| | | | | | | | | | | | | | |
|---|---|---|---|---|---|---|---|---|---|---|---|---|---|
| VAL | 292 | Main | ILE | 304 | Main | 31,13% | ASN | 195 | Side | PRO | 259 | Main | 32,28% |
| SER | 180 | Side | PRO | 224 | Main | 31,13% | GLY | 361 | Main | GLU | 247 | Side | 31,53% |
| SER | 336 | Side | ALA | 337 | Main | 31,03% | SER | 319 | Side | CYX | 280 | Main | 30,68% |
| ASN | 257 | Main | TYR | 253 | Main | 30,28% | ASN | 241 | Side | ASN | 207 | Main | 30,48% |
| ASN | 273 | Side | TYR | 272 | Main | 28,74% | GLN | 181 | Main | SER | 308 | Main | 30,33% |
| ASN | 204 | Side | ASN | 207 | Main | 28,59% | ILE | 342 | Main | ILE | 319 | Main | 30,23% |
| GLN | 284 | Main | ASP | 265 | Side | 28,54% | ASN | 294 | Main | TYR | 253 | Main | 29,74% |
| GLN | 176 | Side | SER | 218 | Side | 28,34% | LEU | 280 | Main | VAL | 334 | Main | 29,19% |
| VAL | 338 | Main | ASP | 239 | Main | 27,99% | VAL | 167 | Main | TYR | 294 | Main | 28,49% |
| ASN | 331 | Main | ASN | 326 | Side | 27,99% | SER | 209 | Side | GLU | 160 | Side | 28,49% |
| LEU | 277 | Main | ARG | 320 | Main | 27,84% | ASN | 368 | Side | GLN | 245 | Main | 28,44% |
| GLN | 144 | Main | SER | 308 | Main | 27,74% | LYS | 179 | Side | GLU | 211 | Side | 27,39% |
| ASN | 158 | Side | SER | 336 | Side | 27,69% | VAL | 245 | Main | ALA | 235 | Main | 27,34% |
| SER | 308 | Side | GLN | 144 | Side | 27,69% | SER | 372 | Main | ALA | 261 | Main | 27,09% |
| THR | 155 | Main | TRP | 151 | Main | 27,19% | VAL | 375 | Main | ASP | 239 | Main | 26,19% |
| TYR | 294 | Main | VAL | 130 | Main | 26,94% | ASN | 332 | Main | TYR | 302 | Main | 25,59% |
| SER | 335 | Side | ASP | 242 | Side | 26,19% | PHE | 306 | Main | ALA | 266 | Main | 25,59% |
| HIP | 287 | Main | VAL | 264 | Main | 26,09% | THR | 304 | Main | ASP | 265 | Side | 25,19% |
| HIP | 287 | Side | CYM | 150 | Side | 25,29% | ILE | 221 | Main | ASP | 182 | Side | 24,99% |
| GLN | 327 | Side | THR | 325 | Side | 25,09% | GLN | 213 | Side | SER | 180 | Side | 23,49% |
| TRP | 163 | Main | ILE | 159 | Main | 23,99% | TYR | 253 | Main | GLU | 211 | Side | 22,59% |
| CYX | 188 | Main | GLN | 223 | Side | 23,89% | VAL | 329 | Main | ILE | 304 | Main | 22,24% |
| ASN | 326 | Main | GLU | 247 | Side | 23,79% | LEU | 314 | Main | ARG | 320 | Main | 21,94% |
| GLN | 146 | Main | GLY | 219 | Main | 23,74% | SER | 191 | Side | CYM | 150 | Main | 21,44% |
| GLN | 176 | Side | SER | 180 | Side | 23,69% | ALA | 289 | Main | ASP | 248 | Main | 21,34% |
| VAL | 208 | Main | ALA | 235 | Main | 23,29% | PHE | 223 | Main | GLY | 190 | Main | 21,14% |
| VAL | 165 | Main | GLY | 161 | Main | 23,19% | GLY | 377 | Main | ALA | 236 | Main | 21,04% |
| LEU | 260 | Main | LEU | 291 | Main | 22,54% | CYX | 225 | Main | GLN | 223 | Side | 20,94% |
| LYS | 142 | Side | GLU | 160 | Side | 22,39% | TYR | 331 | Main | VAL | 130 | Main | 20,84% |
| ILE | 184 | Main | ASP | 182 | Side | 22,29% | GLY | 354 | Main | TRP | 313 | Main | 20,64% |
| PHE | 186 | Main | GLY | 190 | Main | 22,29% | SER | 372 | Side | ASP | 242 | Side | 20,59% |
| ALA | 255 | Main | ALA | 251 | Main | 21,89% | GLU | 352 | Main | TYR | 318 | Main | 20,39% |
| ASN | 307 | Main | GLY | 317 | Main | 21,09% | ASN | 344 | Main | GLY | 317 | Main | 20,29% |
| LYS | 306 | Side | PRO | 140 | Main | 20,59% | SER | 373 | Main | VAL | 241 | Main | 20,24% |
| GLY | 221 | Main | SER | 218 | Main | 20,39% | SER | 186 | Side | TYR | 216 | Side | 20,04% |
| LEU | 243 | Main | VAL | 334 | Main | 20,24% | ASP | 302 | Main | CYX | 328 | Main | 19,94% |
| ASN | 295 | Main | TYR | 302 | Main | 19,79% | ALA | 174 | Main | TRP | 132 | Main | 19,49% |
| TRP | 199 | Main | ASN | 195 | Main | 19,49% | ASN | 310 | Side | TYR | 272 | Main | 19,24% |
| HIP | 231 | Side | GLN | 227 | Main | 19,24% | HIP | 268 | Side | GLN | 227 | Main | 19,09% |
| ALA | 137 | Main | TRP | 132 | Main | 18,89% | SER | 217 | Side | GLN | 176 | Main | 18,89% |
| SER | 154 | Side | ILE | 262 | Main | 18,84% | TRP | 188 | Side | GLY | 187 | Main | 18,84% |
| SER | 336 | Main | VAL | 241 | Main | 18,64% | GLN | 321 | Main | ASP | 265 | Side | 18,44% |
| ALA | 252 | Main | ASP | 248 | Main | 18,59% | TYR | 290 | Main | ALA | 249 | Main | 18,39% |
| ASN | 257 | Side | GLN | 348 | Side | 18,54% | TRP | 236 | Main | ASN | 195 | Main | 18,34% |
| LEU | 192 | Main | ASH | 185 | Side | 18,39% | LYS | 343 | Side | THR | 139 | Main | 18,19% |
| GLY | 340 | Main | ALA | 236 | Main | 18,19% | GLN | 364 | Side | THR | 325 | Side | 18,04% |
| THR | 210 | Main | GLH | 232 | Main | 18,19% | ASN | 363 | Main | GLU | 247 | Side | 17,94% |

| | | | | | | | | | | | | | |
|---|---|---|---|---|---|---|---|---|---|---|---|---|---|
| ALA | 196 | Main | LEU | 192 | Main | 18,04% | SER | 191 | Side | VAL | 289 | Main | 17,29% |
| TYR | 253 | Main | ALA | 249 | Main | 17,79% | GLN | 183 | Main | GLY | 219 | Main | 16,94% |
| ILE | 200 | Main | ALA | 196 | Main | 16,99% | TRP | 350 | Side | PHE | 269 | Main | 16,79% |
| VAL | 130 | Main | TYR | 294 | Main | 16,49% | GLY | 258 | Main | SER | 218 | Main | 16,64% |
| TRP | 151 | Side | GLY | 187 | Main | 16,04% | ILE | 356 | Main | ILE | 305 | Main | 16,54% |
| VAL | 201 | Main | PHE | 197 | Main | 15,99% | THR | 247 | Main | GLH | 232 | Main | 16,49% |
| TRP | 313 | Side | PHE | 269 | Main | 15,89% | LEU | 229 | Main | ASH | 185 | Side | 16,44% |
| GLN | 162 | Side | ASN | 257 | Main | 15,79% | ARG | 170 | Main | ASP | 131 | Side | 16,44% |
| SER | 149 | Side | TYR | 216 | Side | 15,64% | THR | 192 | Main | TRP | 151 | Main | 15,99% |
| SER | 298 | Main | ASN | 295 | Side | 14,69% | HIP | 324 | Side | CYM | 150 | Side | 15,29% |
| PHE | 347 | Main | ASN | 344 | Main | 14,19% | LEU | 297 | Main | LEU | 291 | Main | 15,24% |
| GLU | 315 | Main | TYR | 318 | Main | 14,04% | VAL | 238 | Main | PHE | 197 | Main | 14,99% |
| LEU | 290 | Main | LYS | 306 | Main | 14,04% | ASN | 363 | Side | GLU | 247 | Side | 14,44% |
| GLH | 232 | Main | SER | 213 | Side | 13,74% | ILE | 299 | Main | VAL | 289 | Main | 13,99% |
| LYS | 142 | Side | GLU | 211 | Side | 13,74% | GLN | 260 | Side | VAL | 179 | Main | 13,89% |
| ASH | 185 | Main | ASP | 182 | Side | 13,64% | ALA | 292 | Main | ALA | 251 | Main | 13,59% |
| LYS | 323 | Side | ASP | 248 | Side | 13,49% | SER | 191 | Side | ILE | 262 | Main | 13,24% |
| GLN | 348 | Side | ASN | 257 | Side | 13,24% | SER | 217 | Side | PRO | 224 | Main | 13,19% |
| GLN | 223 | Side | VAL | 179 | Main | 13,19% | TYR | 251 | Main | THR | 210 | Main | 13,19% |
| TYR | 214 | Main | THR | 210 | Main | 12,59% | SER | 335 | Main | ASN | 295 | Side | 13,09% |
| ARG | 133 | Main | ASP | 131 | Side | 12,54% | GLH | 269 | Main | SER | 213 | Side | 12,99% |
| ASN | 195 | Side | ASH | 185 | Side | 11,49% | TRP | 169 | Side | LEU | 254 | Main | 12,99% |
| GLY | 145 | Main | ASH | 143 | Side | 11,44% | GLY | 182 | Main | ASH | 143 | Side | 12,54% |
| TRP | 132 | Side | LEU | 254 | Main | 11,34% | GLN | 369 | Main | LEU | 329 | Main | 12,44% |
| ASN | 207 | Main | ASN | 204 | Main | 11,19% | ASN | 239 | Side | ASN | 198 | Main | 12,14% |
| LYS | 142 | Side | SER | 172 | Side | 10,79% | ILE | 237 | Main | ALA | 196 | Main | 12,04% |
| TYR | 216 | Main | GLU | 211 | Side | 10,74% | ASH | 222 | Main | ASP | 182 | Side | 11,69% |
| THR | 325 | Side | GLH | 322 | Side | 10,69% | MET | 367 | Main | GLN | 327 | Main | 11,69% |
| GLN | 162 | Side | PRO | 259 | Main | 10,54% | SER | 347 | Side | TRP | 309 | Main | 11,59% |
| ASN | 198 | Main | ASH | 194 | Main | 10,49% | ASN | 244 | Main | ASN | 204 | Main | 11,44% |
| GLY | 157 | Main | PHE | 153 | Main | 10,34% | ALA | 233 | Main | LEU | 192 | Main | 11,19% |
| TYR | 302 | Main | ASN | 295 | Main | 10,29% | HIP | 324 | Main | VAL | 264 | Main | 10,84% |
| GLH | 175 | Side | GLU | 160 | Side | 10,24% | SER | 240 | Main | TRP | 199 | Main | 10,54% |
| ASN | 202 | Side | ASN | 198 | Main | 10,24% | PHE | 384 | Main | ASN | 344 | Main | 10,19% |
| PHE | 269 | Main | ALA | 266 | Main | 10,14% | GLN | 264 | Side | ASN | 229 | Main | 10,00% |
| GLY | 206 | Main | ILE | 200 | Main | 9,90% | THR | 318 | Main | SER | 268 | Side | 9,75% |
| LYS | 306 | Side | THR | 139 | Main | 9,70% | TYR | 383 | Main | GLH | 343 | Main | 9,60% |
| GLY | 317 | Main | TRP | 313 | Main | 9,50% | TYR | 339 | Main | ASN | 295 | Main | 9,55% |
| LYS | 142 | Side | GLH | 175 | Side | 9,15% | GLN | 369 | Side | ASN | 331 | Side | 9,55% |
| ASN | 344 | Side | ASN | 257 | Side | 8,85% | ALA | 288 | Main | GLU | 247 | Main | 9,50% |
| SER | 203 | Main | TRP | 199 | Main | 8,45% | ASP | 219 | Main | LEU | 178 | Main | 9,50% |
| GLU | 222 | Main | ASN | 220 | Side | 8,40% | ASN | 235 | Main | ASH | 194 | Main | 9,45% |
| ASP | 248 | Main | ASP | 246 | Side | 8,40% | ASN | 381 | Side | GLY | 167 | Main | 9,35% |
| ASN | 257 | Side | TYR | 253 | Main | 8,35% | LYS | 360 | Side | ASH | 296 | Side | 9,20% |
| GLN | 227 | Side | ASN | 229 | Main | 8,35% | GLY | 194 | Main | PHE | 153 | Main | 9,15% |
| ILE | 319 | Main | ILE | 305 | Main | 8,05% | ASN | 368 | Main | ASN | 326 | Side | 8,55% |
| ASP | 265 | Main | CYX | 328 | Main | 8,00% | ASP | 285 | Main | ASP | 246 | Side | 8,15% |

| | | | | | | | | | | | | | |
|---|---|---|---|---|---|---|---|---|---|---|---|---|---|
| LYS | 323 | Side | ASH | 296 | Side | 7,85% | VAL | 216 | Main | GLH | 175 | Main | 7,90% |
| VAL | 179 | Main | GLH | 175 | Main | 7,60% | GLY | 243 | Main | ILE | 200 | Main | 7,85% |
| TYR | 318 | Main | GLU | 315 | Main | 7,30% | LEU | 327 | Main | LYS | 306 | Main | 6,95% |
| GLY | 189 | Main | PHE | 186 | Main | 7,25% | GLU | 259 | Main | ASN | 220 | Side | 6,90% |
| ASN | 331 | Side | GLU | 247 | Side | 7,15% | ALA | 298 | Main | SER | 335 | Main | 6,75% |
| ASP | 182 | Main | LEU | 178 | Main | 6,95% | TYR | 309 | Main | PHE | 269 | Main | 6,60% |
| HIP | 240 | Side | ASP | 242 | Side | 6,85% | SER | 373 | Side | ALA | 337 | Main | 6,50% |
| MET | 330 | Main | GLN | 327 | Main | 6,85% | ASN | 244 | Side | ASN | 204 | Main | 6,40% |
| SER | 310 | Side | TRP | 309 | Main | 6,80% | THR | 362 | Side | GLH | 322 | Side | 6,30% |
| ALA | 333 | Main | MET | 330 | Main | 6,80% | LYS | 172 | Main | ASP | 131 | Main | 6,10% |
| SER | 172 | Side | GLU | 160 | Side | 6,65% | ASN | 294 | Side | TYR | 253 | Main | 6,10% |
| THR | 281 | Main | SER | 268 | Side | 6,55% | ASN | 232 | Side | ASH | 185 | Side | 5,95% |
| ASN | 207 | Side | ASN | 204 | Main | 5,95% | GLY | 226 | Main | PHE | 186 | Main | 5,95% |
| GLH | 256 | Side | GLN | 348 | Side | 5,95% | ASN | 266 | Side | GLN | 227 | Side | 5,85% |
| CYX | 181 | Main | MET | 177 | Main | 5,85% | LYS | 179 | Side | GLH | 175 | Side | 5,75% |
| ASN | 295 | Side | TYR | 302 | Side | 5,50% | SER | 255 | Side | GLN | 176 | Side | 5,75% |
| SER | 218 | Main | GLN | 176 | Side | 5,30% | SER | 372 | Side | ALA | 261 | Main | 5,70% |
| GLN | 245 | Side | ASN | 331 | Side | 5,30% | ASN | 344 | Side | TRP | 313 | Side | 5,45% |
| ASN | 344 | Side | GLH | 256 | Main | 5,25% | ASN | 266 | Main | GLN | 227 | Side | 5,35% |
| SER | 180 | Side | GLN | 176 | Main | 5,10% | GLH | 293 | Side | GLN | 348 | Main | 5,35% |
| GLN | 348 | Side | GLN | 348 | Side | 5,10% | LEU | 382 | Main | ALA | 166 | Main | 5,30% |
| | | | | | | | SER | 255 | Main | GLN | 176 | Side | 5,25% |
| | | | | | | | ASN | 381 | Side | ASN | 168 | Side | 5,15% |
| | | | | | | | HIP | 268 | Side | SER | 213 | Main | 5,10% |

**Table S5a: Relative occurrence of hydrogen bonds within the catalytic domain.** Four respective MD simulations at pH 8 (pH8_1 and pH8_2, see Table S5b for pH8_3 and pH8_4) were carried out. `Main' refers to interactions of the backbone. 'Side' indicates interactions of side chains. Occurrences below 5% were left out for clarity.

| pH8_1 | | | | | | | pH8_2 | | | | | | |
|---|---|---|---|---|---|---|---|---|---|---|---|---|---|
| donor | | | acceptor | | | occupancy | donor | | | acceptor | | | occupancy |
| THR | 267 | Side | ASP | 265 | Side | 73,80% | THR | 267 | Side | ASP | 265 | Side | 76,40% |
| THR | 155 | Side | TRP | 151 | Main | 72,80% | TYR | 272 | Side | GLY | 275 | Main | 73,35% |
| TYR | 272 | Side | GLY | 275 | Main | 69,25% | THR | 155 | Side | TRP | 151 | Main | 72,40% |
| TRP | 303 | Main | ILE | 321 | Main | 67,35% | TRP | 303 | Main | ILE | 321 | Main | 67,95% |
| THR | 139 | Side | GLU | 160 | Side | 66,75% | LYS | 306 | Main | LEU | 290 | Main | 63,10% |
| ILE | 156 | Main | ALA | 152 | Main | 63,70% | ARG | 320 | Side | GLU | 315 | Side | 62,70% |
| SER | 218 | Side | GLU | 222 | Main | 63,65% | SER | 218 | Side | GLU | 222 | Main | 60,05% |
| GLU | 283 | Main | ASP | 265 | Side | 61,80% | ILE | 156 | Main | ALA | 152 | Main | 59,80% |
| ASN | 195 | Side | ASP | 185 | Side | 59,85% | ARG | 133 | Side | ASP | 131 | Side | 59,15% |
| LYS | 306 | Main | LEU | 290 | Main | 59,45% | THR | 139 | Side | GLU | 160 | Side | 58,05% |
| ARG | 320 | Side | GLU | 315 | Side | 58,95% | GLU | 283 | Main | ASP | 265 | Side | 57,30% |
| PHE | 197 | Main | MET | 193 | Main | 58,00% | VAL | 217 | Main | GLN | 176 | Side | 55,50% |
| HIP | 287 | Side | ASN | 307 | Side | 57,95% | LYS | 323 | Side | ASP | 296 | Side | 54,90% |
| LEU | 173 | Main | GLU | 160 | Side | 57,65% | HIP | 287 | Side | ASN | 307 | Side | 54,30% |
| ALA | 266 | Main | ASP | 286 | Side | 57,65% | PHE | 197 | Main | MET | 193 | Main | 53,75% |
| SER | 335 | Side | ASP | 194 | Side | 56,80% | TYR | 216 | Side | GLN | 144 | Main | 53,15% |

| | | | | | | | | | | | | | |
|---|---|---|---|---|---|---|---|---|---|---|---|---|---|
| SER | 336 | Side | ALA | 337 | Main | 55,25% | VAL | 241 | Main | SER | 336 | Main | 52,70% |
| VAL | 217 | Main | GLN | 176 | Side | 54,25% | LEU | 173 | Main | GLU | 160 | Side | 52,60% |
| ARG | 133 | Side | ASP | 131 | Side | 54,10% | ALA | 337 | Main | ASN | 158 | Side | 52,10% |
| CYX | 280 | Main | GLN | 327 | Side | 54,05% | ASP | 296 | Main | ALA | 128 | Main | 50,70% |
| ALA | 337 | Main | ASN | 158 | Side | 53,65% | SER | 336 | Side | ALA | 337 | Main | 50,65% |
| GLU | 160 | Main | ILE | 156 | Main | 52,05% | SER | 298 | Side | PRO | 300 | Main | 50,40% |
| LYS | 323 | Side | ASP | 296 | Side | 50,95% | SER | 335 | Side | ASP | 242 | Side | 50,20% |
| VAL | 241 | Main | SER | 336 | Main | 50,70% | GLN | 162 | Main | ASN | 158 | Main | 49,45% |
| GLN | 162 | Main | ASN | 158 | Main | 49,25% | ALA | 266 | Main | ASP | 286 | Side | 49,20% |
| SER | 282 | Side | CYX | 280 | Main | 47,80% | GLU | 160 | Main | ILE | 156 | Main | 46,60% |
| GLU | 322 | Main | LEU | 277 | Main | 47,40% | LYS | 142 | Side | GLU | 175 | Side | 46,30% |
| GLN | 144 | Side | CYX | 147 | Main | 46,90% | GLN | 144 | Side | CYX | 147 | Main | 45,65% |
| ASN | 202 | Main | ASN | 198 | Main | 46,70% | GLU | 322 | Main | LEU | 277 | Main | 45,15% |
| ASP | 296 | Main | ALA | 128 | Main | 46,20% | ASN | 158 | Side | SER | 336 | Side | 44,85% |
| ILE | 250 | Main | ASP | 246 | Main | 45,65% | CYX | 280 | Main | GLN | 327 | Side | 44,10% |
| TYR | 216 | Side | GLN | 144 | Main | 45,10% | GLN | 176 | Side | SER | 218 | Side | 43,90% |
| LYS | 142 | Side | GLU | 175 | Side | 44,65% | ASN | 168 | Side | TRP | 163 | Main | 43,60% |
| SER | 154 | Side | VAL | 289 | Main | 43,90% | SER | 282 | Side | CYX | 280 | Main | 43,25% |
| ASN | 168 | Side | TRP | 163 | Main | 43,15% | SER | 213 | Side | THR | 210 | Side | 43,00% |
| LEU | 291 | Main | LEU | 260 | Main | 42,90% | VAL | 289 | Main | ILE | 262 | Main | 42,20% |
| GLN | 176 | Side | SER | 218 | Side | 42,70% | GLY | 275 | Main | GLU | 315 | Side | 41,55% |
| SER | 172 | Side | GLU | 160 | Side | 42,10% | ILE | 237 | Main | GLY | 206 | Main | 41,55% |
| SER | 298 | Side | PRO | 300 | Main | 41,20% | LEU | 291 | Main | LEU | 260 | Main | 40,65% |
| VAL | 289 | Main | ILE | 262 | Main | 41,15% | SER | 335 | Main | ALA | 261 | Main | 40,60% |
| GLY | 275 | Main | GLU | 315 | Side | 40,75% | ILE | 250 | Main | ASP | 246 | Main | 39,95% |
| GLY | 324 | Main | GLU | 247 | Side | 40,40% | GLY | 324 | Main | GLU | 247 | Side | 38,25% |
| SER | 154 | Main | CYM | 150 | Main | 40,00% | SER | 308 | Side | GLN | 144 | Side | 38,15% |
| ASN | 158 | Side | SER | 336 | Side | 39,55% | SER | 174 | Main | PHE | 209 | Main | 37,85% |
| SER | 213 | Side | THR | 210 | Side | 39,45% | SER | 154 | Side | VAL | 289 | Main | 37,65% |
| ILE | 237 | Main | GLY | 206 | Main | 39,10% | GLN | 162 | Side | PRO | 259 | Main | 37,30% |
| ILE | 305 | Main | ILE | 319 | Main | 36,85% | SER | 336 | Main | VAL | 241 | Main | 36,05% |
| TRP | 303 | Side | GLU | 247 | Side | 35,90% | SER | 174 | Side | TYR | 214 | Main | 35,55% |
| VAL | 264 | Main | HIP | 287 | Main | 35,25% | ILE | 305 | Main | ILE | 319 | Main | 35,30% |
| SER | 174 | Main | PHE | 209 | Main | 35,10% | SER | 154 | Main | CYM | 150 | Main | 34,55% |
| SER | 180 | Main | GLN | 176 | Main | 34,45% | THR | 238 | Main | VAL | 338 | Main | 34,15% |
| ILE | 304 | Main | GLY | 293 | Main | 33,95% | SER | 180 | Main | GLN | 176 | Main | 34,05% |
| LEU | 254 | Main | ILE | 250 | Main | 33,90% | ILE | 304 | Main | GLY | 293 | Main | 33,90% |
| ILE | 200 | Main | ALA | 196 | Main | 33,45% | LYS | 323 | Main | PRO | 301 | Main | 33,45% |
| SER | 180 | Side | PRO | 224 | Main | 33,35% | SER | 172 | Side | GLU | 160 | Side | 33,35% |
| ILE | 184 | Main | ASP | 182 | Side | 33,25% | LYS | 142 | Side | GLU | 160 | Side | 33,15% |
| LYS | 142 | Side | GLU | 160 | Side | 33,05% | TRP | 132 | Main | VAL | 292 | Main | 32,80% |
| TRP | 151 | Side | GLY | 187 | Main | 32,95% | GLU | 211 | Main | SER | 172 | Main | 31,85% |
| SER | 174 | Side | TYR | 214 | Main | 32,75% | VAL | 264 | Main | HIP | 287 | Main | 31,75% |
| LYS | 323 | Main | PRO | 301 | Main | 32,45% | GLN | 176 | Side | SER | 180 | Side | 31,50% |
| GLN | 146 | Main | GLY | 219 | Main | 32,40% | ASN | 331 | Main | ASN | 326 | Side | 30,70% |
| TRP | 199 | Main | ASN | 195 | Main | 31,65% | TRP | 303 | Side | GLU | 247 | Side | 30,55% |
| GLN | 176 | Side | SER | 180 | Side | 31,30% | GLY | 234 | Main | VAL | 208 | Main | 29,65% |

| | | | | | | | | | | | | | |
|---|---|---|---|---|---|---|---|---|---|---|---|---|---|
| THR | 325 | Side | GLU | 322 | Side | 31,05% | ASN | 202 | Main | ASN | 198 | Main | 29,15% |
| CYX | 188 | Main | GLN | 223 | Side | 30,35% | VAL | 292 | Main | ILE | 304 | Main | 29,10% |
| VAL | 292 | Main | ILE | 304 | Main | 29,80% | ASN | 273 | Side | TYR | 272 | Main | 29,10% |
| ASN | 331 | Main | ASN | 326 | Side | 29,75% | LEU | 260 | Main | LEU | 291 | Main | 29,05% |
| SER | 335 | Main | ALA | 261 | Main | 29,65% | GLN | 146 | Main | GLY | 219 | Main | 28,35% |
| THR | 238 | Main | VAL | 338 | Main | 29,35% | ILE | 200 | Main | ALA | 196 | Main | 28,25% |
| ASP | 286 | Main | VAL | 264 | Main | 27,70% | THR | 267 | Main | ASP | 265 | Side | 27,70% |
| VAL | 130 | Main | TYR | 294 | Main | 27,55% | ASN | 204 | Side | ASN | 207 | Main | 27,45% |
| GLU | 211 | Main | SER | 172 | Main | 27,10% | LEU | 254 | Main | ILE | 250 | Main | 26,95% |
| THR | 267 | Main | ASP | 265 | Side | 27,10% | VAL | 130 | Main | TYR | 294 | Main | 26,75% |
| TRP | 132 | Main | VAL | 292 | Main | 26,45% | ASN | 307 | Main | GLY | 317 | Main | 26,15% |
| LYS | 306 | Side | PRO | 140 | Main | 26,20% | GLN | 284 | Main | ASP | 265 | Side | 25,85% |
| GLY | 234 | Main | VAL | 208 | Main | 25,85% | VAL | 338 | Main | ASP | 239 | Main | 25,85% |
| ASN | 257 | Main | TYR | 253 | Main | 25,85% | LEU | 243 | Main | VAL | 334 | Main | 25,25% |
| GLN | 144 | Main | SER | 308 | Main | 25,55% | THR | 210 | Main | GLU | 232 | Main | 25,10% |
| TYR | 294 | Main | VAL | 130 | Main | 25,15% | VAL | 208 | Main | ALA | 235 | Main | 24,80% |
| HIP | 287 | Side | CYM | 150 | Side | 24,60% | ASP | 286 | Main | VAL | 264 | Main | 24,80% |
| GLN | 162 | Side | PRO | 259 | Main | 23,85% | CYX | 188 | Main | GLN | 223 | Side | 24,20% |
| THR | 210 | Main | GLU | 232 | Main | 23,40% | THR | 325 | Side | GLU | 322 | Side | 23,80% |
| LEU | 277 | Main | ARG | 320 | Main | 23,30% | TYR | 294 | Main | VAL | 130 | Main | 23,60% |
| TYR | 253 | Main | ALA | 249 | Main | 22,80% | ASN | 295 | Main | TYR | 302 | Main | 22,45% |
| ASN | 295 | Main | TYR | 302 | Main | 22,35% | GLN | 144 | Main | SER | 308 | Main | 22,45% |
| TRP | 163 | Main | ILE | 159 | Main | 22,30% | LEU | 277 | Main | ARG | 320 | Main | 22,25% |
| ASN | 307 | Main | GLY | 317 | Main | 21,15% | TRP | 163 | Main | ILE | 159 | Main | 21,95% |
| GLN | 284 | Main | ASP | 265 | Side | 20,60% | THR | 155 | Main | TRP | 151 | Main | 21,75% |
| ASN | 204 | Side | ASN | 207 | Main | 20,45% | HIP | 287 | Side | CYM | 150 | Side | 21,50% |
| ASN | 158 | Side | PRO | 259 | Main | 20,45% | GLU | 232 | Main | SER | 213 | Side | 21,35% |
| VAL | 165 | Main | GLY | 161 | Main | 20,40% | VAL | 165 | Main | GLY | 161 | Main | 21,00% |
| GLU | 232 | Main | SER | 213 | Side | 20,35% | HIP | 287 | Main | VAL | 264 | Main | 20,70% |
| THR | 155 | Main | TRP | 151 | Main | 19,75% | ASN | 326 | Main | GLU | 247 | Side | 19,90% |
| ASN | 326 | Main | GLU | 247 | Side | 19,70% | SER | 203 | Main | TRP | 199 | Main | 19,90% |
| ALA | 255 | Main | ALA | 251 | Main | 19,60% | TRP | 199 | Main | ASN | 195 | Main | 19,75% |
| HIP | 287 | Main | VAL | 264 | Main | 19,55% | LYS | 306 | Side | PRO | 140 | Main | 19,45% |
| GLY | 145 | Main | ASP | 143 | Side | 19,50% | ASN | 198 | Side | ASP | 194 | Main | 18,65% |
| VAL | 208 | Main | ALA | 235 | Main | 19,25% | ALA | 255 | Main | ALA | 251 | Main | 18,50% |
| ALA | 252 | Main | ASP | 248 | Main | 17,75% | GLU | 315 | Main | TYR | 318 | Main | 18,40% |
| VAL | 338 | Main | ASP | 239 | Main | 17,70% | ASN | 195 | Side | ASP | 185 | Side | 17,15% |
| GLU | 315 | Main | TYR | 318 | Main | 17,70% | GLY | 221 | Main | SER | 218 | Main | 16,70% |
| ALA | 196 | Main | LEU | 192 | Main | 17,40% | LEU | 290 | Main | LYS | 306 | Main | 16,60% |
| ASP | 185 | Main | ASP | 182 | Side | 16,65% | ALA | 137 | Main | TRP | 132 | Main | 16,45% |
| VAL | 201 | Main | PHE | 197 | Main | 16,60% | GLN | 327 | Side | THR | 325 | Side | 16,35% |
| LEU | 260 | Main | LEU | 291 | Main | 16,50% | GLY | 189 | Main | PHE | 186 | Main | 16,10% |
| ASN | 273 | Side | TYR | 272 | Main | 16,45% | ASN | 158 | Side | PRO | 259 | Main | 15,70% |
| ALA | 137 | Main | TRP | 132 | Main | 14,35% | GLY | 317 | Main | TRP | 313 | Main | 15,70% |
| LEU | 290 | Main | LYS | 306 | Main | 13,95% | SER | 180 | Side | GLN | 176 | Main | 14,90% |
| TYR | 214 | Main | THR | 210 | Main | 13,90% | ASP | 182 | Main | LEU | 178 | Main | 14,80% |
| GLN | 332 | Side | ASN | 331 | Side | 13,90% | TYR | 253 | Main | ALA | 249 | Main | 14,55% |

| | | | | | | | | | | | | | |
|---|---|---|---|---|---|---|---|---|---|---|---|---|---|
| PHE | 269 | Main | ALA | 266 | Main | 13,70% | GLY | 145 | Main | ASP | 143 | Side | 14,55% |
| GLY | 221 | Main | SER | 218 | Main | 13,55% | ALA | 196 | Main | LEU | 192 | Main | 14,45% |
| SER | 310 | Main | LYS | 142 | Main | 13,35% | ALA | 252 | Main | ASP | 248 | Main | 14,40% |
| ARG | 133 | Main | ASP | 131 | Side | 12,90% | ASP | 265 | Main | CYX | 328 | Main | 14,10% |
| SER | 310 | Side | TRP | 309 | Main | 12,85% | ASN | 202 | Side | ASN | 198 | Main | 14,10% |
| ILE | 319 | Main | ILE | 305 | Main | 12,65% | TRP | 151 | Side | GLY | 187 | Main | 14,00% |
| SER | 308 | Side | GLN | 144 | Side | 12,60% | TYR | 346 | Main | GLU | 343 | Main | 13,40% |
| GLY | 157 | Main | PHE | 153 | Main | 12,15% | ARG | 133 | Main | ASP | 131 | Side | 13,05% |
| ASN | 257 | Side | TYR | 253 | Main | 12,05% | TYR | 214 | Main | THR | 210 | Main | 12,85% |
| GLU | 222 | Main | ASN | 220 | Side | 11,85% | GLN | 223 | Side | VAL | 179 | Main | 12,80% |
| GLN | 327 | Side | THR | 325 | Side | 11,75% | ASN | 257 | Main | TYR | 253 | Main | 12,60% |
| SER | 154 | Side | ILE | 262 | Main | 11,55% | SER | 180 | Side | PRO | 224 | Main | 12,50% |
| SER | 172 | Side | GLU | 211 | Side | 10,75% | HIP | 231 | Side | GLN | 227 | Main | 12,50% |
| SER | 203 | Main | TRP | 199 | Main | 10,65% | ASN | 207 | Main | ASN | 204 | Main | 12,45% |
| TYR | 302 | Main | ASN | 295 | Main | 10,60% | TRP | 132 | Side | LEU | 254 | Main | 11,75% |
| ASN | 207 | Main | ASN | 204 | Main | 10,35% | TYR | 302 | Main | ASN | 295 | Main | 11,35% |
| SER | 298 | Side | ASN | 295 | Side | 10,20% | SER | 218 | Main | GLN | 176 | Side | 11,35% |
| LEU | 345 | Main | GLU | 343 | Side | 10,05% | VAL | 179 | Main | GLU | 175 | Main | 11,30% |
| HIP | 231 | Side | GLN | 227 | Main | 9,85% | SER | 154 | Side | ILE | 262 | Main | 11,25% |
| ASP | 248 | Main | ASP | 246 | Side | 9,85% | ASP | 248 | Main | ASP | 246 | Side | 11,15% |
| ASN | 331 | Side | GLU | 247 | Side | 9,85% | THR | 281 | Main | SER | 268 | Side | 11,15% |
| LEU | 192 | Main | ASP | 185 | Side | 9,75% | GLY | 157 | Main | PHE | 153 | Main | 11,15% |
| ALA | 333 | Main | MET | 330 | Main | 9,50% | TRP | 313 | Side | PHE | 269 | Main | 10,75% |
| GLY | 206 | Main | ILE | 200 | Main | 9,45% | SER | 310 | Side | TRP | 309 | Main | 10,75% |
| GLY | 189 | Main | PHE | 186 | Main | 9,45% | ILE | 319 | Main | ILE | 305 | Main | 9,95% |
| SER | 203 | Side | ASN | 202 | Side | 9,45% | ASP | 185 | Main | ASP | 182 | Side | 9,75% |
| SER | 298 | Main | ASN | 295 | Side | 9,40% | ILE | 184 | Main | ASP | 182 | Side | 9,20% |
| GLY | 317 | Main | TRP | 313 | Main | 8,90% | GLU | 222 | Main | ASN | 220 | Side | 9,05% |
| TRP | 132 | Side | LEU | 254 | Main | 8,75% | GLY | 206 | Main | ILE | 200 | Main | 8,85% |
| ASN | 207 | Side | ASN | 204 | Main | 8,70% | SER | 172 | Side | GLU | 211 | Side | 8,70% |
| TYR | 346 | Main | GLU | 343 | Main | 8,65% | ASN | 207 | Side | ASN | 204 | Main | 8,60% |
| ASN | 168 | Side | GLY | 340 | Main | 7,95% | VAL | 201 | Main | PHE | 197 | Main | 8,50% |
| SER | 336 | Main | VAL | 241 | Main | 7,80% | SER | 149 | Side | TYR | 216 | Side | 8,35% |
| LEU | 243 | Main | VAL | 334 | Main | 7,35% | MET | 270 | Main | ALA | 266 | Main | 7,40% |
| TRP | 313 | Side | PHE | 269 | Main | 6,90% | PHE | 269 | Main | ALA | 266 | Main | 7,35% |
| PHE | 347 | Main | GLU | 343 | Main | 6,80% | ALA | 333 | Main | MET | 330 | Main | 7,10% |
| VAL | 179 | Main | GLU | 175 | Main | 6,65% | GLY | 340 | Main | ALA | 236 | Main | 6,75% |
| GLN | 332 | Side | GLN | 245 | Side | 6,55% | HIP | 231 | Side | SER | 213 | Main | 6,40% |
| ASN | 202 | Side | ASN | 198 | Main | 6,35% | TYR | 318 | Main | GLU | 315 | Main | 6,35% |
| GLN | 164 | Side | ASN | 168 | Main | 6,25% | SER | 298 | Main | ASN | 295 | Side | 6,20% |
| TYR | 346 | Side | GLY | 234 | Main | 6,25% | ASN | 295 | Side | TYR | 302 | Side | 6,20% |
| SER | 180 | Side | GLN | 176 | Main | 6,15% | GLY | 167 | Main | GLN | 164 | Main | 6,00% |
| GLN | 245 | Side | GLN | 332 | Side | 6,00% | GLN | 332 | Side | LEU | 329 | Main | 6,00% |
| MET | 330 | Main | GLN | 327 | Main | 5,90% | ASN | 257 | Side | TYR | 253 | Main | 5,75% |
| GLY | 167 | Main | GLN | 164 | Main | 5,85% | ALA | 251 | Main | GLU | 247 | Main | 5,65% |
| GLN | 227 | Side | ASN | 229 | Main | 5,75% | ASN | 198 | Main | ASP | 194 | Main | 5,60% |
| TYR | 318 | Main | GLU | 315 | Main | 5,70% | MET | 330 | Main | GLN | 327 | Main | 5,55% |

| | | | | | | | | | | | | | |
|---|---|---|---|---|---|---|---|---|---|---|---|---|---|
| ASP | 265 | Main | CYX | 328 | Main | 5,55% | PHE | 347 | Main | ASN | 344 | Main | 5,45% |
| GLN | 162 | Side | ASN | 257 | Main | 5,45% | ASN | 229 | Main | GLN | 227 | Side | 5,35% |
| GLN | 223 | Side | VAL | 179 | Main | 5,45% | ASN | 311 | Side | ASP | 316 | Main | 5,35% |
| GLN | 348 | Main | ASN | 344 | Main | 5,30% | ASN | 229 | Side | GLN | 227 | Side | 5,25% |
| SER | 218 | Main | GLN | 176 | Side | 5,30% | SER | 268 | Side | THR | 281 | Main | 5,25% |
| | | | | | | | LEU | 178 | Main | GLU | 175 | Main | 5,20% |
| | | | | | | | GLN | 227 | Side | ASN | 229 | Main | 5,10% |

**Table S5b: Relative occurrence of hydrogen bonds within the catalytic domain.** Two respective MD simulations at pH 8 (pH8_3 and pH8_4, see Table S5a for pH8_1 and pH8_4) were carried out. `Main' refers to interactions of the backbone. 'Side' indicates interactions of side chains. Occurrences below 5% were left out for clarity.

| pH_8_3 | | | | | | | pH_8_4 | | | | | | |
|---|---|---|---|---|---|---|---|---|---|---|---|---|---|
| Found | | | 388 | | | hbonds, | Found | | | 387 | | | hbonds, |
| donor | | | acceptor | | | occupancy | donor | | | acceptor | | | occupancy |
| TYR | 272 | Side | GLY | 275 | Main | 75,56% | TYR | 272 | Side | GLY | 309 | Main | 73,21% |
| THR | 267 | Side | ASP | 265 | Side | 74,11% | THR | 155 | Side | TRP | 192 | Main | 72,36% |
| THR | 155 | Side | TRP | 151 | Main | 71,71% | SER | 335 | Side | ASP | 372 | Side | 72,21% |
| ARG | 320 | Side | GLU | 315 | Side | 69,82% | ALA | 266 | Main | ASP | 303 | Side | 71,86% |
| TRP | 303 | Main | ILE | 321 | Main | 66,57% | SER | 268 | Side | ASP | 305 | Side | 71,01% |
| SER | 218 | Side | GLU | 222 | Main | 63,12% | TRP | 303 | Main | ILE | 340 | Main | 70,56% |
| ILE | 156 | Main | ALA | 152 | Main | 62,22% | LYS | 306 | Main | LEU | 343 | Main | 64,32% |
| GLU | 283 | Main | ASP | 265 | Side | 61,57% | ILE | 156 | Main | ALA | 193 | Main | 64,07% |
| LEU | 173 | Main | GLU | 160 | Side | 58,52% | ARG | 320 | Side | GLU | 357 | Side | 58,02% |
| GLU | 160 | Main | ILE | 156 | Main | 57,07% | SER | 218 | Side | GLU | 255 | Main | 57,17% |
| THR | 139 | Side | GLU | 160 | Side | 56,97% | HIP | 287 | Side | ASN | 324 | Side | 56,12% |
| VAL | 241 | Main | SER | 336 | Main | 53,32% | ARG | 133 | Side | ASP | 170 | Side | 55,37% |
| TYR | 216 | Side | GLN | 144 | Main | 52,72% | ASN | 195 | Side | ASP | 232 | Side | 54,82% |
| ALA | 337 | Main | ASN | 158 | Side | 52,02% | LEU | 173 | Main | GLU | 210 | Side | 54,57% |
| HIP | 287 | Side | ASN | 307 | Side | 52,02% | ASP | 296 | Main | ALA | 333 | Main | 52,77% |
| ALA | 266 | Main | ASP | 286 | Side | 51,57% | PHE | 197 | Main | MET | 234 | Main | 52,57% |
| VAL | 217 | Main | GLN | 176 | Side | 50,87% | LYS | 323 | Side | ASP | 360 | Side | 52,17% |
| LYS | 306 | Main | LEU | 290 | Main | 50,47% | TYR | 216 | Side | GLN | 253 | Main | 50,82% |
| ASP | 296 | Main | ALA | 128 | Main | 50,47% | GLN | 162 | Main | ASN | 199 | Main | 50,32% |
| CYX | 280 | Main | GLN | 327 | Side | 50,22% | SER | 336 | Side | ALA | 373 | Main | 48,23% |
| GLN | 162 | Main | ASN | 158 | Main | 49,68% | VAL | 217 | Main | GLN | 254 | Side | 47,63% |
| LYS | 323 | Side | ASP | 296 | Side | 47,88% | ASN | 168 | Side | TRP | 205 | Main | 47,23% |
| LYS | 142 | Side | GLU | 175 | Side | 47,33% | GLN | 144 | Side | CYX | 181 | Main | 47,08% |
| GLN | 144 | Side | CYX | 147 | Main | 46,68% | ALA | 337 | Main | ASN | 374 | Side | 45,08% |
| GLY | 275 | Main | GLU | 315 | Side | 46,23% | LEU | 291 | Main | LEU | 328 | Main | 44,93% |
| SER | 282 | Side | CYX | 280 | Main | 45,98% | THR | 139 | Side | GLU | 176 | Side | 44,68% |
| SER | 154 | Side | VAL | 289 | Main | 45,48% | LYS | 142 | Side | GLU | 179 | Side | 44,58% |
| GLU | 322 | Main | LEU | 277 | Main | 45,48% | LEU | 277 | Main | ARG | 314 | Main | 44,33% |
| PHE | 197 | Main | MET | 193 | Main | 44,68% | VAL | 241 | Main | SER | 278 | Main | 42,73% |
| SER | 336 | Side | ALA | 337 | Main | 44,28% | VAL | 289 | Main | ILE | 326 | Main | 42,68% |
| LEU | 291 | Main | LEU | 260 | Main | 43,43% | ILE | 237 | Main | GLY | 274 | Main | 42,43% |
| VAL | 289 | Main | ILE | 262 | Main | 43,33% | GLN | 146 | Main | GLY | 183 | Main | 42,38% |

| | | | | | | | | | | | | | |
|---|---|---|---|---|---|---|---|---|---|---|---|---|---|
| ASN | 168 | Side | TRP | 163 | Main | 43,18% | ILE | 250 | Main | ASP | 287 | Main | 41,53% |
| SER | 213 | Side | THR | 210 | Side | 43,13% | SER | 154 | Main | CYM | 191 | Main | 40,73% |
| ILE | 237 | Main | GLY | 206 | Main | 41,88% | SER | 154 | Side | VAL | 191 | Main | 40,58% |
| SER | 180 | Main | GLN | 176 | Main | 41,18% | GLN | 176 | Side | SER | 213 | Side | 40,48% |
| GLN | 176 | Side | SER | 218 | Side | 40,93% | VAL | 264 | Main | HIP | 301 | Main | 40,23% |
| ARG | 133 | Side | ASP | 131 | Side | 39,98% | SER | 298 | Side | PRO | 335 | Main | 39,98% |
| SER | 335 | Main | ALA | 261 | Main | 38,78% | ASN | 158 | Side | SER | 195 | Side | 38,48% |
| ILE | 250 | Main | ASP | 246 | Main | 38,78% | CYX | 188 | Main | GLN | 225 | Side | 38,38% |
| SER | 154 | Main | CYM | 150 | Main | 38,48% | ILE | 305 | Main | ILE | 342 | Main | 38,18% |
| SER | 298 | Side | PRO | 300 | Main | 37,03% | SER | 213 | Side | THR | 250 | Side | 36,78% |
| GLN | 146 | Main | GLY | 219 | Main | 36,88% | SER | 174 | Side | TYR | 211 | Main | 36,58% |
| ASN | 158 | Side | SER | 336 | Side | 36,73% | GLU | 322 | Main | LEU | 359 | Main | 36,33% |
| ASN | 202 | Main | ASN | 198 | Main | 36,63% | LYS | 323 | Main | PRO | 360 | Main | 36,23% |
| SER | 172 | Side | GLU | 160 | Side | 36,58% | ASN | 202 | Main | ASN | 239 | Main | 35,43% |
| LYS | 142 | Side | GLU | 160 | Side | 35,38% | ILE | 304 | Main | GLY | 341 | Main | 35,43% |
| VAL | 264 | Main | HIP | 287 | Main | 34,78% | TRP | 151 | Side | GLY | 188 | Main | 35,38% |
| GLU | 211 | Main | SER | 172 | Main | 34,68% | THR | 238 | Main | VAL | 275 | Main | 34,78% |
| SER | 174 | Side | TYR | 214 | Main | 34,43% | TRP | 132 | Main | VAL | 169 | Main | 34,68% |
| SER | 174 | Main | PHE | 209 | Main | 33,53% | VAL | 292 | Main | ILE | 329 | Main | 34,43% |
| THR | 238 | Main | VAL | 338 | Main | 31,98% | SER | 335 | Main | ALA | 372 | Main | 34,38% |
| ILE | 304 | Main | GLY | 293 | Main | 31,28% | LEU | 254 | Main | ILE | 291 | Main | 34,33% |
| ASN | 331 | Main | ASN | 326 | Side | 31,23% | SER | 174 | Main | PHE | 211 | Main | 34,03% |
| LYS | 323 | Main | PRO | 301 | Main | 30,88% | GLU | 211 | Main | SER | 248 | Main | 33,53% |
| ASP | 286 | Main | VAL | 264 | Main | 30,13% | GLU | 160 | Main | ILE | 197 | Main | 33,53% |
| GLY | 234 | Main | VAL | 208 | Main | 29,99% | ASP | 185 | Main | ASP | 222 | Side | 33,18% |
| THR | 267 | Main | ASP | 265 | Side | 29,79% | LYS | 142 | Side | GLU | 179 | Side | 32,58% |
| GLY | 324 | Main | GLU | 247 | Side | 29,44% | ILE | 200 | Main | ALA | 237 | Main | 32,03% |
| LEU | 254 | Main | ILE | 250 | Main | 29,34% | GLN | 176 | Side | SER | 213 | Side | 31,98% |
| GLN | 284 | Main | ASP | 265 | Side | 29,29% | SER | 180 | Main | GLN | 217 | Main | 31,78% |
| SER | 308 | Side | GLN | 144 | Side | 29,04% | ILE | 184 | Main | ASP | 221 | Side | 31,38% |
| TRP | 303 | Side | GLU | 247 | Side | 28,99% | SER | 180 | Side | PRO | 217 | Main | 30,08% |
| ASN | 257 | Main | TYR | 253 | Main | 28,94% | GLY | 275 | Main | GLU | 312 | Side | 29,89% |
| ILE | 305 | Main | ILE | 319 | Main | 28,64% | TRP | 199 | Main | ASN | 236 | Main | 28,99% |
| GLN | 176 | Side | SER | 180 | Side | 28,34% | GLY | 234 | Main | VAL | 271 | Main | 28,74% |
| VAL | 292 | Main | ILE | 304 | Main | 26,79% | VAL | 130 | Main | TYR | 167 | Main | 27,94% |
| VAL | 130 | Main | TYR | 294 | Main | 26,69% | HIP | 287 | Side | CYM | 324 | Side | 27,09% |
| ASN | 204 | Side | ASN | 207 | Main | 26,69% | THR | 210 | Main | GLU | 247 | Main | 27,04% |
| THR | 325 | Side | GLU | 322 | Side | 26,59% | ASN | 295 | Main | TYR | 332 | Main | 27,04% |
| VAL | 338 | Main | ASP | 239 | Main | 25,59% | VAL | 208 | Main | ALA | 245 | Main | 26,74% |
| GLU | 232 | Main | SER | 213 | Side | 24,64% | ASN | 257 | Main | TYR | 294 | Main | 25,94% |
| LEU | 243 | Main | VAL | 334 | Main | 24,29% | TRP | 303 | Side | GLU | 340 | Side | 25,34% |
| TYR | 294 | Main | VAL | 130 | Main | 23,79% | GLU | 232 | Main | SER | 213 | Side | 25,29% |
| TRP | 163 | Main | ILE | 159 | Main | 23,49% | PHE | 347 | Main | GLU | 384 | Main | 24,89% |
| SER | 335 | Side | ASP | 242 | Side | 23,39% | GLN | 162 | Side | PRO | 199 | Main | 24,54% |
| LEU | 277 | Main | ARG | 320 | Main | 23,19% | CYX | 280 | Main | GLN | 317 | Side | 23,94% |
| VAL | 165 | Main | GLY | 161 | Main | 23,19% | TYR | 294 | Main | VAL | 331 | Main | 23,89% |
| LYS | 306 | Side | PRO | 140 | Main | 22,99% | TYR | 294 | Side | PRO | 331 | Main | 23,79% |

| | | | | | | | | | | | | | |
|---|---|---|---|---|---|---|---|---|---|---|---|---|---|
| ASN | 307 | Main | GLY | 317 | Main | 22,84% | TRP | 163 | Main | ILE | 200 | Main | 22,99% |
| ASN | 295 | Main | TYR | 302 | Main | 22,84% | ASN | 204 | Side | ASN | 241 | Main | 22,74% |
| HIP | 287 | Side | CYM | 150 | Side | 22,84% | VAL | 179 | Main | GLU | 216 | Main | 22,64% |
| ASN | 195 | Side | ASP | 185 | Side | 22,79% | VAL | 165 | Main | GLY | 202 | Main | 22,59% |
| THR | 210 | Main | GLU | 232 | Main | 22,74% | VAL | 338 | Main | ASP | 375 | Main | 22,49% |
| PHE | 269 | Main | ALA | 266 | Main | 22,64% | THR | 281 | Side | ASP | 318 | Side | 22,24% |
| SER | 172 | Side | GLU | 211 | Side | 22,64% | GLN | 144 | Main | SER | 181 | Main | 21,44% |
| GLU | 315 | Main | TYR | 318 | Main | 22,54% | ASN | 331 | Main | ASN | 368 | Side | 21,39% |
| HIP | 287 | Main | VAL | 264 | Main | 22,49% | ALA | 255 | Main | ALA | 292 | Main | 20,79% |
| GLN | 223 | Side | VAL | 179 | Main | 22,49% | LEU | 260 | Main | LEU | 297 | Main | 20,69% |
| TRP | 132 | Main | VAL | 292 | Main | 22,19% | ALA | 137 | Main | TRP | 174 | Main | 19,44% |
| ILE | 200 | Main | ALA | 196 | Main | 21,79% | ASN | 158 | Side | PRO | 195 | Main | 19,09% |
| VAL | 208 | Main | ALA | 235 | Main | 21,29% | HIP | 231 | Side | GLN | 268 | Main | 19,04% |
| GLN | 144 | Main | SER | 308 | Main | 21,14% | GLY | 145 | Main | ASP | 182 | Side | 18,89% |
| GLN | 162 | Side | PRO | 259 | Main | 20,94% | GLY | 324 | Main | GLU | 361 | Side | 18,79% |
| SER | 335 | Side | ASP | 194 | Side | 20,79% | LYS | 306 | Side | PRO | 343 | Main | 17,99% |
| ASN | 158 | Side | PRO | 259 | Main | 19,99% | GLU | 315 | Main | TYR | 352 | Main | 16,84% |
| SER | 336 | Main | VAL | 241 | Main | 19,54% | ASN | 326 | Main | GLU | 363 | Side | 16,74% |
| TYR | 253 | Main | ALA | 249 | Main | 19,49% | ASN | 307 | Main | GLY | 344 | Main | 16,74% |
| TRP | 199 | Main | ASN | 195 | Main | 19,09% | TYR | 253 | Main | ALA | 290 | Main | 16,29% |
| THR | 155 | Main | TRP | 151 | Main | 18,64% | VAL | 201 | Main | PHE | 238 | Main | 16,29% |
| GLN | 327 | Side | THR | 325 | Side | 18,34% | ASN | 198 | Side | ASP | 235 | Main | 15,84% |
| PHE | 347 | Main | GLU | 343 | Main | 18,24% | THR | 281 | Main | SER | 318 | Side | 15,59% |
| GLY | 145 | Main | ASP | 143 | Side | 18,14% | ASP | 182 | Main | LEU | 219 | Main | 15,54% |
| LEU | 260 | Main | LEU | 291 | Main | 17,79% | ALA | 252 | Main | ASP | 289 | Main | 15,14% |
| SER | 180 | Side | PRO | 224 | Main | 17,39% | ASP | 286 | Main | VAL | 323 | Main | 14,99% |
| ASN | 326 | Main | GLU | 247 | Side | 16,84% | THR | 155 | Main | TRP | 192 | Main | 14,94% |
| ILE | 184 | Main | ASP | 182 | Side | 16,69% | ARG | 133 | Main | ASP | 170 | Side | 14,54% |
| GLY | 221 | Main | SER | 218 | Main | 16,64% | ALA | 196 | Main | LEU | 233 | Main | 14,39% |
| TRP | 132 | Side | LEU | 254 | Main | 16,39% | LEU | 290 | Main | LYS | 327 | Main | 14,09% |
| LYS | 135 | Main | ASP | 131 | Main | 16,39% | SER | 203 | Main | TRP | 240 | Main | 13,99% |
| ALA | 255 | Main | ALA | 251 | Main | 16,29% | ASN | 331 | Side | GLN | 368 | Main | 13,99% |
| ALA | 252 | Main | ASP | 248 | Main | 15,39% | HIP | 287 | Main | VAL | 324 | Main | 13,49% |
| GLN | 332 | Side | ASN | 331 | Side | 15,19% | GLY | 221 | Main | SER | 258 | Main | 13,44% |
| ILE | 319 | Main | ILE | 305 | Main | 15,09% | ASN | 257 | Side | TYR | 294 | Main | 13,39% |
| ASP | 185 | Main | ASP | 182 | Side | 14,64% | SER | 172 | Side | GLU | 209 | Side | 12,89% |
| LEU | 290 | Main | LYS | 306 | Main | 13,54% | ASN | 202 | Side | ASN | 239 | Main | 12,84% |
| ASN | 273 | Side | TYR | 272 | Main | 13,49% | TYR | 346 | Main | GLU | 383 | Main | 12,69% |
| HIP | 231 | Side | GLN | 227 | Main | 13,24% | SER | 308 | Side | GLN | 345 | Side | 12,09% |
| GLU | 222 | Main | ASN | 220 | Side | 13,19% | SER | 268 | Main | ASP | 305 | Side | 12,09% |
| SER | 203 | Main | TRP | 199 | Main | 13,04% | ALA | 251 | Main | GLU | 288 | Main | 11,99% |
| VAL | 201 | Main | PHE | 197 | Main | 12,99% | TYR | 302 | Main | ASN | 339 | Main | 11,84% |
| LEU | 345 | Main | GLU | 343 | Side | 12,49% | GLU | 222 | Main | ASN | 259 | Side | 11,79% |
| ASN | 202 | Side | ASN | 198 | Main | 12,39% | GLY | 206 | Main | ILE | 243 | Main | 10,94% |
| ASN | 207 | Main | ASN | 204 | Main | 11,79% | ASP | 248 | Main | ASP | 285 | Side | 10,89% |
| ALA | 137 | Main | TRP | 132 | Main | 11,59% | ASN | 344 | Side | GLU | 381 | Side | 10,84% |
| ALA | 196 | Main | LEU | 192 | Main | 11,54% | HIP | 231 | Side | SER | 268 | Main | 10,69% |

| | | | | | | | | | | | | | |
|---|---|---|---|---|---|---|---|---|---|---|---|---|---|
| SER | 218 | Main | GLN | 176 | Side | 11,44% | SER | 180 | Side | GLN | 217 | Main | 10,69% |
| CYX | 188 | Main | GLN | 223 | Side | 11,39% | SER | 310 | Side | TRP | 347 | Main | 10,14% |
| TYR | 302 | Main | ASN | 295 | Main | 11,34% | TRP | 132 | Side | LEU | 169 | Main | 10,09% |
| GLN | 162 | Side | ASN | 257 | Main | 11,19% | SER | 310 | Main | LYS | 347 | Main | 9,95% |
| ASP | 248 | Main | ASP | 246 | Side | 11,14% | MET | 330 | Main | GLN | 367 | Main | 9,90% |
| VAL | 179 | Main | GLU | 175 | Main | 10,54% | SER | 298 | Side | ASN | 335 | Side | 9,30% |
| ASN | 257 | Side | TYR | 253 | Main | 10,14% | ILE | 319 | Main | ILE | 356 | Main | 9,25% |
| TYR | 214 | Main | THR | 210 | Main | 10,04% | GLN | 162 | Side | ASN | 199 | Main | 9,10% |
| GLN | 227 | Side | ASN | 229 | Main | 9,95% | GLN | 332 | Side | ASN | 369 | Side | 8,95% |
| ALA | 251 | Main | GLU | 247 | Main | 9,95% | LEU | 192 | Main | ASP | 229 | Side | 8,80% |
| SER | 180 | Side | GLN | 176 | Main | 9,90% | TRP | 199 | Side | THR | 236 | Side | 8,60% |
| GLY | 157 | Main | PHE | 153 | Main | 9,65% | TYR | 346 | Main | PRO | 383 | Main | 8,55% |
| GLY | 317 | Main | TRP | 313 | Main | 9,65% | SER | 154 | Side | ILE | 191 | Main | 8,50% |
| TYR | 346 | Main | GLU | 343 | Main | 9,30% | ASN | 207 | Main | ASN | 244 | Main | 8,30% |
| GLY | 206 | Main | ILE | 200 | Main | 9,05% | GLY | 157 | Main | PHE | 194 | Main | 8,25% |
| SER | 149 | Side | TYR | 216 | Side | 8,95% | GLN | 332 | Side | LEU | 369 | Main | 7,95% |
| GLN | 164 | Side | ALA | 137 | Main | 8,45% | GLY | 317 | Main | TRP | 354 | Main | 7,85% |
| MET | 330 | Main | GLN | 327 | Main | 8,40% | GLY | 189 | Main | PHE | 226 | Main | 7,70% |
| SER | 154 | Side | ILE | 262 | Main | 8,35% | ASN | 207 | Side | ASN | 244 | Main | 7,70% |
| GLY | 189 | Main | PHE | 186 | Main | 8,25% | LYS | 306 | Side | THR | 343 | Main | 7,45% |
| SER | 310 | Main | LYS | 142 | Main | 8,10% | TYR | 214 | Main | THR | 251 | Main | 7,45% |
| ALA | 261 | Main | SER | 335 | Main | 8,00% | ASN | 273 | Side | TYR | 310 | Main | 7,25% |
| SER | 298 | Main | ASN | 295 | Side | 7,85% | LEU | 243 | Main | VAL | 280 | Main | 7,05% |
| ARG | 133 | Main | ASP | 131 | Side | 7,45% | LEU | 178 | Main | GLU | 215 | Main | 6,95% |
| ASN | 207 | Side | ASN | 204 | Main | 7,40% | TYR | 318 | Main | GLU | 355 | Main | 6,65% |
| ASN | 198 | Side | ASP | 194 | Main | 7,30% | SER | 336 | Main | VAL | 373 | Main | 6,55% |
| TRP | 151 | Side | GLY | 187 | Main | 7,20% | ALA | 333 | Main | MET | 370 | Main | 6,40% |
| HIP | 231 | Side | SER | 213 | Main | 7,10% | ILE | 262 | Main | VAL | 299 | Main | 6,25% |
| ASP | 265 | Main | CYX | 328 | Main | 6,95% | ALA | 261 | Main | SER | 298 | Main | 6,15% |
| GLN | 164 | Main | GLU | 160 | Main | 6,95% | SER | 298 | Main | ASN | 335 | Side | 6,10% |
| ASN | 295 | Side | TYR | 302 | Side | 6,20% | SER | 218 | Main | GLN | 255 | Side | 5,90% |
| GLN | 245 | Side | GLN | 332 | Side | 6,15% | GLY | 167 | Main | GLN | 204 | Main | 5,90% |
| ASN | 168 | Side | GLY | 340 | Main | 5,70% | ASN | 295 | Side | TYR | 332 | Side | 5,85% |
| TRP | 313 | Side | PHE | 269 | Main | 5,45% | LEU | 345 | Main | PRO | 382 | Main | 5,80% |
| TYR | 318 | Main | GLU | 315 | Main | 5,35% | SER | 218 | Side | VAL | 255 | Main | 5,70% |
| GLN | 164 | Side | ASN | 168 | Main | 5,15% | GLN | 332 | Main | LEU | 369 | Main | 5,40% |
| LEU | 192 | Main | ASP | 185 | Side | 5,05% | ILE | 321 | Main | TRP | 358 | Main | 5,25% |
| ILE | 262 | Main | VAL | 289 | Main | 5,05% | | | | | | | |
| ALA | 166 | Main | GLN | 162 | Main | 5,05% | | | | | | | |
| GLY | 340 | Main | ALA | 236 | Main | 5,05% | | | | | | | |
| GLU | 134 | Main | ASP | 131 | Side | 5,05% | | | | | | | |

**Table S6a: Relative occurrence of hydrogen bonds between pro- and catalytic domain.** Four respective MD simulations at pH 4 (pH4_1 and pH4_2, see Table S6b for pH4_3 and pH4_4) were carried out. `Main' refers to interactions of the backbone. 'Side' indicates interactions of side chains. Occurrences below 5% were left out for clarity.

| pH4_1 | | | | | | | pH 4_2 | | | | | | |
|---|---|---|---|---|---|---|---|---|---|---|---|---|---|
| donor | | | acceptor | | | occupancy | donor | | | acceptor | | | occupancy |
| TYR | 101 | Side | CYM | 150 | Side | 64,50% | THR | 83 | Main | GLY | 274 | Main | 43,50% |
| ASH | 271 | Side | GLN | 71 | Side | 63,10% | ASH | 271 | Side | ASN | 68 | Side | 41,40% |
| SER | 268 | Side | GLN | 75 | Side | 57,50% | THR | 87 | Side | MET | 270 | Main | 39,25% |
| SER | 90 | Side | MET | 270 | Main | 46,15% | GLY | 85 | Main | ASH | 271 | Main | 38,65% |
| VAL | 119 | Main | ASP | 242 | Main | 44,55% | SER | 279 | Main | ASN | 79 | Side | 36,25% |
| ASN | 79 | Side | SER | 279 | Main | 38,85% | VAL | 119 | Main | ASP | 242 | Main | 33,60% |
| THR | 83 | Main | GLY | 274 | Main | 36,10% | SER | 90 | Side | ASH | 271 | Side | 32,15% |
| GLY | 85 | Main | ASH | 271 | Main | 34,95% | ILE | 276 | Main | TYR | 81 | Main | 25,70% |
| SER | 279 | Main | ASN | 79 | Side | 26,20% | ARG | 102 | Side | ASP | 265 | Side | 20,35% |
| ASP | 242 | Main | LYS | 117 | Main | 26,20% | TYR | 107 | Side | ILE | 184 | Main | 19,85% |
| GLN | 71 | Side | THR | 267 | Main | 22,75% | ASP | 242 | Main | LYS | 117 | Main | 18,00% |
| LYS | 117 | Main | ASP | 242 | Side | 21,15% | ASN | 273 | Side | VAL | 86 | Main | 17,80% |
| ASN | 273 | Side | VAL | 86 | Main | 19,05% | ARG | 100 | Side | GLN | 146 | Side | 17,15% |
| GLN | 75 | Side | SER | 268 | Main | 18,30% | LYS | 117 | Main | ASP | 242 | Side | 15,85% |
| GLN | 75 | Side | ASH | 271 | Side | 17,15% | ASN | 273 | Main | THR | 83 | Main | 12,10% |
| ARG | 114 | Side | ASN | 195 | Side | 16,85% | LYS | 117 | Side | HIP | 240 | Main | 8,85% |
| TYR | 107 | Side | ILE | 184 | Main | 16,55% | ARG | 116 | Side | ASP | 242 | Side | 8,00% |
| THR | 278 | Main | ASN | 79 | Side | 16,10% | THR | 278 | Main | ASN | 79 | Side | 7,85% |
| LYS | 117 | Side | ASP | 239 | Side | 13,30% | LYS | 117 | Side | ASP | 239 | Side | 6,75% |
| GLN | 245 | Side | GLN | 112 | Side | 12,80% | TYR | 107 | Side | GLY | 189 | Main | 6,50% |
| THR | 281 | Side | GLN | 75 | Side | 12,60% | ASN | 79 | Side | SER | 279 | Main | 6,35% |
| ASN | 120 | Main | TYR | 346 | Main | 12,25% | ARG | 116 | Side | VAL | 334 | Main | 5,95% |
| ASN | 273 | Main | THR | 83 | Main | 12,20% | GLN | 245 | Side | GLN | 112 | Side | 5,30% |
| THR | 281 | Main | GLN | 75 | Side | 11,90% | GLN | 112 | Side | GLN | 332 | Side | 5,15% |
| ILE | 276 | Main | TYR | 81 | Main | 11,65% | | | | | | | |
| GLN | 284 | Side | ALA | 105 | Main | 9,15% | | | | | | | |
| ARG | 116 | Side | VAL | 334 | Main | 7,25% | | | | | | | |
| THR | 281 | Side | GLN | 71 | Side | 6,85% | | | | | | | |
| GLN | 112 | Side | GLN | 245 | Side | 6,35% | | | | | | | |
| ARG | 98 | Side | GLH | 283 | Side | 5,85% | | | | | | | |
| GLY | 104 | Main | GLY | 189 | Main | 5,70% | | | | | | | |
| GLY | 124 | Main | ALA | 252 | Main | 5,00% | | | | | | | |

**Table S6b: Relative occurrence of hydrogen bonds between pro- and catalytic domain.** Four respective MD simulations at pH 4 (pH4_3 and pH4_4, see Table S6a for pH4_1 and pH4_2) were carried out. `Main' refers to interactions of the backbone. 'Side' indicates interactions of side chains. Occurrences below 5% were left out for clarity.

| pH 4_3 | | | | | | | pH 4_4 | | | | | | |
|---|---|---|---|---|---|---|---|---|---|---|---|---|---|
| donor | | | acceptor | | | occupancy | donor | | | acceptor | | | occupancy |
| THR | 87 | Side | MET | 270 | Main | 57,47% | SER | 279 | Main | ASN | 79 | Side | 38,43% |
| THR | 83 | Main | GLY | 274 | Main | 41,98% | GLY | 85 | Main | ASH | 271 | Main | 36,78% |
| GLY | 85 | Main | ASH | 271 | Main | 38,63% | THR | 83 | Main | GLY | 274 | Main | 35,83% |
| SER | 279 | Main | ASN | 79 | Side | 34,98% | VAL | 119 | Main | ASP | 242 | Main | 32,53% |
| VAL | 119 | Main | ASP | 242 | Main | 32,23% | ASN | 273 | Side | VAL | 86 | Main | 28,44% |
| ASN | 79 | Side | SER | 279 | Main | 23,34% | ILE | 276 | Main | TYR | 81 | Main | 17,24% |
| ASH | 271 | Side | ASN | 68 | Side | 21,94% | ASN | 273 | Main | THR | 83 | Main | 16,94% |
| THR | 122 | Side | GLN | 348 | Main | 21,74% | THR | 87 | Side | MET | 270 | Main | 15,39% |
| ASN | 273 | Side | VAL | 86 | Main | 20,94% | ASN | 79 | Side | SER | 279 | Main | 12,94% |
| TYR | 107 | Side | ILE | 184 | Main | 20,94% | SER | 268 | Side | GLN | 75 | Side | 12,04% |
| ASN | 273 | Main | THR | 83 | Main | 14,39% | GLN | 112 | Side | GLN | 332 | Side | 10,54% |
| SER | 268 | Side | GLN | 75 | Side | 13,94% | THR | 278 | Main | ASN | 79 | Side | 8,65% |
| THR | 278 | Main | ASN | 79 | Side | 13,84% | GLN | 75 | Side | THR | 281 | Side | 8,65% |
| ILE | 276 | Main | TYR | 81 | Main | 13,49% | TYR | 107 | Side | ILE | 184 | Main | 7,80% |
| SER | 90 | Side | ASH | 271 | Side | 13,29% | TYR | 107 | Side | GLY | 189 | Main | 6,05% |
| ASN | 103 | Side | GLY | 189 | Main | 11,74% | ARG | 116 | Side | VAL | 334 | Main | 5,55% |
| GLN | 112 | Side | GLN | 332 | Side | 9,60% | ARG | 100 | Side | GLN | 146 | Side | 5,50% |
| THR | 122 | Side | GLN | 348 | Side | 9,60% | TYR | 101 | Side | ALA | 266 | Main | 5,00% |
| ARG | 100 | Side | GLN | 146 | Side | 9,30% | | | | | | | |
| ARG | 116 | Side | ASP | 242 | Side | 9,00% | | | | | | | |
| THR | 122 | Main | GLN | 348 | Side | 6,50% | | | | | | | |
| TYR | 48 | Side | MET | 312 | Main | 6,15% | | | | | | | |
| GLN | 75 | Side | THR | 281 | Side | 6,15% | | | | | | | |
| ASH | 271 | Side | GLN | 71 | Side | 5,40% | | | | | | | |

**Table S7a: Relative occurrence of hydrogen bonds between pro- and catalytic domain.** Four respective MD simulations at pH 8 (pH8_1 and pH8_2, see Table 7b for pH8_3 and pH8_4) were carried out. `Main' refers to interactions of the backbone. 'Side' indicates interactions of side chains. Occurrences below 5% were left out for clarity.

| pH_8_1 | | | | | | | pH 8_2 | | | | | | |
|---|---|---|---|---|---|---|---|---|---|---|---|---|---|
| donor | | | acceptor | | | occupancy | donor | | | acceptor | | | occupancy |
| GLN | 71 | Side | ASP | 271 | Side | 66,50% | GLN | 75 | Side | ASP | 271 | Side | 60,30% |
| THR | 87 | Side | MET | 270 | Main | 43,95% | ARG | 116 | Side | ASP | 242 | Side | 52,35% |
| THR | 83 | Main | GLY | 274 | Main | 39,15% | THR | 83 | Main | GLY | 274 | Main | 41,35% |
| SER | 279 | Main | ASN | 79 | Side | 36,40% | VAL | 119 | Main | ASP | 242 | Main | 41,10% |
| VAL | 119 | Main | ASP | 242 | Main | 34,05% | ASP | 242 | Main | LYS | 117 | Main | 37,95% |
| GLY | 85 | Main | ASP | 271 | Main | 33,05% | THR | 87 | Side | MET | 270 | Main | 37,65% |
| ARG | 116 | Side | VAL | 334 | Main | 29,15% | LYS | 117 | Main | ASP | 242 | Side | 34,70% |
| ASN | 273 | Side | VAL | 86 | Main | 27,30% | SER | 279 | Main | ASN | 79 | Side | 27,05% |

| | | | | | | | | | | | | | |
|---|---|---|---|---|---|---|---|---|---|---|---|---|---|
| ASN | 68 | Side | ASP | 271 | Side | 25,45% | TYR | 107 | Side | ASP | 185 | Side | 25,60% |
| GLN | 75 | Side | ASP | 271 | Side | 24,20% | ILE | 276 | Main | TYR | 81 | Main | 24,30% |
| ILE | 276 | Main | TYR | 81 | Main | 23,70% | TYR | 48 | Side | MET | 312 | Main | 23,90% |
| ARG | 114 | Side | ASP | 242 | Side | 21,85% | GLY | 85 | Main | ASP | 271 | Main | 23,80% |
| TYR | 107 | Side | ASP | 185 | Side | 21,05% | ARG | 116 | Side | ASP | 194 | Side | 22,05% |
| ARG | 116 | Side | ASP | 194 | Side | 19,60% | ARG | 98 | Side | THR | 267 | Side | 20,35% |
| THR | 281 | Side | GLN | 75 | Side | 18,35% | ASN | 273 | Side | VAL | 86 | Main | 17,85% |
| ASN | 273 | Main | THR | 83 | Main | 11,80% | ARG | 114 | Side | ASN | 195 | Side | 17,55% |
| ASN | 103 | Side | GLY | 189 | Main | 11,20% | ASN | 103 | Side | GLY | 189 | Main | 16,45% |
| ARG | 98 | Side | THR | 267 | Side | 9,75% | ARG | 114 | Side | ASP | 194 | Side | 15,05% |
| ARG | 114 | Side | ASP | 194 | Side | 9,05% | LYS | 117 | Side | HIE | 240 | Main | 14,35% |
| LYS | 117 | Main | ASP | 242 | Side | 7,25% | THR | 281 | Side | GLN | 75 | Side | 14,30% |
| GLN | 112 | Side | GLN | 332 | Main | 6,70% | ARG | 98 | Side | ASP | 286 | Side | 13,00% |
| ASN | 79 | Side | SER | 279 | Main | 5,05% | ASN | 273 | Main | THR | 83 | Main | 9,20% |
| GLN | 112 | Side | GLN | 332 | Side | 5,00% | GLN | 71 | Side | ASP | 271 | Side | 8,90% |
| | | | | | | | LYS | 117 | Side | ASP | 239 | Side | 8,65% |
| | | | | | | | GLN | 75 | Side | SER | 268 | Main | 7,60% |
| | | | | | | | ASN | 79 | Side | SER | 279 | Main | 7,50% |
| | | | | | | | ARG | 100 | Side | GLN | 146 | Side | 7,15% |
| | | | | | | | ARG | 125 | Side | GLU | 256 | Side | 6,55% |
| | | | | | | | SER | 268 | Side | GLN | 75 | Side | 5,70% |
| | | | | | | | GLN | 75 | Side | SER | 268 | Side | 5,40% |
| | | | | | | | GLN | 112 | Side | GLN | 332 | Side | 5,00% |

**Table S7b: Relative occurrence of hydrogen bonds between pro- and catalytic domain.** Four respective MD simulations at pH 8 (pH8_3 and pH8_4, see Table S7a for pH8_1 and pH8_2) were carried out. \`Main' refers to interactions of the backbone. 'Side' indicates interactions of side chains. Occurrences below 5% were left out for clarity.

| pH 8_3 | | | | | | | pH 8_4 | | | | | | |
|---|---|---|---|---|---|---|---|---|---|---|---|---|---|
| donor | | | acceptor | | | occupancy | donor | | | acceptor | | | occupancy |
| ASN | 68 | Side | ASP | 271 | Side | 70,51% | THR | 87 | Side | MET | 270 | Main | 64,42% |
| GLN | 75 | Side | ASP | 271 | Side | 51,67% | ARG | 116 | Side | ASP | 194 | Side | 50,07% |
| ARG | 116 | Side | ASP | 194 | Side | 49,58% | THR | 83 | Main | GLY | 274 | Main | 41,83% |
| THR | 87 | Side | MET | 270 | Main | 48,03% | ASN | 68 | Side | ASP | 271 | Side | 39,38% |
| VAL | 119 | Main | ASP | 242 | Main | 42,78% | ARG | 116 | Side | VAL | 334 | Main | 36,68% |
| ARG | 125 | Side | GLU | 256 | Side | 37,68% | GLN | 75 | Side | ASP | 271 | Side | 27,89% |
| THR | 83 | Main | GLY | 274 | Main | 37,58% | THR | 83 | Side | ASN | 273 | Side | 27,69% |
| SER | 279 | Main | ASN | 79 | Side | 33,23% | ASN | 273 | Main | THR | 83 | Main | 25,74% |
| ARG | 94 | Side | ASP | 271 | Side | 30,38% | ILE | 276 | Main | TYR | 81 | Main | 25,14% |
| TYR | 107 | Side | ASP | 185 | Side | 26,34% | ARG | 94 | Side | ASP | 271 | Side | 24,94% |
| LYS | 117 | Main | ASP | 242 | Side | 24,69% | GLY | 85 | Main | ASP | 271 | Main | 24,84% |
| ASN | 273 | Side | VAL | 86 | Main | 19,49% | THR | 122 | Side | GLU | 256 | Side | 23,99% |
| GLY | 85 | Main | ASP | 271 | Main | 17,59% | LYS | 117 | Main | ASP | 242 | Side | 19,94% |
| GLN | 71 | Side | ASP | 271 | Side | 16,49% | GLN | 71 | Side | ASP | 271 | Side | 19,54% |
| ILE | 276 | Main | TYR | 81 | Main | 16,19% | LYS | 117 | Side | ASP | 242 | Side | 17,04% |
| ASN | 103 | Side | GLY | 189 | Main | 15,29% | SER | 279 | Main | ASN | 79 | Side | 14,04% |

| | | | | | | | | | | | | | |
|---|---|---|---|---|---|---|---|---|---|---|---|---|---|
| ASP | 242 | Main | LYS | 117 | Main | 10,84% | ARG | 102 | Side | GLU | 283 | Side | 13,04% |
| ARG | 102 | Side | ASP | 286 | Side | 9,65% | GLN | 75 | Side | SER | 268 | Main | 11,49% |
| ARG | 116 | Side | VAL | 334 | Main | 9,45% | THR | 122 | Main | ALA | 252 | Main | 11,44% |
| ARG | 114 | Side | ASN | 195 | Side | 8,35% | ARG | 114 | Side | ASN | 195 | Side | 10,59% |
| ASN | 273 | Main | THR | 83 | Main | 7,70% | TYR | 107 | Side | GLY | 189 | Main | 8,75% |
| ASN | 79 | Side | SER | 279 | Main | 7,10% | GLN | 112 | Side | GLN | 332 | Main | 7,75% |
| ARG | 102 | Side | GLN | 284 | Side | 6,60% | ASN | 273 | Side | VAL | 86 | Main | 7,60% |
| GLN | 75 | Side | SER | 268 | Main | 6,45% | ARG | 114 | Side | ASP | 242 | Side | 7,60% |
| | | | | | | | TYR | 48 | Side | MET | 312 | Main | 7,10% |
| | | | | | | | ARG | 125 | Side | GLU | 256 | Side | 6,25% |
| | | | | | | | ASN | 103 | Side | GLY | 189 | Main | 5,35% |
| | | | | | | | LYS | 117 | Main | ASP | 242 | Main | 5,30% |

### *References*

1. Caffrey, C. R. *et al.* Active site mapping, biochemical properties and subcellular localization of rhodesain, the major cysteine protease of Trypanosoma brucei rhodesiense. *Mol. Biochem. Parasitol.* **118**, 61–73 (2001).
2. Kerr, I. D. *et al.* Vinyl sulfones as antiparasitic agents and a structural basis for drug design. *J. Biol. Chem.* **284**, 25697–25703 (2009).
3. Micsonai, A. *et al.* Accurate secondary structure prediction and fold recognition for circular dichroism spectroscopy. 1–9 (2015).
